# Genomic properties of variably methylated retrotransposons in mouse

**DOI:** 10.1101/2020.10.21.349217

**Authors:** Jessica L. Elmer, Amir D. Hay, Noah J. Kessler, Tessa M. Bertozzi, Eve Ainscough, Anne C. Ferguson-Smith

## Abstract

Transposable elements (TEs) are enriched in cytosine methylation, preventing their mobility within the genome. We previously identified a genome-wide repertoire of candidate intracisternal A particle (IAP) TEs in mice that exhibit inter-individual variability in this methylation (VM-IAPs) with implications for genome function. Here we validate these metastable epialleles and discover a novel class that exhibit tissue specificity (tsVM-IAPs) in addition to those with uniform methylation in all tissues (constitutive- or cVM-IAPs); both types have the potential to regulate genes in *cis*. Screening for variable methylation at other TEs shows that this phenomenon is largely limited to IAPs, which are amongst the youngest and most active endogenous retroviruses. We identify sequences enriched within cVM-IAPs, but determine that these are not sufficient to confer epigenetic variability. CTCF is enriched at VM-IAPs with binding inversely correlated with DNA methylation. We uncover dynamic physical interactions between cVM-IAPs with low methylation ranges and other genomic loci, suggesting that VM-IAPs have the potential for long-range regulation. Our findings indicate that a recently evolved interplay between genetic sequence, CTCF binding, and DNA methylation at young TEs can result in inter-individual variability in transcriptional outcomes with implications for phenotypic variation.

## Introduction

Transposable elements (TEs) are DNA sequences that account for about 40% of the mouse genome (Mouse Genome Sequencing et al. 2002). The vast majority (96%) of TEs are retrotransposons, which mobilise via an RNA intermediate prior to re-integration into the genome (Mouse Genome Sequencing et al. 2002; Nellaker et al. 2012). There are three classes of retrotransposons in mammals: long-interspersed nuclear elements (LINEs), short-interspersed nuclear elements (SINEs), and long-terminal repeat elements (LTRs; which include endogenous retroviruses (ERVs)). Whilst ERVs mostly exist as solo LTRs, and the majority of full-length elements have lost coding potential due to an accumulation of mutations, some mouse ERVs retain the ability to retrotranspose and account for up to 12% of all germline mutations (Maksakova et al. 2006). Due to the risk of insertional mutation and the potential activity of internal regulatory sequences, retrotransposons are commonly targeted for silencing by ncRNAs, repressive histone modifications, and DNA methylation (Bourque et al. 2018).

Of all the types of ERVs in the mouse genome, intracisternal A-particle elements (IAPs) are amongst the most active (Heidmann and Heidmann 1991; Dewannieux et al. 2004) and evolutionarily young (Maksakova et al. 2006; Zhang et al. 2008; Nellaker et al. 2012; Gagnier et al. 2019). Unlike 90% of the genome, IAPs have been reported to resist the epigenetic reprogramming that occurs during early embryonic development and remain methylated (Lane et al. 2003; Seisenberger et al. 2013). It is therefore rare for IAPs to escape methylation. *Agouti viable yellow* (*A*^*vy*^) and *Axin Fused* (*Axin*^*Fu*^) represent two-well studied examples of IAPs that are not fully methylated. These alleles were first identified due to observable differences between littermates in coat colour and tail morphology, respectively (Reed 1937; Dickies 1962) and were termed ‘metastable epialleles’ (Rakyan et al. 2002; Bertozzi and Ferguson-Smith 2020). It was determined that these phenotypic differences result from an IAP insertion and that the methylation level of the element is inversely correlated with the expression of the affected gene (Vasicek et al. 1997; Morgan et al. 1999). These two IAP insertions have inter-individual variation in DNA methylation that gives rise to individual mice exhibiting coat colours ranging from yellow to pseudoagouti (*A*^*vy*^), and tail morphology ranging from highly kinked to straight (*Axin*^*Fu*^). Furthermore, both models transmit a memory of the parental methylation state to the offspring, providing a paradigm for transgenerational epigenetic inheritance (Morgan et al. 1999; Rakyan et al. 2003).

Previously, our group performed a systematic genome-wide screen in C57BL/6J mice to identify other variably methylated IAPs (VM-IAPs) (Kazachenka et al. 2018). Unlike what was observed at the *A*^*vy*^ and *Axin*^*Fu*^ loci, the majority of VM-IAPs identified in that screen did not have any detectable effect on the expression of adjacent genes. In offspring, VM-IAPs do not possess a memory of the parental methylation state; instead, the DNA methylation at VM-IAPs is predictably reprogrammed following fertilization and re-established stochastically in the next generation (Bertozzi and Ferguson-Smith 2020). The functional implications of DNA methylation at these elements and the mechanisms behind the establishment of variable methylation are not fully understood.

In this study, we validate candidate VM-IAPs that show consistent levels of methylation between tested tissues, which we term constitutive variably methylated IAPs (cVM-IAPs). In addition, we identify and characterise IAP elements that only have variable methylation in some, but not all, tested tissues and term these tissue-specific variably methylated IAPs (tsVM-IAPs). We find that cVM-IAPs are enriched for specific sequences that may play a role in the acquisition of variable methylation and test whether they are sufficient to confer variable methylation. We show an inverse correlation between binding levels of the multifunctional transcription factor CTCF and DNA methylation at cVM-IAPs, and identify distinct patterns of chromatin interactions at several of these loci with levels of DNA methylation at these loci correlating with levels of H3K9me3. We expand our screen beyond IAP elements and find that variable methylation is uncommon in other types of retrotransposons. Overall, our findings suggest that variable methylation at IAP elements occurs via complex interactions between IAP sequence, CTCF binding, histone modifications and DNA methylation machinery, and may represent a transient evolutionary state with the potential to cause inter-individual variability in transcription and phenotype.

## Results

### Individual VM-IAPs possess constitutive or tissue-specific methylation variability

Our previous screen for VM-IAPs was performed using whole genome bisulphite sequencing (WGBS) datasets from B and T cells of C57BL/6J mice generated as part of the BLUEPRINT consortium (Adams et al., 2012; Kazachenka et al. 2018). This resulted in the identification of 104 candidate VM-IAP elements. The IAP annotations used in the screen were created by RepeatMasker, a program that identifies repetitive portions of the genome (including TEs). Upon closer inspection we noted that many of the annotated elements were ‘fragmented’, i.e. neither solo LTRs nor fully-structured IAPs with tandem LTRs. Fragmented elements may arise naturally in the genome, through later insertions of other elements, accumulation of polymorphisms over evolutionary time, or through interchromosomal recombination. RepeatMasker often separates the components of an intact IAP into several distinct annotations, resulting in an inflated proportion of fragmented elements in the mouse genome and an inaccurate understanding of IAP boundaries. We therefore developed a method to piece together elements that were artificially fragmented in the annotation. Applying this method to the existing annotation decreased the total count of IAP elements in the mouse genome from 13,065 to 10,678, and reduced the proportion of fragmented elements from 36% to 19% (https://github.com/knowah/vm-retrotransposons/blob/master/data/repeat_annotations/mm10.IAP.mended.tsv).

This improved IAP annotation, along with an updated WGBS screen algorithm (Methods), identified twelve new candidate VM-IAPs, bringing the total to 116. We were able to assay 103 of these IAPs (Supplementary Table S1) using bisulphite pyrosequencing in ear – a distinct tissue from the cell types used in the screen. Of the tested candidates, 51 elements showed variable methylation in ear, passing a threshold of ≥10% inter-individual methylation variation. We termed these ‘constitutive VM-IAPs’ (cVM-IAPs) (Fig. 1A, left panel and Supplemental Data).

**Figure 1.**
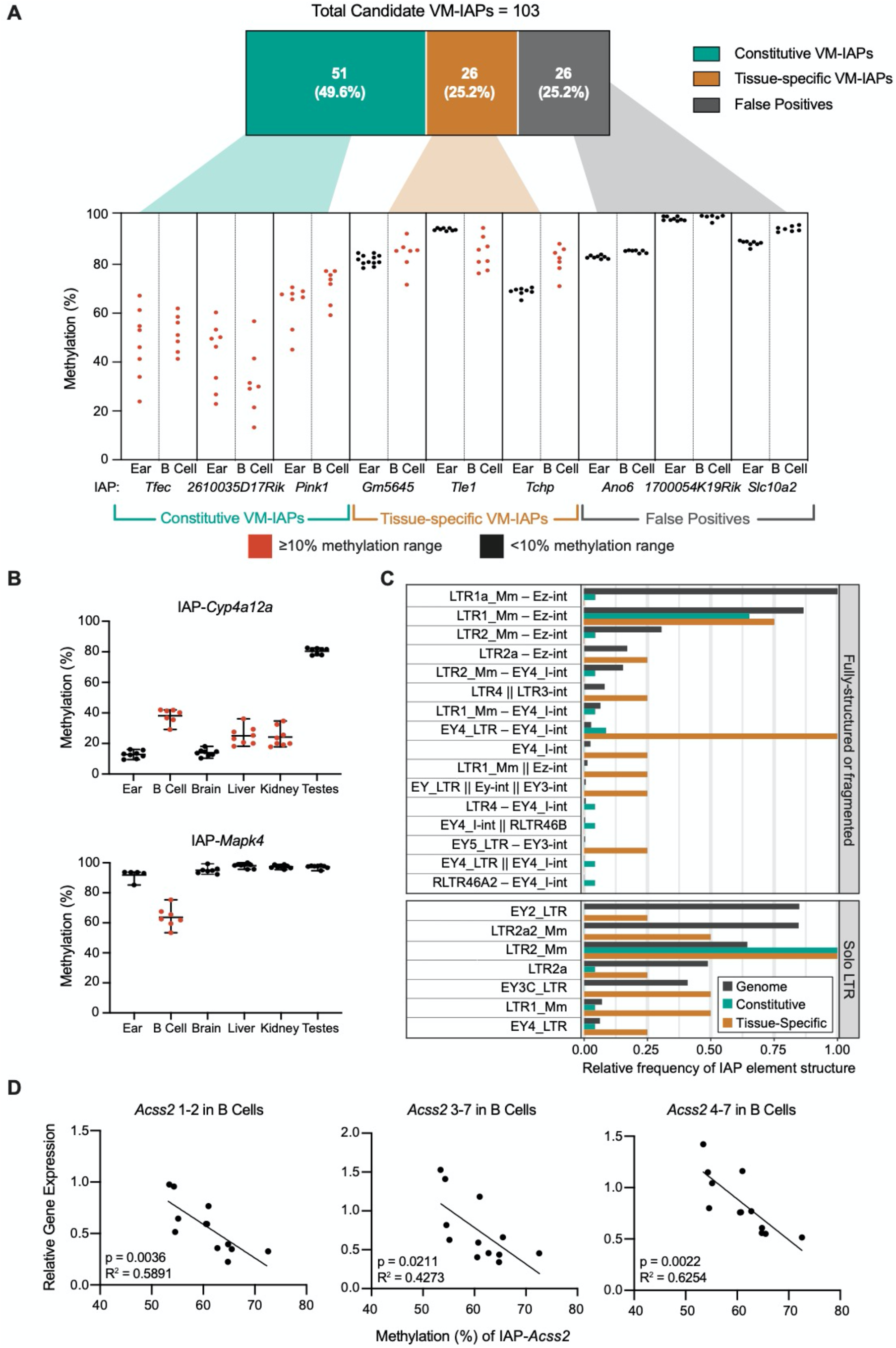
Genome-wide screens and site-specific validation of candidate variably methylated IAP elements (VM-IAPs) identify 51 loci with constitutive variable methylation and 26 loci with tissue-specific variability. **(A)** Bisulphite pyrosequencing of 103 candidate VM-IAPs in ear tissue and B cells led to classification of constitutive VM-IAPs (cVM-IAPs), tissue-specific VM-IAPs (tsVM-IAPs), or false positives – three representative examples are shown for each. The threshold for validation as a VM-IAP is a 10% methylation range amongst individuals. IAP elements with ≥10% and <10% range in methylation are coloured red and black, respectively. Each point represents an individual and is the average methylation of four distal CpGs in the LTR. n = 8 for ear samples and 7 for B cell samples. **(B)** IAP-*Cyp4a12a* (top) is a representative example of tsVM-IAPs with variable methylation in multiple tissues. IAP-*Mapk4* (bottom) is a representative example of tsVM-IAPs whose variable methylation is restricted to B cells. **(C)** IAP elements of the LTR1_Mm–Ez-int (fully-structured) and the LTR2_Mm (solo LTR) types are over-represented in cVM-IAPs. The frequencies of cVM-IAP and tsVM-IAP structures are compared with the relative frequency of genome-wide IAP structures. In the IAP type labels, fully-structured IAPs with flanking LTRs are indicated with “–”, and incomplete IAPs missing one or both flanking LTRs are indicated with “||”. Only IAP types with at least one cVM-IAP or tsVM-IAP are shown. **(D)** In B cells, expression of *Acss2* is inversely correlated with the methylation level of the nearby tsVM-IAP, IAP-*Acss2* (two-tailed Pearson). Expression was quantified by qPCR, normalised to housekeeping genes, *Pgk1* and *Gapdh*, and analysed across multiple exon-exon junctions: 1-2, 3-7 and 4-7.

We next assessed whether the 52 candidates that did not validate in ear samples represent IAPs which are variably methylated in other tissues. Using the same tissues as in the WGBS VM-IAP screen, we sorted B and T cells from C57BL/6J mice (detailed in Kazachenka et al. 2018), and performed bisulphite pyrosequencing for each candidate using these samples. Half (26) of the candidates that did not validate in ear were variably methylated in B cells; we termed these ‘tissue-specific VM-IAPs’ (tsVM-IAPs) (Fig. 1A, middle panel and Supplemental Data). The candidates not variably methylated in ear or B cells were also not variably methylated in T cells; these IAPs were termed ‘false positives’, representing 25% (26 out of 103) of the tested loci and 22% of all identified candidates (Fig. 1A, right panel and Supplemental Data).

To determine whether tsVM-IAPs possess consistent intra-individual methylation variation, we assayed multiple tissues, including brain, liver, kidney, and testes. These represent a diverse set of tissues derived from all three embryonic germ layers. Ten tsVM-IAPs are variably methylated in multiple tissues (Fig. 1B, upper panel and Supplemental Fig. S1A). These tsVM-IAPs display inter-tissue methylation consistency (Supplemental Fig. S1B), although the inter-individual ranges at these loci are lower on average compared to cVM-IAPs (Supplemental Fig. S1C). Of note, the majority of tsVM-IAPs (16) are variably methylated in B cells but not in any other tested tissue (Fig. 1B, lower panel and Supplemental Fig. S1D). This multi-tissue assay also confirmed the lack of methylation variability in five false positive IAPs (Supplemental Fig. S1E). We cannot rule out the presence of additional tsVM-IAPs occurring in cell types that we have not assayed.

### A role for tsVM-IAPs in transcriptional regulation

We previously reported examples of cVM-IAP–initiated transcripts which overlap with annotated genes and found that in some instances, the expression level of these transcripts correlated inversely with the methylation level of the cVM-IAP (Kazachenka et al. 2018). However, in all cases this correlation was tissue-specific. We therefore probed our set of tsVM-IAPs to characterise their effect on transcription of nearby genes. As no tsVM-IAP was located in the vicinity of the transcriptional start site of a gene, we focussed on four tsVM-IAPs located within introns of genes. We investigated whether expression of the surrounding gene correlates with the methylation level of the tsVM-IAP; both expression and methylation were assayed in the tissues in which the tsVM-IAP is variably methylated (Supplemental Fig. S1A). Using three primer sets targeted to different exons, we found a statistically significant inverse correlation between *Acss2* gene expression and IAP-*Acss2* methylation level in B cells (Fig. 1D), but not in brain or kidney where the IAP was also variably methylated. Because expression levels of exons on either side of IAP-*Acss2* were inversely correlated with the methylation level of the element, it is unlikely that this tsVM-IAP is acting as an alternative promoter. There was no transcriptional effect in the other three tsVM-IAPs investigated (Supplemental Fig. S2).

### Repeat-associated variable methylation is mainly a feature of IAPs

To determine whether other families of retrotransposons exhibit the properties of VM-IAPs, we carried out a genome-wide screen for variably methylated LINEs, SINEs, ERVs and non-ERV LTRs in the same WGBS datasets used for the IAP screen. The methylation ranges for candidate variably methylated LINEs, SINEs, ERVs, and non-ERV LTRs were lower compared to IAPs (LINEs, SINEs, non-ERV LTRs, Fig. 2A; IAPs, Supplemental Fig. S3A; ERVs, Supplemental Fig. S3B). We experimentally validated a total of 34 elements representing the top candidates in each repeat family and found two new variably methylated ERVs (VM-ERVs) in addition to 13 previously validated (Kazachenka et al. 2018) (Supplemental Fig. S3C). We also checked whether the ERVs which did not pass the validation threshold in ear samples were variably methylated in B cells, as this was the hallmark of tissue-specific variability in IAPs. None of the tested ERVs were variably methylated in B cells, suggesting that tissue-specific variable methylation occurs exclusively at IAP elements (Supplemental Fig. S3D).

**Figure 2.**
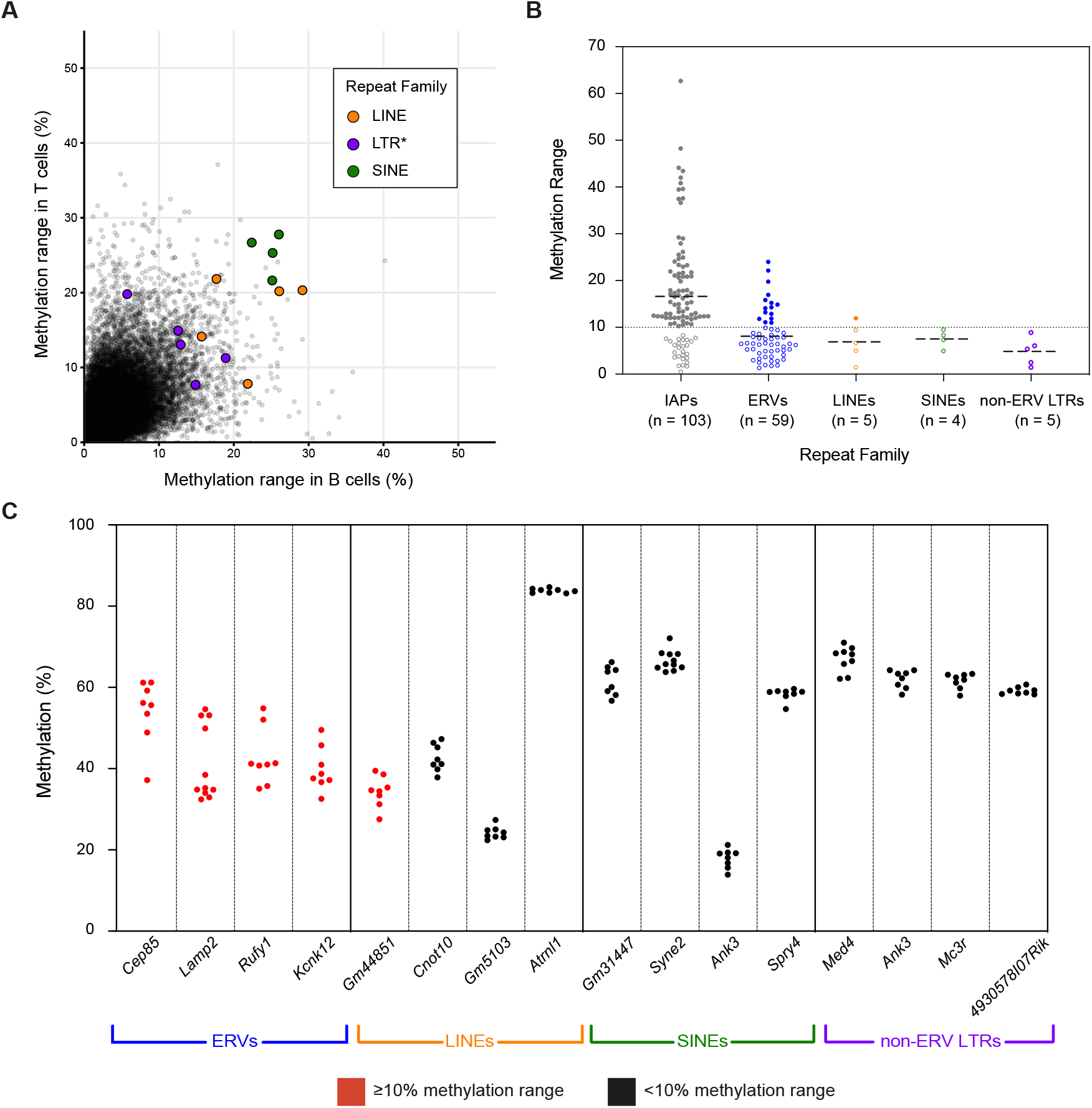
Variable methylation in TEs is unique to evolutionarily young ERV insertions. **(A)** Methylation ranges at the edges of LINEs, SINEs and non-ERV LTRs (denoted LTR*) in B and T cell samples from the WGBS dataset. Each point represents an individual element. Coloured points represent the top candidates in each family, which were selected for bisulphite pyrosequencing validation. **(B)** Methylation ranges of validated elements grouped by repeat family suggests that variable methylation is uncommon in LINEs, SINEs and non-ERV LTRs, while validated. IAP elements have the highest range. Each point represents an individual element. Elements with ≥10% and <10% range in methylation between individuals are denoted by filled and blank circles, respectively. The thin black dotted line shows the 10% threshold. The thick black dashed lines represent the average methylation range per repeat family. **(C)** Bisulphite pyrosequencing validation of the top candidates from the WGBS screen in each repeat family. Elements with ≥10% and <10% range in methylation between individuals are coloured in red and black, respectively. Each point represents an individual and is the average methylation of 2-4 distal CpGs in the element; n ≥ 8 for all elements shown.

Aside from VM-ERVs, LINE-*Gm44851* was the only variably methylated retrotransposon that passed the validation threshold of ≥10% methylation range (Fig. 2C and Supplemental Fig. S3E). Two SINEs located on the X chromosome with high methylation ranges in the WGBS dataset were found to be bistable epialleles – loci which have two methylation states within a population – with the two methylation states segregating by sex; these elements were excluded from further analysis (Supplemental Fig. S3F). In total we have identified 77 VM-IAPs (51 constitutive and 26 tissue-specific), 15 VM-ERVs and one VM-LINE. We compared the validated methylation ranges of individual TEs across the different repeat families and found that VM-IAPs are more variable than the other variably methylated TEs (Fig. 2B). These findings indicate that variable methylation most commonly occurs at IAPs and is not a universal property of TEs.

### Sequence specificity of variable methylation at cVM-IAPs

IAPs can be classified into different types based on the sequence of the LTRs (15 types) and internal portions (10 types) (Hubley et al. 2016). The previously reported VM-IAPs were shown to be enriched in the LTR2_Mm, LTR1_Mm, and EY4_LTR types of IAP LTRs (Kazachenka et al. 2018; ‘IAP’ prefixes omitted for brevity). With the methylation ranges of all candidate IAPs confirmed by pyrosequencing and the improved annotation of IAP elements, we were able to reassess this enrichment by incorporating the internal portions of non–solo LTR elements into the analysis. Genome-wide, the most common types of IAPs are, in order, LTR1a_Mm – Ez-int and LTR1_Mm – Ez-int (both fully-structured IAPs), followed by EY2_LTR and LTR2a2_Mm (both solo LTR IAPs). The black bars in Figure 1C show the frequency of each type of IAP relative to the most common type in the genome, LTR1a_Mm – Ez-int. The majority (74%) of cVM-IAPs are made up of just two types of IAPs, LTR2_Mm and LTR1_Mm – Ez-int (Fig. 1C, green bars), despite these being only the second and fifth most common types of IAP genome-wide. The majority of solo LTR VM-IAPs are of the LTR2_Mm type and the majority of full-length VM-IAPs are of the LTR1_Mm – Ez-int type. In contrast to cVM-IAPs, we did not find that tsVM-IAPs are enriched in any one type of element (Fig. 1C, orange bars). These differences – in particular, the underlying sequence difference between the IAP types – may underpin divergent mechanisms responsible for variable methylation at these elements.

It is known that CpG density is generally positively correlated with DNA methylation, but at high CpG densities there is an inverse correlation with DNA methylation (Meissner et al. 2008; Edwards et al. 2010). Previously, we found that IAPs have higher CpG density than other ERV retrotransposons in the mouse genome (Kazachenka et al. 2018). When comparing CpG density between different types of LTRs, we found that the IAP types for which cVM-IAPs are enriched, namely LTR1_Mm – Ez-int and LTR2_Mm, have higher CpG density than other IAP type LTRs (Supplemental Fig. S4A). Furthermore, cVM-IAPs of the LTR2_Mm type have significantly higher CpG density than non-variable LTR2_Mm elements. On the other hand, cVM-IAPs of the LTR1_Mm – Ez-int type have slightly lower CpG density in their LTRs than non-variable LTR1_Mm – Ez-int elements. Given these contrasts, if CpG density is involved in establishing variable methylation states, the mechanism by which that occurs is dependent on IAP type. Similar to IAPs, VM-ERVs have higher CpG density compared to ERV elements in general, but they have lower CpG density than both cVM-IAPs and tsVM-IAPs (Supplemental Fig. S4B).

Since IAP type and CpG density are both potential sequence-based determinants of methylation variability at cVM-IAPs, we sought to ascertain whether specific sequences within the IAP LTRs may be conferring the variable methylation. We performed a *k*-mer analysis to identify enriched sequences amongst the LTRs of cVM-IAPs (Supplemental Table S2). We identified 14 sequences which are each present in multiple cVM-IAP LTRs and which are present in no more than 2% of all IAP LTRs in the genome. A total of 37 out of the 51 cVM-IAPs contained at least one of these sequences, as well as 6 out of the 26 tsVM-IAPs. Each sequence is mostly present in only one type of LTR, either LTR2_Mm or LTR1_Mm. We experimentally assessed DNA methylation at 18 IAP elements containing multiple sequences enriched in cVM-IAPs and found that they were all hypermethylated with little inter-individual variability (Supplemental Fig S5). This proves that sequence is not the sole determinant of variable methylation. However, the existence of sequences enriched amongst cVM-IAPs suggests that variable methylation may be driven, at least in part, by underlying genetic features.

### CTCF and its motif are enriched at VM-IAPs

Using CTCF ChIP-seq data from ENCODE, we previously reported that VM-IAPs appear closer to CTCF binding sites compared to non-variable IAPs (Kazachenka et al. 2018). To expand and refine our previous findings, we generated CTCF ChIP-seq datasets from livers of eight individuals. Although the presence of a binding site can be easily determined for a given element, it is difficult to discern whether CTCF binds within the IAP itself or its flanking regions. This is because the ChIP-seq fragments from IAPs often do not contain unique sequence, meaning they cannot be mapped confidently to a specific IAP element. To address this, we mapped the CTCF ChIP-seq datasets to consensus sequences of all named IAP LTR types and found that CTCF is enriched in four LTR types (Supplemental Fig. S6A), including those that are specifically enriched among cVM-IAPs (LTR2_Mm and LTR1_Mm) (Fig. 1D).

To generate a robust CTCF binding motif that correlates with strong CTCF binding, we identified a sequence matrix (i.e., motif) derived from the top 10% of ChIP-seq peaks across the pooled eight datasets (Fig. 3A). Binding sites matching this motif are present in around 10% of IAP LTRs at the 5′ end of the U3 region (Fig. 3B-C). When analysing CTCF enrichment at specific IAP elements, we found that most IAPs are not bound by CTCF, including many of those which contain the motif (Fig. 3C). In contrast, almost all the cVM-IAPs are enriched for CTCF binding and only five elements do not have the motif (Fig. 3D). Although tsVM-IAPs are not as ubiquitously bound by CTCF as cVM-IAPs, those that are enriched for CTCF binding tend to be variably methylated in liver, the tissue used to generate the ChIP-seq datasets (Fig. 3E). This validates and refines our previously reported finding that CTCF is specifically enriched at VM-IAPs compared to other IAPs in the genome and suggests that there may also be a relationship between CTCF and tsVM-IAPs.

**Figure 3.**
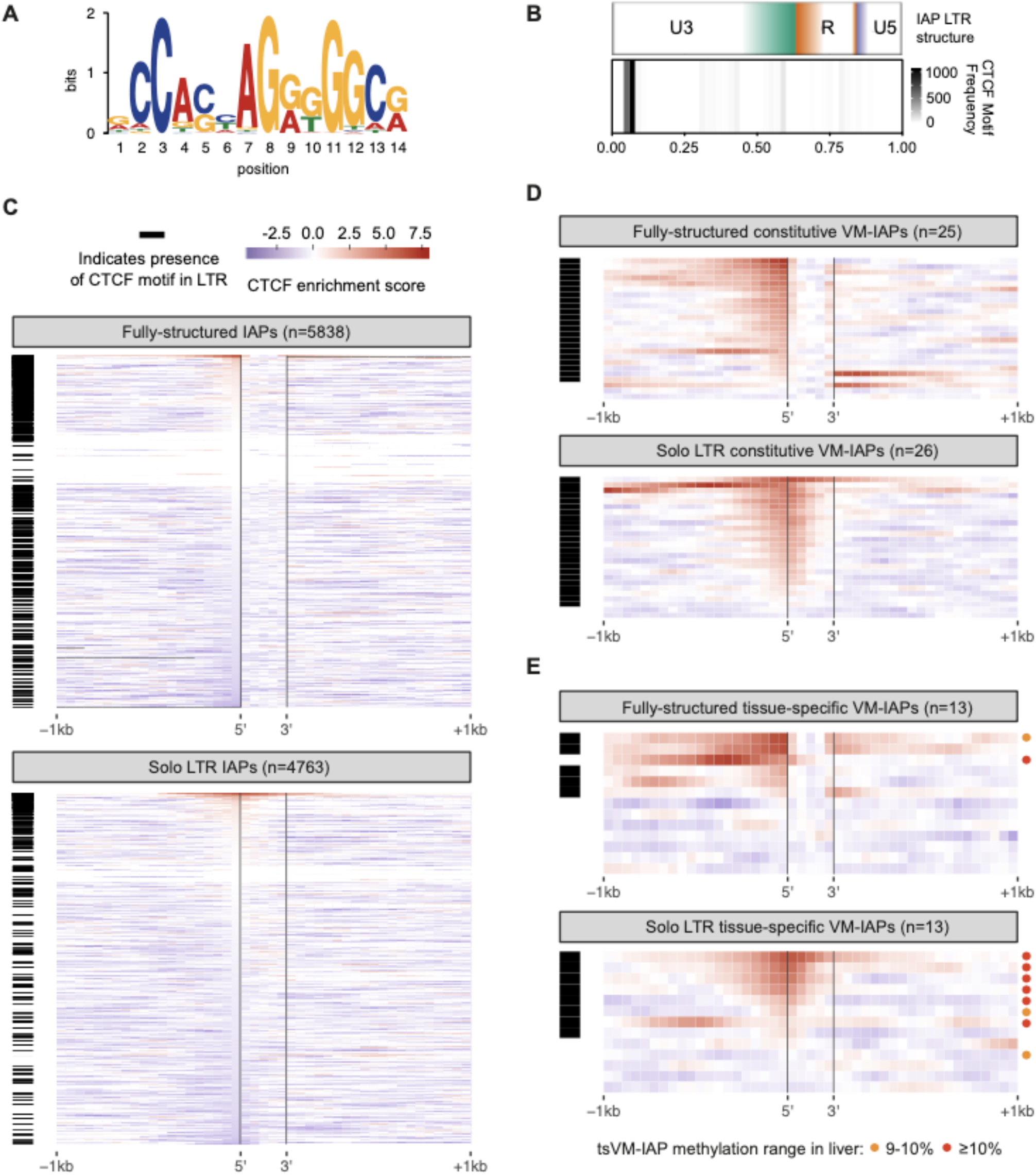
CTCF and its motif are enriched at VM-IAPs relative to other IAPs in the mouse genome. **(A)** CTCF motif derived from the top 10% of CTCF ChIP-seq peaks from the combined peak calls of eight individual liver samples. **(B)** IAP LTRs are divided into U3 (green), R (orange), and U5 (purple) substructures. Colour boundaries represent the median location of the substructure boundaries; colour gradients represent the middle 50% of the distribution of those boundaries (top plot). The CTCF motif is enriched at the 5′ end of the IAP LTRs (bottom plot), within the U3 substructure. **(C-E)** Presence of the CTCF motif within the IAP LTR (horizontal black bars) alongside CTCF ChIP-seq enrichment in liver (heatmaps) at **(C)** fully-structured and solo LTR IAPs (excluding VM-IAPs), **(D)** cVM-IAPs, and **(E)** tsVM-IAPs; circles represent tsVM-IAPs with methylation ranges in liver of 9-10% (orange) and ≥10% (red). CTCF enrichment score scale bar in **(C)** also applies to **(D)** and **(E)**. For the solo LTR plots, the vertical black lines denote the 5′ and 3′ ends of the solo LTR. For the fully-structured IAP plot, the vertical black lines denote the 5′ and 3′ ends of the entire IAP. Note that the ‘fully-structured’ plots include fragmented IAPs.

To ask whether the CTCF enrichment at cVM-IAPs is due to the sequence of the CTCF binding site within those elements, we generated motifs using only the binding sites within IAPs, cVM-IAPs, and tsVM-IAPs (Supplemental Fig. S6B). The similarity of these motifs both to each other and to the more general genome-wide motif suggests that the specific sequence of the binding site is unlikely to be the cause of the specific CTCF enrichment at cVM-IAPs.

### CTCF binding and DNA methylation have an inverse relationship at cVM-IAPs

There is a known inverse relationship between DNA methylation and reduced CTCF binding at many regions in the genome, including imprinted genes and some differentially methylated regions (Bell and Felsenfeld 2000; Hark et al. 2000; Engel et al. 2006; Renda et al. 2007; Han et al. 2008; Lin et al. 2011; Prickett et al. 2013; Maurano et al. 2015; Flavahan et al. 2016; Merkenschlager and Nora 2016). In addition, many CTCF binding sites in the mouse and human genomes are located within TEs (Bourque et al. 2008; Schmidt et al. 2012; Sundaram et al. 2014; Trizzino et al. 2017). We used our ChIP-seq datasets to ask if CTCF binding is variable between individuals at cVM-IAPs, and if so, whether there is a relationship between CTCF binding and DNA methylation. We found that at six out of the seven analysed cVM-IAPs, there is a significant inverse correlation between DNA methylation at the cVM-IAP and CTCF binding (Supplemental Fig. S7A).

This inverse relationship was validated by performing ChIP-qPCR at IAP-*Marveld2* and IAP-*Tfpi* across the same eight individuals on which we performed the ChIP-seq experiments (Fig. 4 and Supplemental Fig. S7B), calculating fold enrichment relative to IAP-*Ell2* (a non-variable intermediately methylated IAP) and to IAP-*Dst* (a non-variable hypermethylated IAP, Supplemental Fig. S7C). We found that CTCF binding is also variable at some non-variably methylated IAPs (Supplemental Fig. S7D), which is consistent with reports that CTCF binding can be methylation sensitive at some loci and not at others (Wang et al. 2012; Maurano et al. 2015). Our data indicate that methylation is associated with the level of CTCF binding at cVM-IAPs. In contrast to CTCF binding, H3K9me3 levels correlated positively with DNA methylation at these loci (Supplemental Fig. S8).

**Figure 4.**
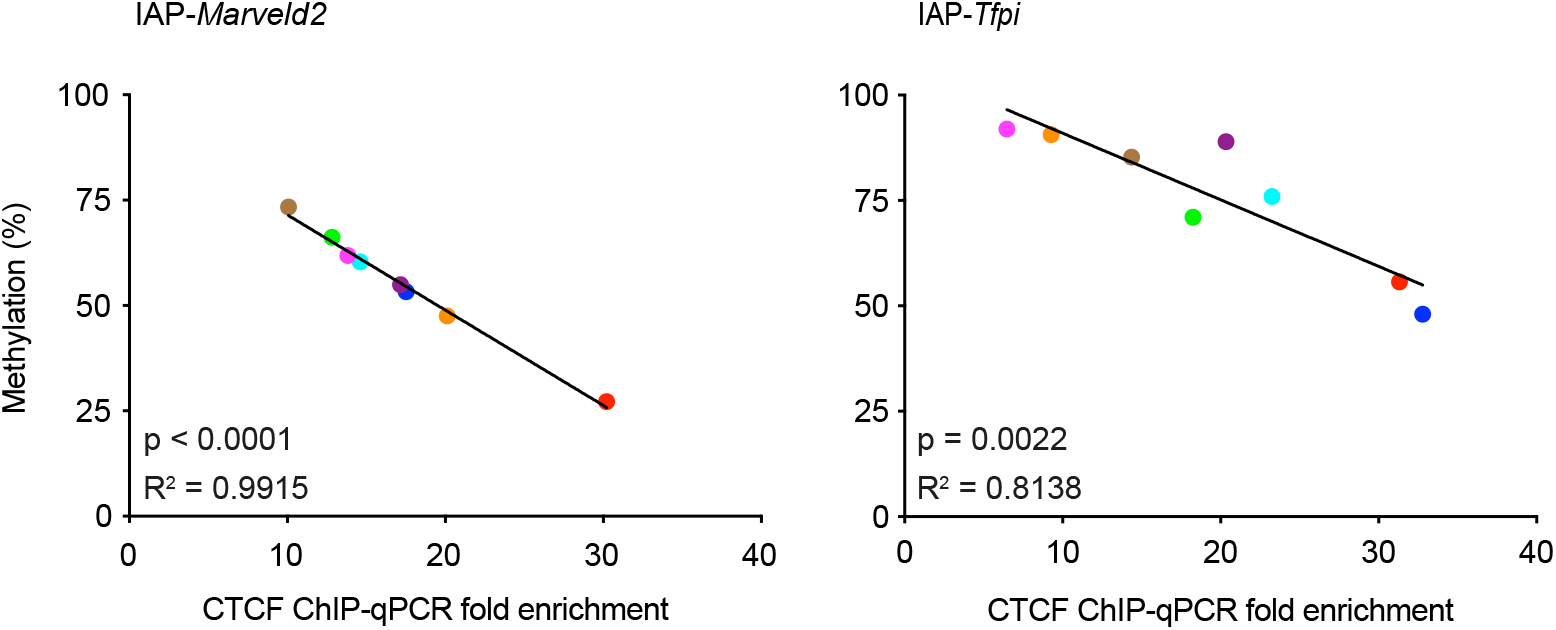
DNA methylation and CTCF binding are inversely correlated at VM-IAPs. There is an inverse relationship between CTCF ChIP-qPCR fold enrichment (relative to CTCF enrichment at IAP-*Ell2*) and DNA methylation (measured by bisulphite pyrosequencing) in eight liver samples at IAP-*Marveld2* and IAP-*Tfpi.* Each coloured point represents the average methylation of the four distal CpGs in the LTR of an individual. The colours represent the same individuals across both plots and show that the relative order of CTCF binding is not the same in each element. P-values are from two-tailed Pearson test.

### Chromatin interactions with cVM-IAPs

CTCF is important for establishing chromatin architecture (Filippova et al. 1996; Moon et al. 2005; Phillips and Corces 2009; Van Bortle et al. 2014) and recent findings suggest that TEs bound by CTCF can contribute to chromatin looping, which in turn can influence gene regulation (Zhang et al. 2019; Choudhary et al. 2020; Diehl et al. 2020). Therefore, we hypothesized that inter-individual variation in CTCF binding at cVM-IAPs could contribute to variation in genome topology. We investigated this using circularised chromatin conformation capture sequencing (4C-seq), a technique used to reveal chromatin interactions between a locus of interest and other parts of the genome. We performed 4C-seq on five individual mice at four cVM-IAPs (IAP-*Marveld2,* IAP-*Tfec*, IAP-*Pink1*, and IAP-*Mbnl1*) and one non-variable IAP (IAP-*Dst*) (Supplemental Fig. S9). In a 400kb window surrounding the elements, we found that the two cVM-IAPs that were lowly methylated within individuals (IAP-*Marveld2* and IAP-*Tfec*) have more long-range and variable interactions compared to the two cVM-IAPs that were highly methylated within individuals, which show fewer interactions (IAP-*Pink1* and IAP-*Mbnl1*). The control non-variable IAP has a more uniform interaction pattern. Due to limited sample size we were unable to confidently quantify differential interactions between individuals.

### Methylation variability is not confined to the LTR boundaries of cVM-IAPs

To determine the extent to which methylation variability exists outside of the LTR, we first assessed the DNA methylation within 1kb of cVM-IAPs by bisulphite pyrosequencing. This revealed that variable methylation between individuals extends outside the TE and into the adjacent unique sequence (Fig. 5A and Supplemental Fig. S10A). The relative order of methylation levels across individuals is maintained until inter-individual variability is lost, at a position around 500-1000bp from the end of the LTR. Although methylation levels beyond this point can show some variability between the individuals (Fig. 5B and C, and Supplemental Fig. S10B and C), this occurs independently of the VM-IAP since the relative order of methylation levels across individuals is no longer retained.

**Figure 5.**
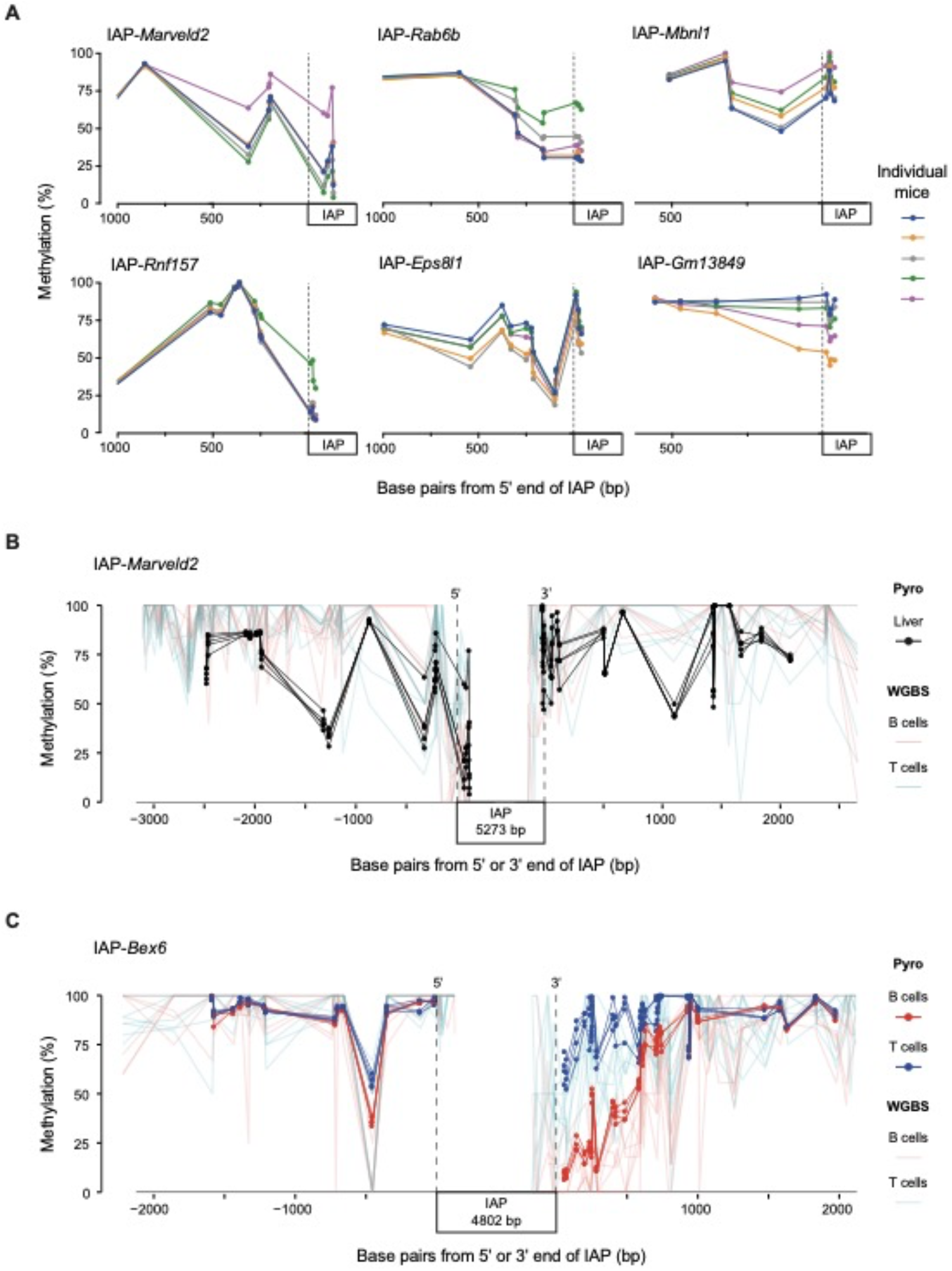
Methylation variability is not confined to the LTRs of VM-IAPs. **(A)** Bisulphite pyrosequencing of liver samples from five individuals shows inter-individual methylation variation within 500-1000bp beyond the edge of six cVM-IAP LTRs. **(B-C)** WGBS data from B and T cells (light red and blue) in the region flanking cVM-IAPs with bisulphite pyrosequencing (Pyro) validation in **(B)** five liver samples (black) at IAP-*Marveld2* and **(C)** B cells and T cells (red and blue) from four individuals at IAP-*Bex6*. IAP length is not to scale. For the pyrosequencing data, each dot represents a CpG. The dashed vertical lines represent the 5′ and 3′ ends of the IAPs.

We next considered whether the methylation variability at cVM-IAPs and beyond is associated with a distinct methylation pattern near the elements. By examining the WGBS datasets, we found that the distribution of DNA methylation in the 5 kb flanking each element is not uniform amongst all 51 cVM-IAPs (Supplemental Fig. S10B). We observed a variety of methylation patterns, including fully hypermethylated flanks, hypomethylated regions, intermediately-methylated regions, and tissue-specific methylation; these patterns are not mutually exclusive. We confirmed this by performing bisulphite pyrosequencing on the flanking regions of six cVM-IAPs representative of these patterns (Fig. 5B and C, and Supplemental Fig. S10C). For example, at the regions flanking IAP-*Marveld2*, we found hyper- and intermediate methylation (Fig. 5B), and at the regions flanking IAP-*Bex6*, we found a tissue-specific methylation pattern that differed between the B and T cells (Fig. 5C), as expected from the WGBS data. These findings suggest that a particular flanking DNA methylation landscape is not required for variable methylation, nor do VM-IAPs influence the flanking DNA methylation landscape in a specific way.

## Discussion

Using WGBS data combined with bisulphite pyrosequencing of independent samples, we classified 51 VM-IAPs as constitutive, meaning variably methylated in all tested tissues, and 26 as tissue-specific, of which 16 were only variably methylated in B cells. Furthermore, fifteen VM-ERVs were also identified. This provides a validated resource of murine (C57BL/6J) metastable epialleles for further studies.

Given the hypothesis that recently active retrotransposons may have facilitated the rapid adaptive evolution undergone by protein-coding immune genes (Chuong et al. 2016; Tie et al. 2018; Ivancevic and Chuong 2020; Ye et al. 2020), it is noteworthy that we find tsVM-IAPs with variable methylation in an immune cell population. We also found an inverse correlation between the methylation level of a tsVM-IAP and gene expression; the transcriptional effect was specific to B cells, despite the presence of variable methylation in other tissue types.

DNA methylation variability between individuals is not confined to the boundaries of cVM-IAPs, but also exists in the immediate flanking regions. A few rare examples of DNA methylation spreading from fully methylated retrotransposons have been reported (Mummaneni et al. 1993; Magewu and Jones 1994; Mummaneni et al. 1995; Graff et al. 1997; Yates et al. 2003; Rebollo et al. 2011; Rebollo et al. 2012; Oey et al. 2015). In all tested cVM-IAPs, we found variable methylation within 500bp of the element boundaries, and sometimes beyond, suggesting that any effects of variable TE methylation may extend into the adjacent sequence. This, along with our finding that the methylation of a tsVM-IAP can correlate with the expression of a nearby gene, indicates that VM-IAPs have potential *cis*-regulatory effects. The absence of a common pattern at these boundaries indicates that VM-IAPs do not confer, or be influenced by, a flanking genomic landscape in a specific way.

We have shown that CTCF is highly enriched at cVM-IAPs and tsVM-IAPs compared to other IAPs in the genome. Unlike VM-IAPs, many non-variable IAP LTRs contain a CTCF motif yet are not bound by the protein; this may be because the majority of IAP LTRs are hypermethylated and CTCF binding is sensitive to methylation. Although it is already known that TEs harbour and spread CTCF binding sites throughout mammalian genomes through mobilisation (Schmidt et al. 2012; Sundaram et al. 2014), it is not clear to what extent these CTCF binding sites affect gene regulatory function. Some studies have indicated that the methylation sensitivity of CTCF is not a major contributor to its function (Wang et al. 2012; Maurano et al. 2015), although this is not the case at imprinted domains (Bell et al. 1999; Bell and Felsenfeld 2000; Hark et al. 2000; Engel et al. 2006). CTCF is known for its role in establishing genomic interactions; we found that these interactions vary depending on the methylation range of the element whereby cVM-IAPs appear to have more interactions than highly-methylated cVM-IAPs. However, whether the CTCF bound to cVM-IAPs contributes to interactions with specific loci is yet to be determined. It has been shown in both mouse and human that TE insertions enriched for CTCF induce differential loop formation that is significantly associated with effects on gene expression (Diehl et al. 2020). Our results indicate that epigenetic differences at TEs might also influence CTCF-mediated conformational states.

Results presented here point to mechanisms for when and how variable methylation at TEs is established. The identification of multiple tsVM-IAPs shows that variable methylation can occur in a tissue-specific manner, which indicates that VM-IAPs can likely arise at different developmental time points. As originally observed at the *A*^*vy*^ and *Axin*^*Fu*^ loci, cVM-IAPs have inter-individual methylation variation but consistent methylation levels within an individual across tissues, indicating that variable methylation establishment at these loci likely occurs prior to germ layer specification (Waterland and Jirtle 2003; Waterland et al. 2006; Kazachenka et al. 2018). Since the extent of methylation variation and the absolute methylation levels differ between tissues at tsVM-IAPs, the mechanisms underlying establishment of variable methylation at these loci are likely to be different from those at cVM-IAPs, and probably occur later in development. These findings also suggest that, rather than being resistant to the global epigenetic reprogramming that occurs in early development as has been proposed (Walsh et al. 1998; Lane et al. 2003), around 1% or more of the ~10,000 IAP elements have increased susceptibility to reprogramming.

Transcription factor binding at cVM-IAPs and the genetic sequence of cVM-IAP LTRs are likely contributors to the mechanism underlying establishment of variable methylation. We rule out the possibility that genomic methylation context is an important factor in establishing variable methylation by showing that there is no discernible common pattern of DNA methylation flanking cVM-IAPs. Moreover, a correlation between methylation and H3K9me3 was observed along with an inverse correlation between the enrichment of transcription factor CTCF and DNA methylation at cVM-IAPs. During early development, CTCF may compete with DNA methylation machinery for access to VM-IAP LTRs. This hypothesis is consistent with research showing that CTCF can influence the presence or absence of DNA methylation during development with functional consequences (Bell et al. 1999; Bell and Felsenfeld 2000; Hark et al. 2000; Engel et al. 2006; Stadler et al. 2011; Wang et al. 2012; Feldmann et al. 2013; Teif et al. 2014; Maurano et al. 2015; Flavahan et al. 2016; Wiehle et al. 2019). Therefore, the molecular antagonism between CTCF and DNA methylation machinery could contribute to the formation of variable methylation levels between genetically identical individuals. CTCF may also facilitate interactions with genomic regions that contribute to the regulation of VM-IAP methylation.

The updated categorisation of VM-IAPs revealed that the majority of cVM-IAPs are solo LTR2_Mm or full length LTR1_Mm – Ez-int elements, while tsVM-IAPs are not enriched for any specific type of IAP. This is further evidence that cVM-IAPs and tsVM-IAPs likely arise via separate mechanisms. We have also identified sequence correlates and CpG density profiles of variable methylation at IAPs. Subsets of cVM-IAPs are highly enriched for specific sequences compared to other IAPs in the genome, however these are unable to predict variable methylation. At this point it is unclear exactly how the enriched sequences contribute to establishing variable methylation. A plausible explanation is that they contain binding sites for transcription factors such as CTCF or KRAB zinc finger proteins (KZFPs) (Wolf and Goff 2009; Sundaram et al. 2014). KZFPs are the largest family of transcription factors in mammals and are known to coevolve with and regulate TEs by recruiting heterochromatic machinery to distinct loci in a sequence specific manner (Rowe et al. 2010; Thomas and Schneider 2011; Wolf et al. 2015; Imbeault et al. 2017). KZFPs are rapidly evolving and highly polymorphic between vertebrates and even among different mouse strains (Elmer and Ferguson-Smith 2020). Variable methylation could arise if there were changes in KZFP binding sites within the VM-IAPs or the binding domains of specific KZFPs that target VM-IAPs. Ultimately, the enrichment of specific sequences at VM-IAPs could contribute to variable methylation via novel or disrupted protein interactions.

The genome-wide screens that we have conducted at TEs in mouse reveal that variable methylation is a rare occurrence that is mostly restricted to IAPs, which are amongst the evolutionarily youngest ERV types (Qin et al. 2010). Whether variable methylation is a phenomenon exclusive to young TEs is an open question. Genome-wide screens for metastable epialleles at non-repetitive regions have been performed on the human genome, and variably methylated non-repetitive regions appear to exist (Silver et al. 2015; Kessler et al. 2018). The extent to which these regions are driven by inter-individual genetic differences has not yet been fully determined as it is difficult to eliminate the confounding effect of human genetic variation. Due to the difference in how TEs and non-repetitive regions are regulated by DNA methylation (Greenberg and Bourc’his 2019), it is likely that the overarching mechanisms of variable methylation establishment and its potential function also vary between these two types of genomic loci.

We have shown that VM-IAPs have the capacity to affect gene regulation by either *cis* or long-range mechanisms and in a tissue-specific manner. Addressing the question of what defines a VM-IAP is made more difficult by the fact that there is no unifying characteristic which distinguishes them from the other 99% of IAPs. Instead, we have shown that there are correlations of varying specificity and confidence by which subpopulations of VM-IAPs differ from the main IAP population with regards to tissue-specific methylation states, CTCF binding, histone modifications, genomic sequence, and CpG density. This shows that there are multiple factors contributing to variable methylation between individuals. The functional implications of TE epigenetic variability and the extent to which this can influence phenotypic outcome remain to be determined.

## Methods

### Improving the catalogue of IAPs

The GRCm38/mm10 RepeatMasker (henceforth RM) annotation (Dfam v4.0.7) was downloaded from the UCSC table browser (Smit 2013-2015; Hubley et al. 2016). This annotation set contains entries for each transposable element, which may be categorized into one or more ‘subelements’, which are united by an ‘element ID’. All transposable elements that were considered IAP elements contained at least one IAP subelement of the following types: IAPLTR1a_Mm, IAPEY2_LTR, IAPEy-int, IAPEY3_LTR, IAPLTR2b, IAPLTR2_Mm, IAPEz-int, IAPEY4_I-int, IAP-d-int, IAPEY3-int, RLTR10B2, IAP1-MM_I-int, IAP1-MM_LTR, IAPLTR2a2_Mm, IAPLTR4, IAPLTR1_Mm, IAPEY3C_LTR, IAPLTR4_I, IAPLTR3, IAPLTR3-int, IAPEY5_LTR, IAPEY5_I-int, IAPEY_LTR, IAPEY4_LTR, IAPA_MM-int.

Elements in the RM annotation that overlap 500kb boundaries were erroneously categorized as two entries with separate element IDs. Each element at one of these boundaries was patched to unify the two entries under the same element ID.

Furthermore, many of the elements are ‘fragmented’ for the following reasons: subsequent insertion of other transposable elements; sequence divergence from the Dfam models; general poor performance of RepeatMasker at ERV elements; or the retrotransposition of an already fragmented IAP element. An element is ‘fragmented’ if is neither fully-structured (containing an internal portion comprised of ERV genes flanked by tandem LTRs) nor a solo LTR (a single LTR with no ERV genes, formed from intra- or inter-element recombination of two LTRs (Ji and DeWoody 2016).

For each fragmented element annotated as missing a 5′ LTR, the following heuristic was used in an attempt to ‘mend’ it, forming a fully-structured IAP: (1) merge the element with an adjacent fragmented element missing a 3′ LTR; (2) merge the element with an adjacent solo LTR; (3) merge the element with an adjacent fully-structured IAP. Adjacent elements must have been within 2000 bp of the 5′ end of annotation to form a match. The same algorithm was used for fragmented elements missing a 3′ LTR, or *vice versa*. Note that sometimes an element was annotated as containing just an internal portion, so the attempt at mending was performed on both edges. Step 3 of the heuristic could result in the formation of a double or higher-order fully-structured IAP.

### Screen for variably methylated transposable elements

A screen for VM-IAPs, similar to that described in Kazachenka et al. 2018, was performed using the improved catalogue of IAPs. The 16 C57BL/6J whole-genome bisulphite sequencing (WGBS) datasets (8 T cell samples, 8 B cell samples) from the BLUEPRINT Epigenome project (Adams et al. 2012) were mapped to the mm10 reference using Bismark v0.20.0 with default options (Krueger and Andrews 2011). This dataset contains 8 ‘standard’ WGBS and 8 oxidative WGBS samples, which were not distinguished in the analysis since hydroxymethylation levels are very low in these samples. Methylation calls were obtained from the aligned reads with a MAPQ ≥10. LTR methylation states in each sample were calculated at the 5′ and 3′ edge of each IAP element by determining the average methylation level of the 8 CpGs nearest each LTR edge (only the two outward-facing edges were considered for each element; the internal-facing LTR edges of fully-structured IAPs were excluded). At each LTR edge, a sample with fewer than 20 methylation calls across the 8 CpGs, or fewer than 4 CpGs with coverage, was considered uninformative. LTR edges with fewer than 5 informative samples in either cell type were then excluded from further analysis. The methylation range at each LTR edge surviving these filtering steps was then calculated in each cell type. Unlike in the previous screen, inter-replicate methylation ranges were calculated without excluding the highest and lowest methylation values in each cell type, as this conservative measure was deemed unnecessary in light of the improved IAP reference and stricter filtering on both read quality and region-level coverage.

The screen in LINE, SINE, and non-ERV LTR elements was performed in a similar manner with some modifications: the RepeatMasker annotation was only fixed by combining adjacent elements of the same class within 100 bp of each other; and only the first 8 CpGs within 200 bp of each element edge were considered.

### *k*-mer and CpG density analysis

All sequences of length 15 nt (*k*-mers with *k*=15) present in two or more of the 5′ LTRs of the cVM-IAPs (N=51) were identified using Jellyfish v2.3.0 (Marcais and Kingsford 2011). Enrichment of each *k*-mer (N=2,363) was then calculated relative to a background set of IAP 5′ LTRs (N=9,650). *k*-mers present in at least 5 cVM-IAPs and with an enrichment of at least 20-fold (N=229) were then grouped by sequence, such that all *k*-mers in a group overlap with another *k*-mer in the group by *k*-1 nucleotides. Each group was then merged into a single extended sequence using *abyss-align* from the tool ABySS v2.2.3 (Jackman et al. 2017). The extended sequences were then trimmed to the sub-sequence maximising the enrichment in cVM-IAPs versus background IAPs. Identified sequences are listed in Supplemental Table S2; pyrosequencing primers are listed in Supplemental Table S1.

CpG density was calculated for IAP LTRs by normalising the number of CpGs in the LTR by the LTR length in base pairs.

### Tissue collection, DNA/RNA extraction and bisulphite pyrosequencing

Immediately following dissection, C57BL/6J tissues were snap frozen in liquid nitrogen and manually pulverised. 30ug of tissue (brain, liver, kidney, testes, B and T cells) was used for simultaneous purification of genomic DNA and total RNA with the AllPrep DNA/RNA Mini Kit - Quick-Start Protocol (QIAGEN, cat. no. 80204). Ear notch samples were lysed (Lysis Buffer: 10mM EDTA, 150mM NaCl, 10mM Tris-HCl pH 8, 0.1% SDS) and DNA was purified using a standard phenol chloroform extraction protocol. 0.5-1μg of DNA per sample was bisulphite converted using the two-step protocol of the Sigma Imprint^®^ DNA Modification Kit according to the manufacturer’s instructions. Following PCR amplification, CpG site-specific methylation was quantified using the PyroMark™ Q96 MD pyrosequencer (Biotage) as previously described (Kazachenka et al. 2018). Primers are listed in Supplemental Table S1 (for IAPs, ERVs, LINEs, SINEs, non-ERV and spreading).

### B and T cell sorts

Splenic tissue was ground through a 70μM cell strainer to obtain a single cell suspension in PBS with 2% heat-inactivated fetal calf serum (FCS). Red blood cell lysis was performed on ice using cold ammonium chloride (Stem Cell Technologies, cat. No 07800). Cells were subsequently stained as described in the panel (Table 1) for 30 minutes at 4°C. Cells were washed twice with 2% FCS in 2ml PBS. 7-aminoactinomycin D (7-AAD) (1:100; Biolegend, 420404) was added as a viability stain. Using an Influx Cell Sorter (BD Biosciences), cells were sequentially gated based on cell size, presence of singlets, and live cells to sort CD3+, CD4+, CD25-, CD44lo, CD62L+ T cells and CD19+ CD43-B cells. The sorted populations include naïve CD4+ T cells and T1, T2, marginal zone and follicular B cells. Gates were confirmed using ‘fluorescence minus one’ controls for CD44 and CD43 markers.

**Table 1.** Fluorophores for B and T cell panel.

| Antigen | Fluorochrome | Clone | Company | Catalogue Number |
| --- | --- | --- | --- | --- |
| CD19 | BV711 | 6D5 | Biolegend | 115555 |
| CD43 | APC | S11 | Biolegend | 143207 |
| CD3 | FITC | 17A2 | Biolegend | 100203 |
| CD4 | APC/Cy7 | RM4-5 | Biolegend | 100525 |
| CD25 | BV421 | PC61 | Biolegend | 102033 |
| CD44 | PE/Cy7 | IM7 | Biolegend | 103029 |
| CD62L | PE | MEL-14 | Biolegend | 104407 |

### RT-qPCR

RNA was treated with RNase-free DNase I (ThermoScientific, EN0521) prior to cDNA synthesis using RevertAid H Minus First Strand cDNA Synthesis Kit with oligo dT and random hexamer primers (ThermoScientific). qPCR primers were designed using Primer-BLAST and are listed in Supplemental Table S3. qPCR was performed with Brilliant II SYBR^®^ Green QPCR Master Mix (Agilent Technologies) in a LightCycler^®^ 480 Instrument II (Roche). Relative gene expression was calculated using the standard curve method and cDNA input was normalised using housekeeping genes Pgk1 and Gapdh. The significance of correlations between gene expression and VM-IAP methylation levels was assessed by computing Pearson correlation coefficients followed by two-tailed p values in GraphPad Prism.

### Chromatin immunoprecipitation (ChIP) - qPCR and sequencing

Chromatin immunoprecipitation (ChIP) was performed as previously described with some modifications (Imbeault et al. 2017). 100mg of powdered frozen mouse liver was crosslinked in 1% formaldehyde for 10 minutes and subsequently quenched with Tris pH 8.0 (250mM final) for 10 minutes. The quenched cells were washed twice with ice-cold PBS supplemented with a protease inhibitor cocktail (EDTA-free cOmplete™, Sigma Aldrich), flash frozen in liquid nitrogen, and stored at −80°C. Crosslinked cells were thawed on ice and then lysed sequentially on ice and for 10 minutes at each step in each of the following buffers: LB1 (50 mM HEPES-KOH pH 7.4, 140 mM NaCl, 1 mM EDTA, 0.5 mM EGTA, 10% Glycerol, 0.5% NP-40, 0.25% Triton-X-100 and EDTA-free cOmplete™), LB2 (10 mM Tris-HCl pH 8.0, 200 mM NaCl, 1 mM EDTA, 0.5 mM EGTA and EDTA-free cOmplete™), and SDS shearing lysis buffer (10 mM Tris-HCl pH 8, 1 mM EDTA, 0.15% SDS and EDTA-free cOmplete™). The lysates were sonicated (Bioruptor) at 4°C to generate DNA fragments of 100 – 500 bp (3 repetitions of 5 sonication cycles of 30 seconds on and 30 seconds off for the CTCF ChIP; 1 repetition of 8 cycles of 30 seconds on and 30 seconds off for the H3K9me3 ChIP) and the sonicated lysates were subsequently clarified by centrifugation (15,000 rpm for 15 min at 4 °C). The sonicated lysate was divided into 1.5-mL Eppendorf tubes based on the number of ChIPs performed and topped up to 1mL with Lysis Buffer 500NaCl (20 mM Tris-HCl, pH 7.5, 500 mM NaCl, 1 mM EDTA, 0.5 mM EGTA, 1% Triton-X-100, 0.1% Sodium deoxycholate, 0.1% SDS and EDTA-free cOmplete™). This mixture was incubated overnight at 4°C with beads (Protein A Dynabeads, Invitrogen, for CTCF ChIP; Protein G Dynabeads, Invitrogen, for H3K9me3 ChIP) that had been pre-blocked with 0.5% BSA and mixed with polyclonal CTCF antibody (C15410210-50, Diagenode) or polyclonal H3K9me3 antibody (AB_2532132, Active Motif). To remove non-specifically bound proteins from the CTCF ChIP, beads were washed five times with RIPA Buffer (50 mM HEPES-KOH, pH 7.4, 500 mM LiCl, 1 mM EDTA, 1% NP-40 and 0.7% Sodium deoxycholate) and once with TE Buffer (50 mM Tris-HCl, pH 8.0, 10 mM EDTA) at 4°C. To remove non-specifically bound proteins from the H3K9me3 ChIP, beads were washed twice with low salt buffer (10 mM Tris-HCl pH 8.0, 150 mM NaCl, 1 mM EDTA, 1% Triton X-100, 0.15% SDS, 1 mM PMSF) followed by single successive washes with high salt buffer (10 mM Tris-HCl pH 8.0, 500 mM NaCl, 1 mM EDTA, 1% Triton X-100, 0.15% SDS, 1 mM PMSF), LiCl buffer (10 mM Tris-HCl pH 8.0, 1 mM EDTA, 0.5 mM EGTA, 250 mM LiCl, 1% NP40, 1% Na-deoxycholate, 1 mM PMSF), and 10 mM Tris pH 8.0 at 4°C. The DNA-protein complex was eluted from the beads in Elution Buffer (50 mM Tris-HCl, pH 8.0, 10 mM EDTA and 1% SDS) for 20 min at 65°C, and reverse-crosslinked overnight at 65°C. The eluted samples were then treated with RNase A (Wako) and Proteinase K (Roche), and purified using a PCR purification kit (NEB).

qPCR was performed with Brilliant II SYBR^®^ Green qPCR Master Mix (Agilent Technologies) in a LightCycler^®^ 480 Instrument II (Roche). The ChIP-qPCR fold enrichments are calculated by 2^ΔCt^, where ΔCt is the difference in qPCR Ct value between the tested IAP and a control IAP (either IAP-*Dst*, IAP-*Ell2*, or IAP-*Asxl3*). qPCR primers were designed using Primer-BLAST and are listed in Supplemental Table S3.

ChIP-seq libraries were prepared using KAPA Adapters, KAPA HyperPrep Kit (KAPA Biosystems), and AMPure XP Beads (Beckman Coulter) and quality checked using Qubit, Bioanalyzer, and Tapestation. The libraries were sequenced as 150bp paired-end reads on the Illumina HiSeq4000. The resulting ChIP-seq data was trimmed by Trim Galore and aligned using bwa 0.7.15. Using only reads with MAPQ≥10, ChIP peaks and summits were called by MACS2 2.1.0 as described in (Thorvaldsdottir et al. 2013) and were visualized using IGV and custom R scripts (see Data Access). The MACS2 summits from all 8 CTCF ChIP-seq samples were combined, and the sequence of a 50bp window centred on each summit (N=97,746 summits) was used as input to MEME 5.0.4, using the strategy of motif finding from (Schmidt et al. 2012). The FIMO tool from MEME was then used to identify genome-wide locations of the top motif.

### 4C-seq

4C-seq was performed as previously described with some modifications (Splinter et al. 2012). Tissues were fixed as outlined in the ChIP protocol above. DpnII (New England Biolabs) was used as the primary restriction enzyme and NlaIII (New England Biolabs) as the secondary restriction enzyme. Prior to library preparation, samples were purified by the Monarch PCR & DNA Cleanup Kit (New England Biolabs). For the library preparation, 16 individual PCR reactions were performed for each sample per viewpoint with reverse primers containing indexes (see Supplemental Table S4). The 16 PCRs were combined and purified using 0.8x Agencourt AMPure XP beads (Beckman Coulter). Five libraries for each of the five tested viewpoints were multiplexed, quality checked using Qubit and Bioanalyzer, and sequenced as 150bp paired-end reads on the Illumina HiSeq4000. The sequencing data for each viewpoint were processed using the *smoothCounts* method from the R Bioconductor package FourCSeq (Klein et al. 2015). Read counts for each restriction fragment were then normalised and log transformed. The trend line and its 95% confidence interval were generated by the *loess.sd* method from the R package msir using span=0.01 (Scrucca 2011). ChromHMM data is from mouse liver, and was downloaded from a public GitHub repository https://github.com/gireeshkbogu/chromatin_states_chromHMM_mm9/blob/master/liver_cStates_HMM.zip and lifted over to mm10 (Bogu et al. 2015). Gene tracks are from the R Bioconductor package EnsDb.Mmusculus.v79.

## Data Access

All raw and processed sequencing data generated in this study have been submitted to the NCBI Gene Expression Omnibus (GEO; https://www.ncbi.nlm.nih.gov/geo/) under accession number GSE159110 (CTCF ChIP-seq and 4C-seq) and GSEXXXX (BLUEPRINT methylation data). All code used to perform the analyses is available at GitHub (https://github.com/knowah/vm-retrotransposons).

## Acknowledgements

Research was funded by grants from the MRC (MR/R009791/1), BBSRC (BB/R009996/1), and a Wellcome Trust Investigator Award (210757/Z/18/Z) to A.C.F.-S. We are grateful for PhD studentships from the BBSRC to J.L.E., from the Cambridge Trust and Department of Genetics to A.D.H., and from the Cambridge Trust, Downing and Pomona Colleges to T.M.B. We are grateful to Kay Harnish, Julie Ahringer and Alex Appert, Iwo Kucinski, Bertie Gottgens for access to facilities and technology optimisation. We thank Michael Imbeault, Felipe Karam Teixeira, Tyler Linderoth, Carol Edwards and Fran Dearden for helpful discussions and Gloria Jansen for technical assistance. Thanks to members of the Ferguson-Smith lab for feedback and discussions during manuscript preparation.

