## Supplemental Data for "Genomic properties of variably methylated retrotransposons in mouse"

Constitutive VM-IAPs (cVM-IAPs)

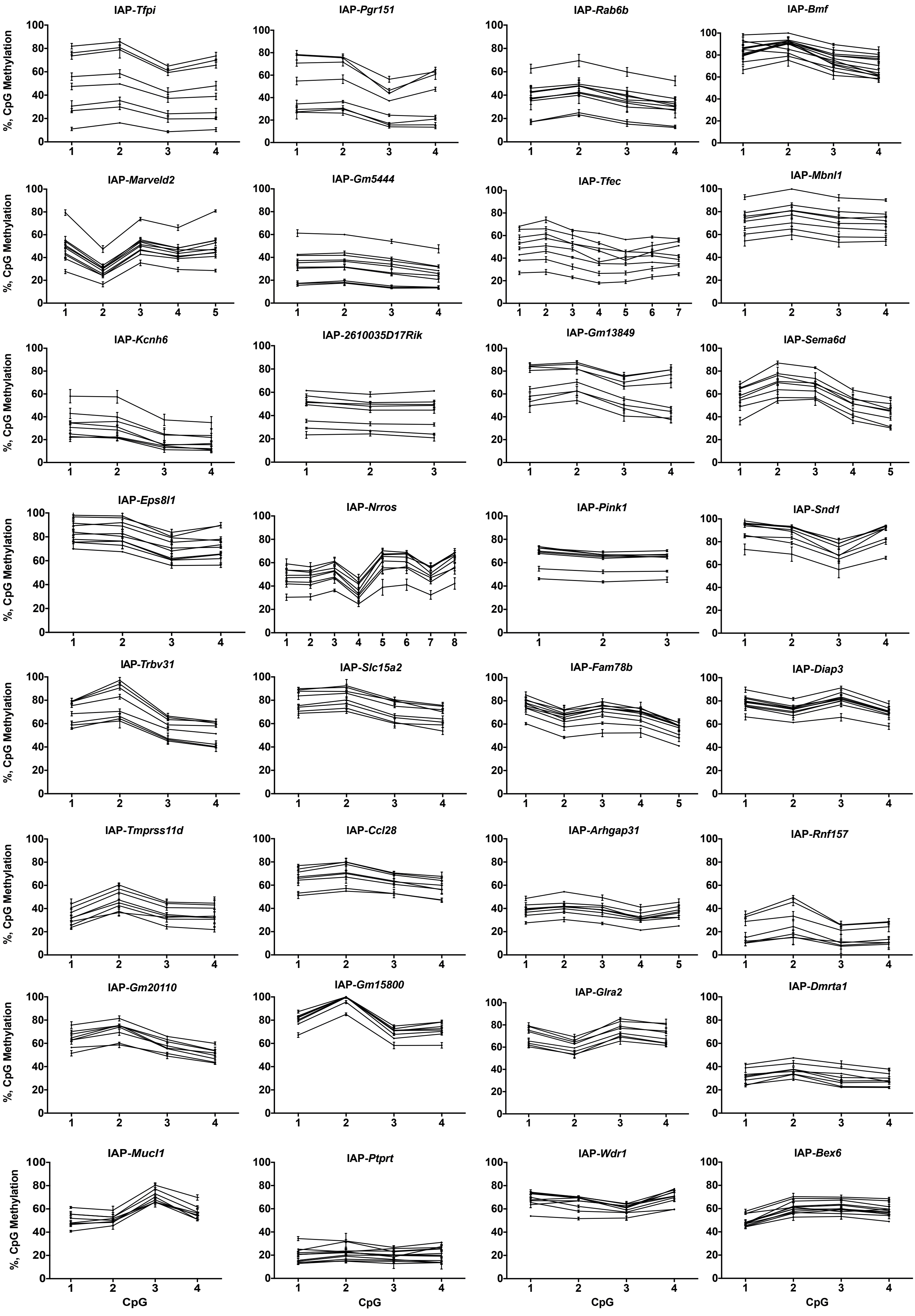

Constitutive VM-IAPs (cVM-IAPs)

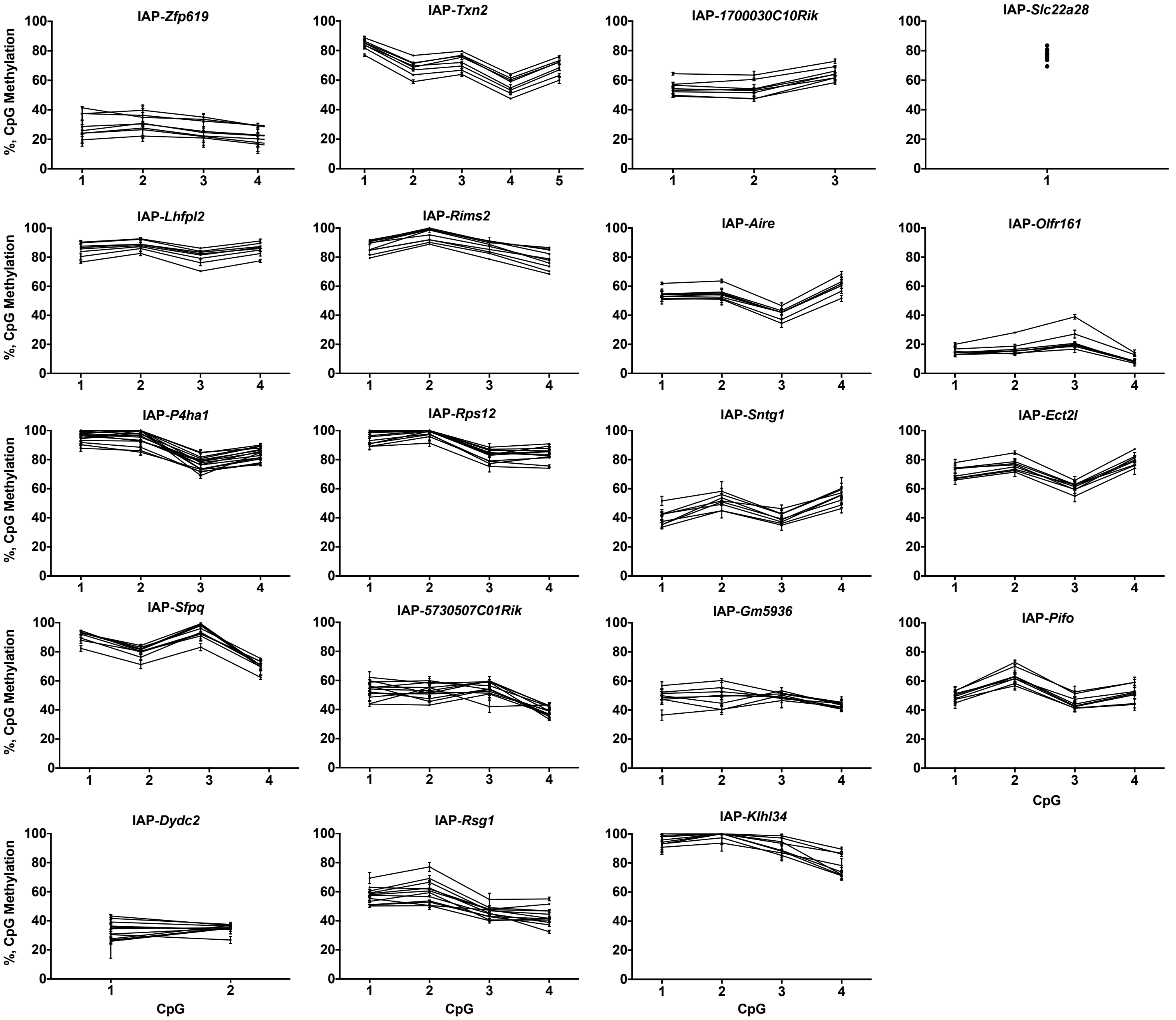

Tissue-specific VM-IAPs (tsVM-IAPs)

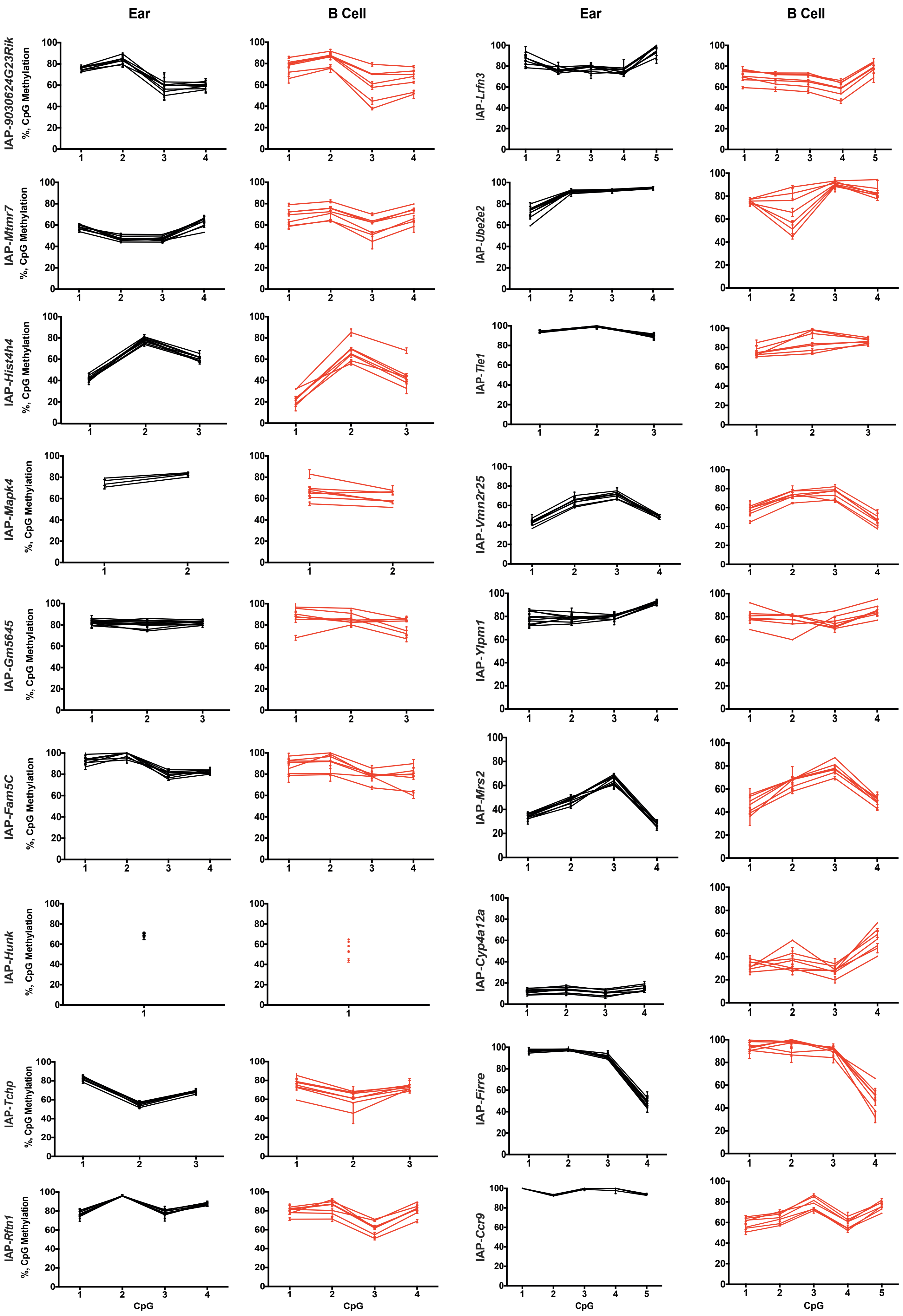

Tissue-specific VM-IAPs (tsVM-IAPs)

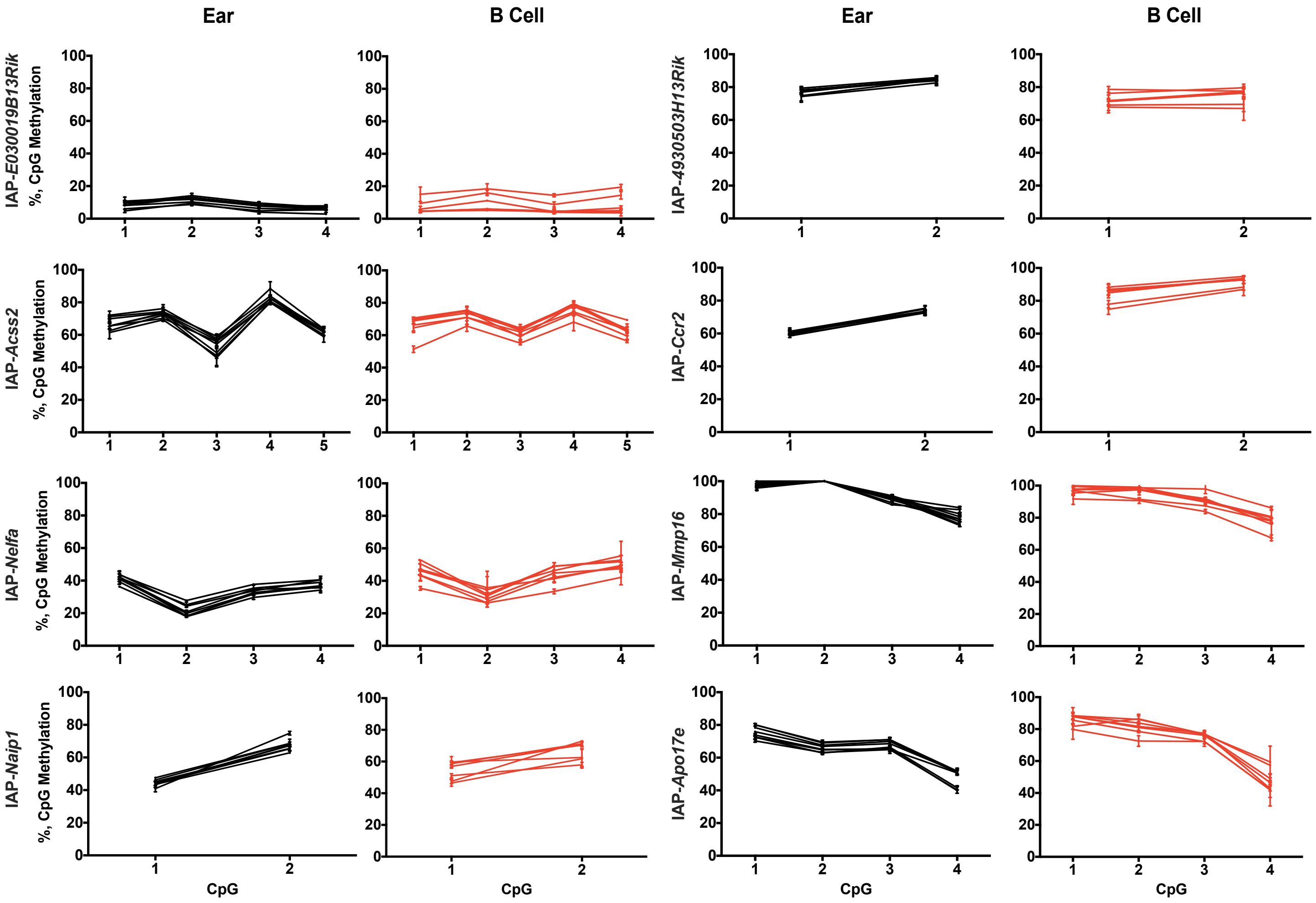

False positive IAP elements

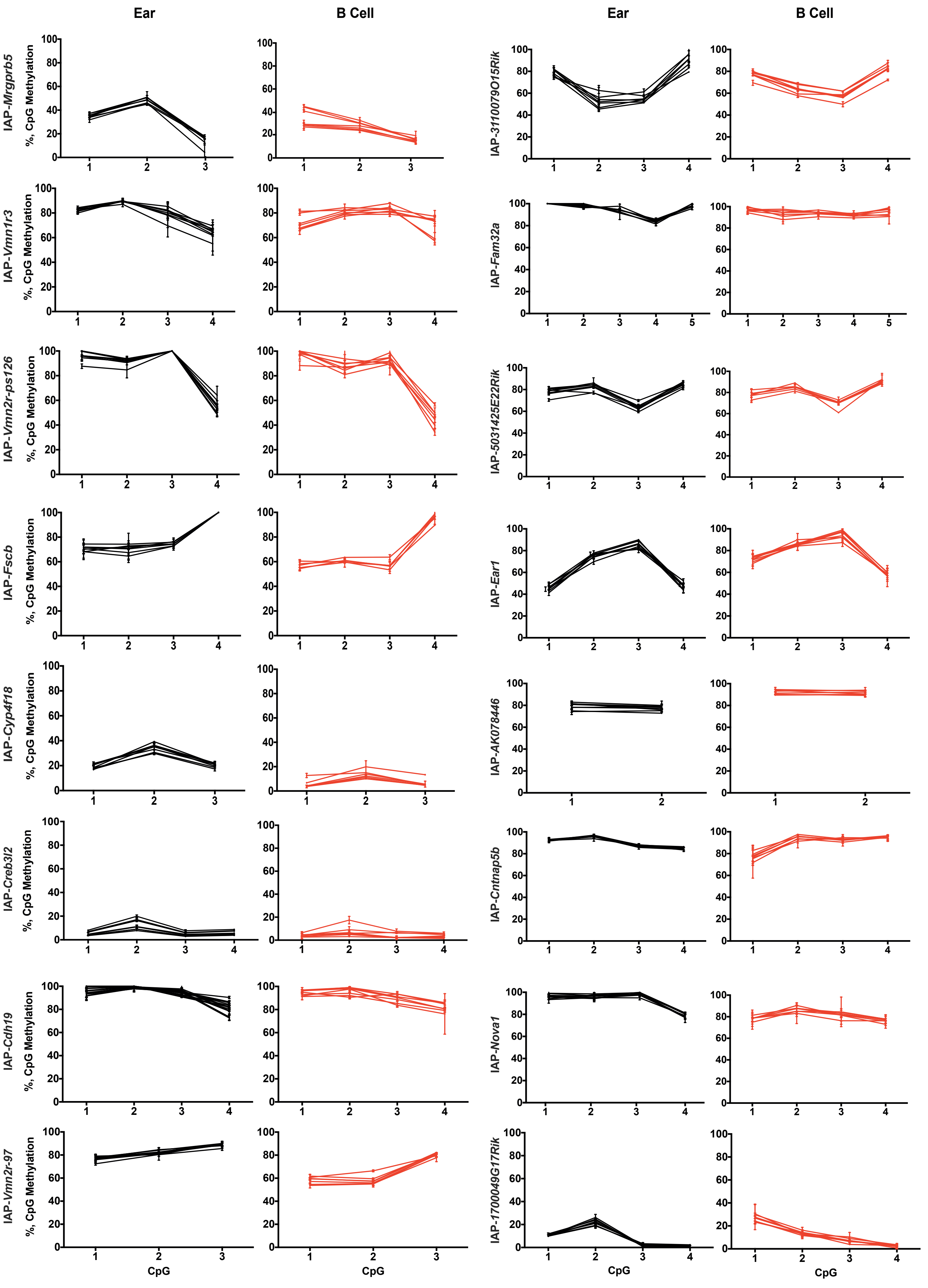

False positive IAP elements

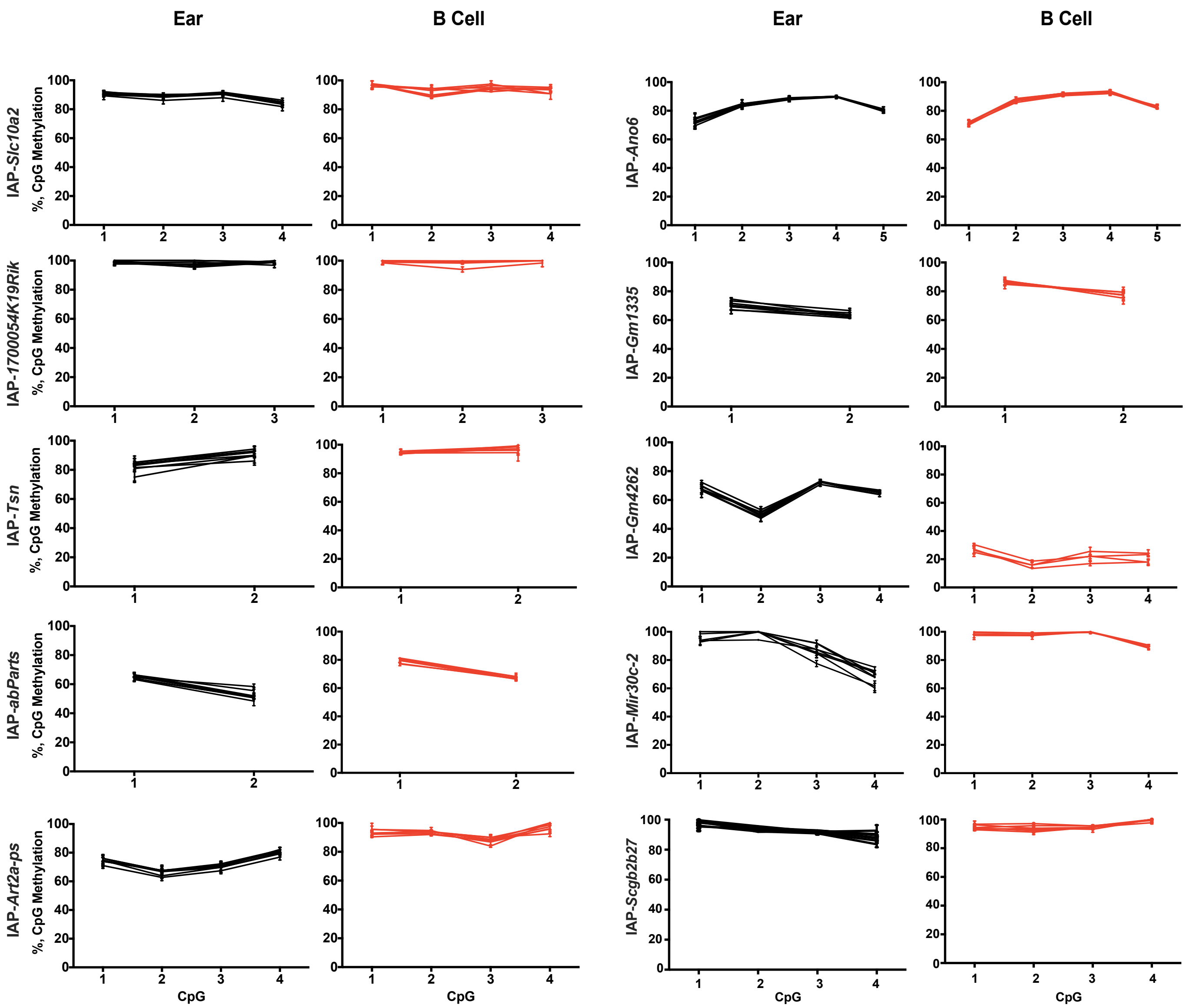

False positive IAP elements - T cells

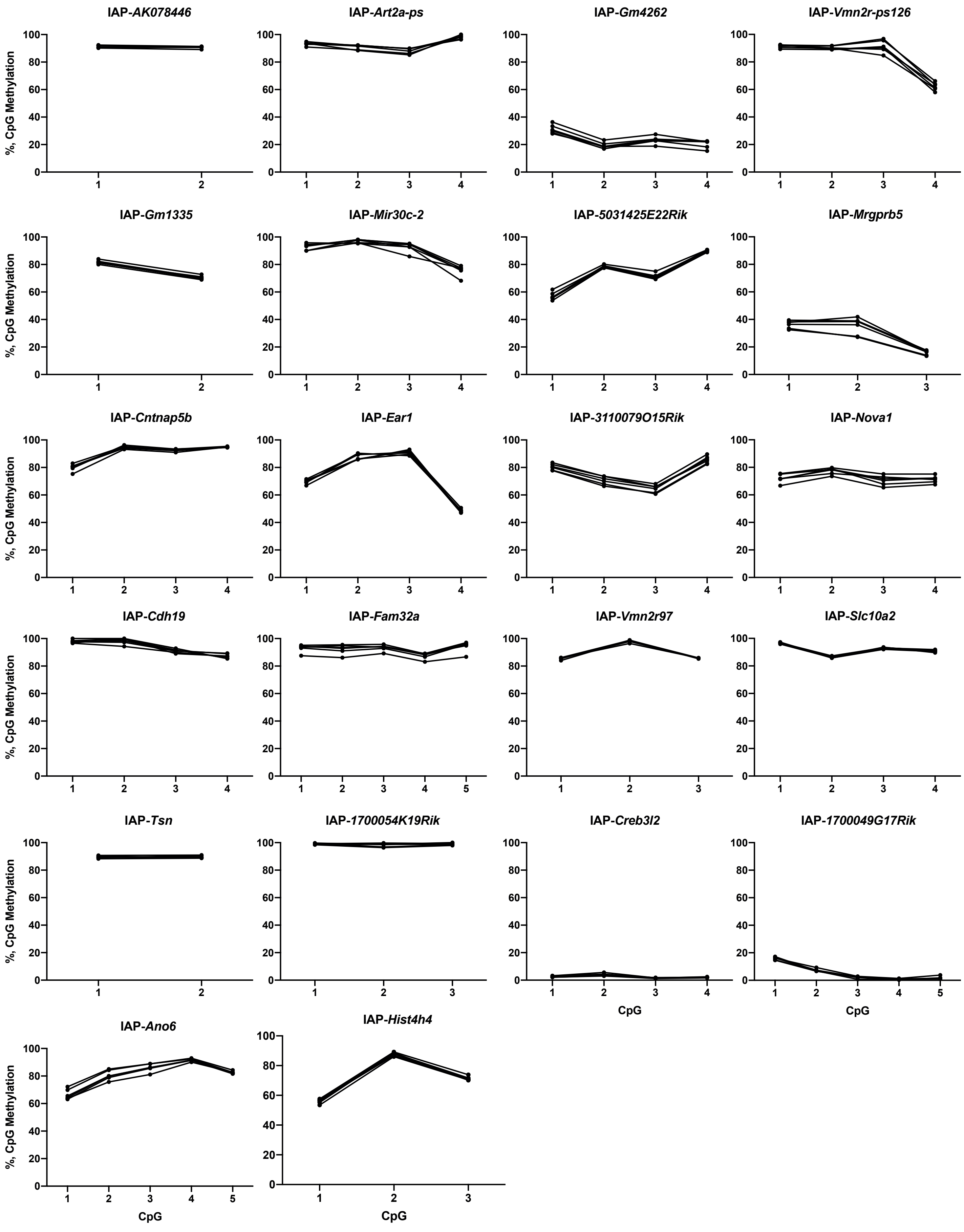
