## Supplemental Figures for "Genomic properties of variably methylated retrotransposons in mouse"

#### Supplemental Figure S1
**A**

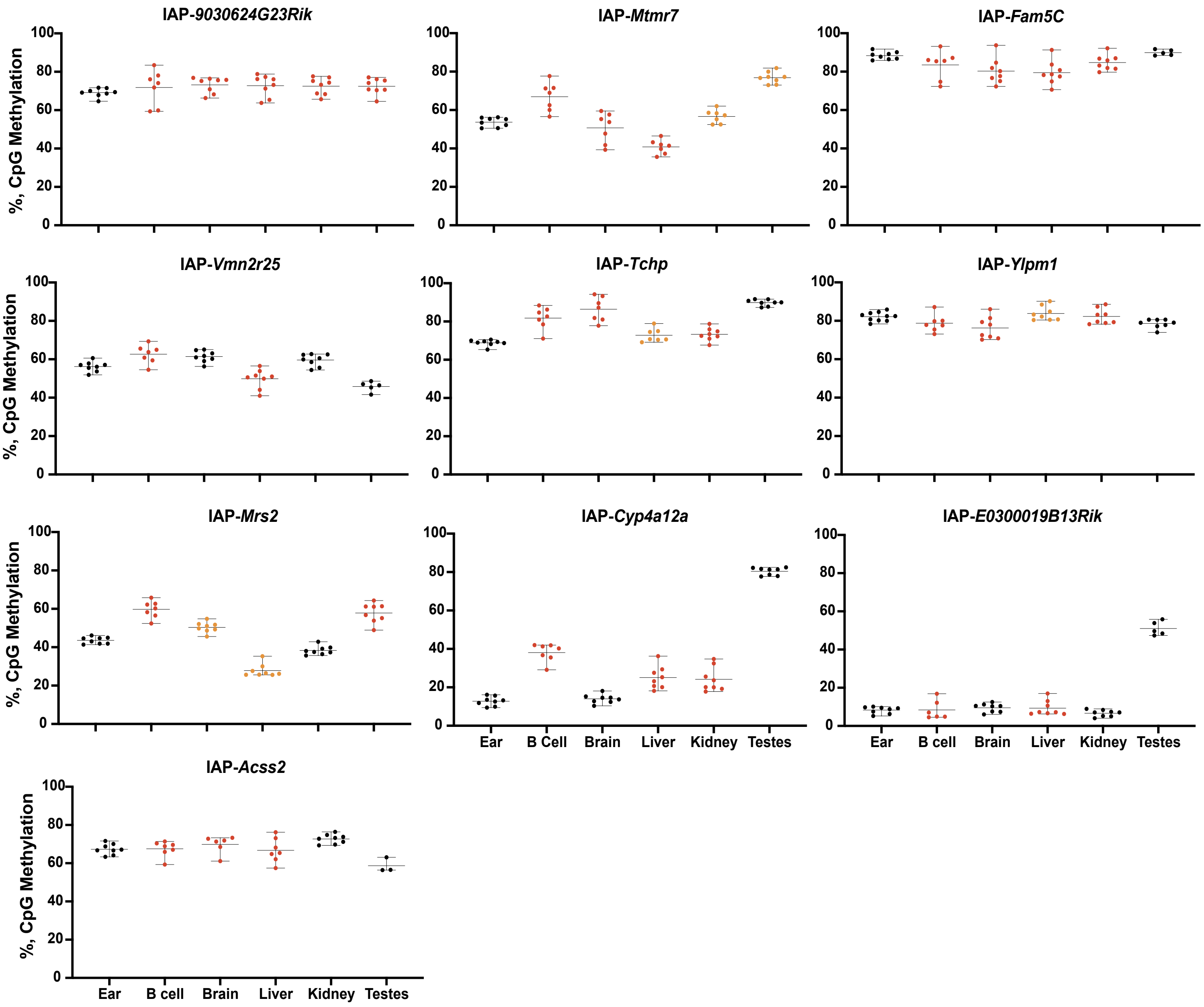

B

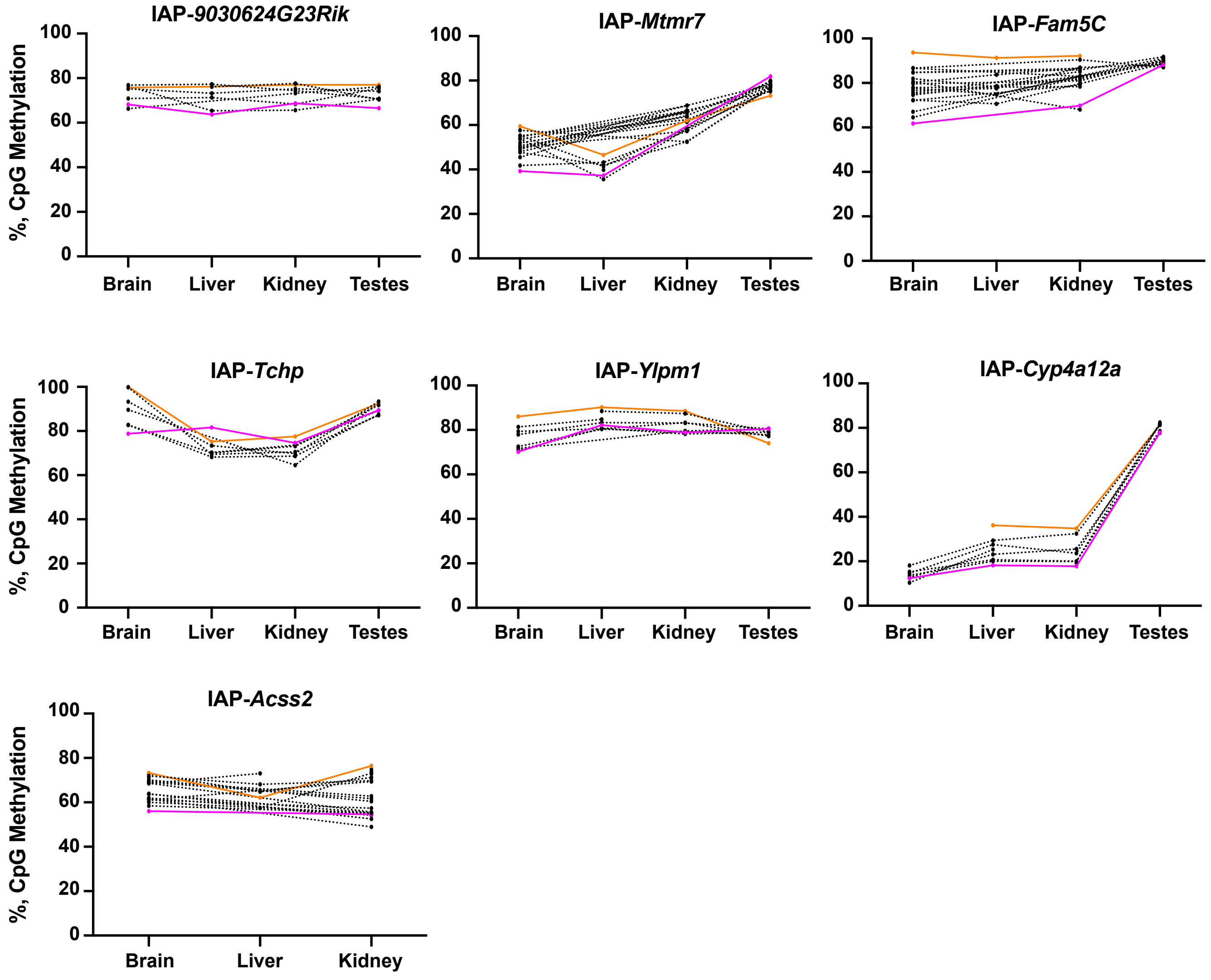

IAP-*9030624G23Rik*
IAP-*Mtmr7*
IAP-*Fam5C*
IAP-*Tchp*
IAP-*Ylpm1*
IAP-*Cyp4a12a*
IAP-*Acss2*
%, CpG Methylation
0
20
40
60
80
100
Brain
Liver
Kidney
Testes


C

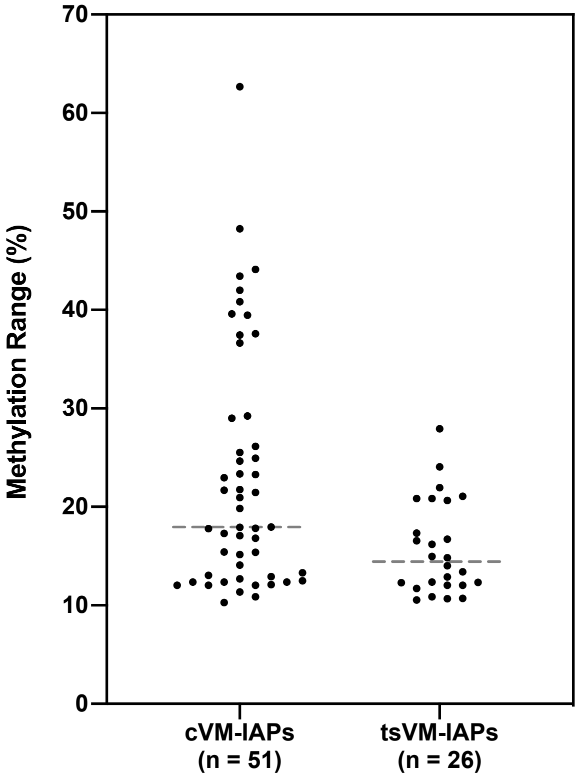

Methylation Range (%)
0
10
20
30
40
50
60
70
cVM-IAPs
(n = 51)
tsVM-IAPs
(n = 26)

D

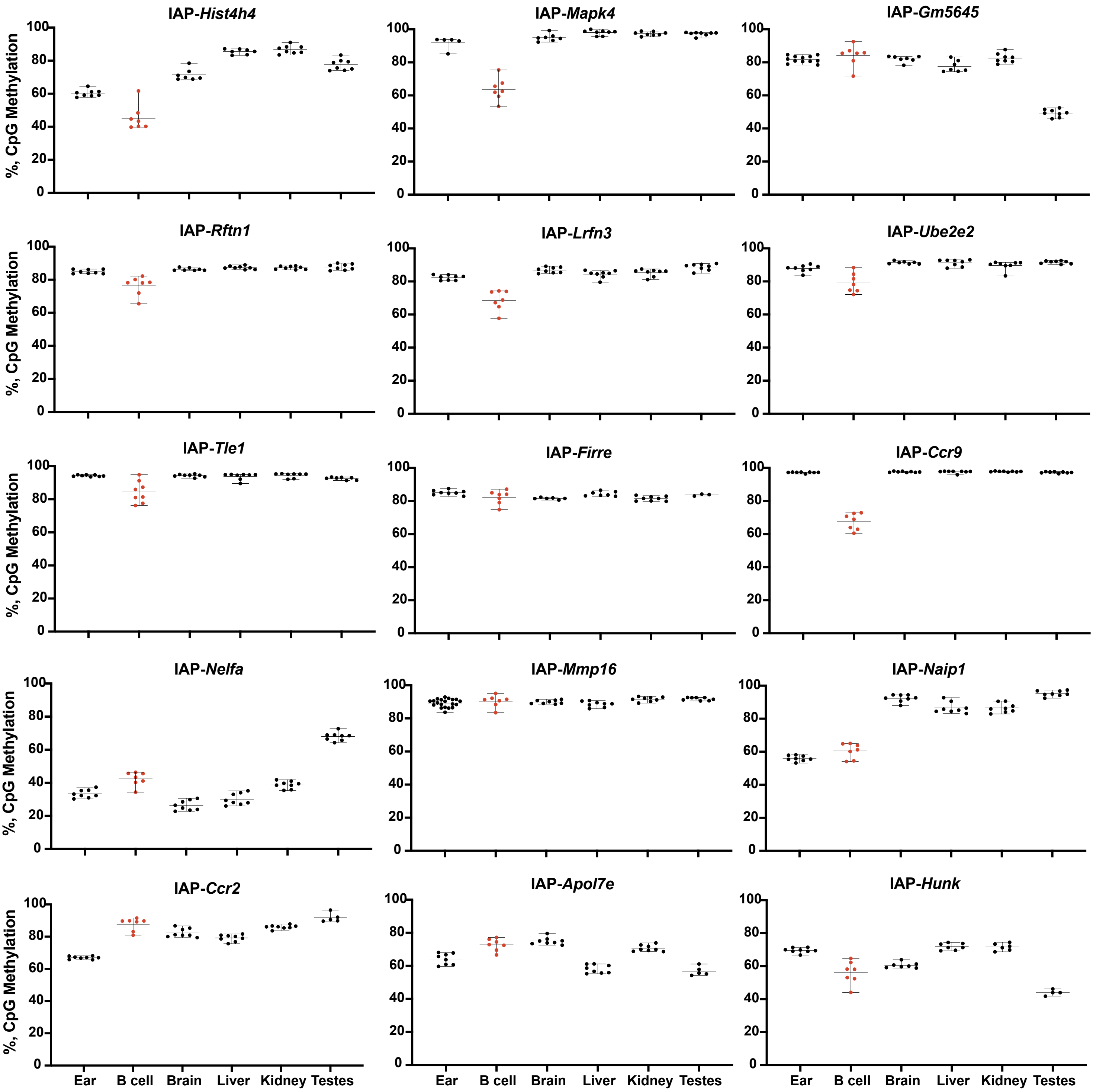
IAP-Hist4h4
IAP-Mapk4
IAP-Gm5645
IAP-Rftn1
IAP-Lrfn3
IAP-Ube2e2
IAP-Tle1
IAP-Firre
IAP-Ccr9
IAP-Nelfa
IAP-Mmp16
IAP-Naip1
IAP-Ccr2
IAP-Apol7e
IAP-Hunk
%, CpG Methylation
0
20
40
60
80
100
Ear
B cell
Brain
Liver
Kidney
Testes

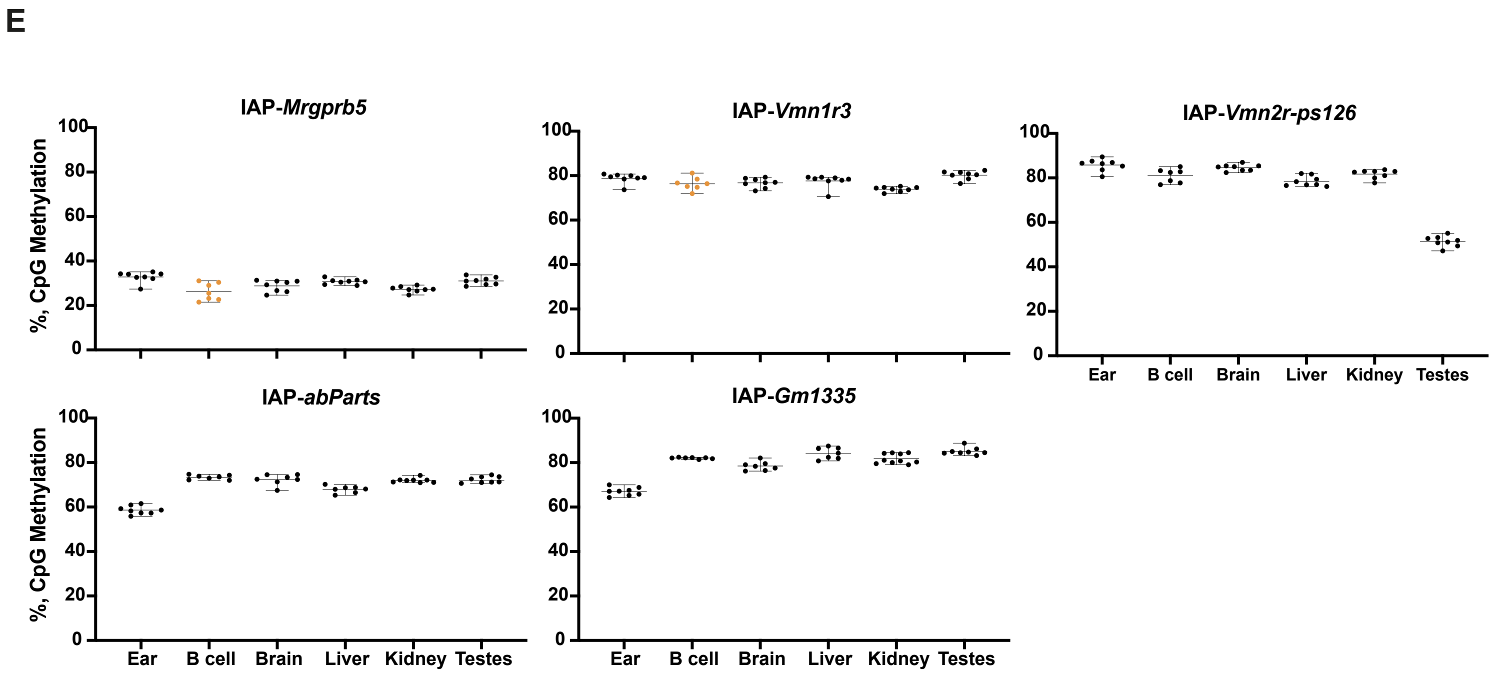

E
IAP-*Mrgprb5*
IAP-*Vmn1r3*
IAP-*Vmn2r-ps126*
IAP-*abParts*
IAP-*Gm1335*
%, CpG Methylation
0
20
40
60
80
100
Ear
B cell
Brain
Liver
Kidney
Testes

**Supplemental Figure S1. Categorisation of VM-IAPs.**

(A) Methylation level as assessed by bisulphite pyrosequencing of 10 tissue-specific VM-IAPs (tsVM-IAPs) that are variably methylated in multiple tissues. Each point represents an individual and is the average methylation level across the CpGs at the distal end of the IAP LTR. Tissues with a methylation range ≥10% and <9% are shown in red and black, respectively. Tissues with a methylation range ≥9% but <10% are shown in orange. Bars represent the median and range of methylation values between individuals.

(B) Tissue-specific VM-IAPs display methylation consistency across tissues within each individual at a given locus, similarly to cVM-IAPs. Each line represents an individual mouse; the individuals with the highest and lowest methylation in the most variable tissue of each IAP are highlighted for comparison.

(C) Methylation range of validated elements separated into constitutive and tissue-specific VM-IAPs. Each point represents an individual element. The methylation range in the most variable tissue is plotted for tsVM-IAPs. The dotted line represents the median methylation range.

(D) As in (A) for the 16 tsVM-IAPs with methylation variability restricted to B cells.

(E) As in (A) for five false positive IAPs with no methylation variability in any tissue assayed.

#### Supplemental Figure S2

**A**

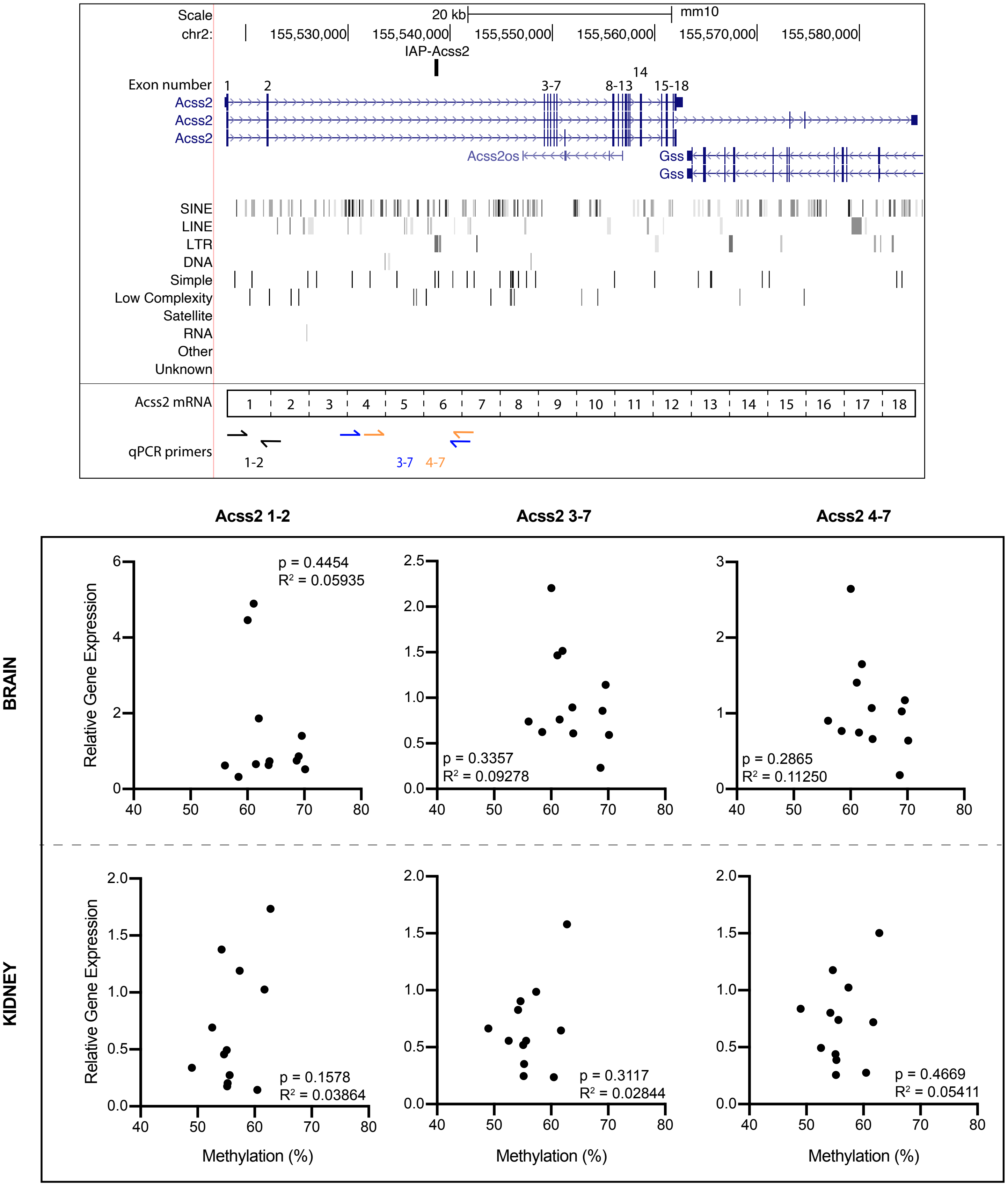

B

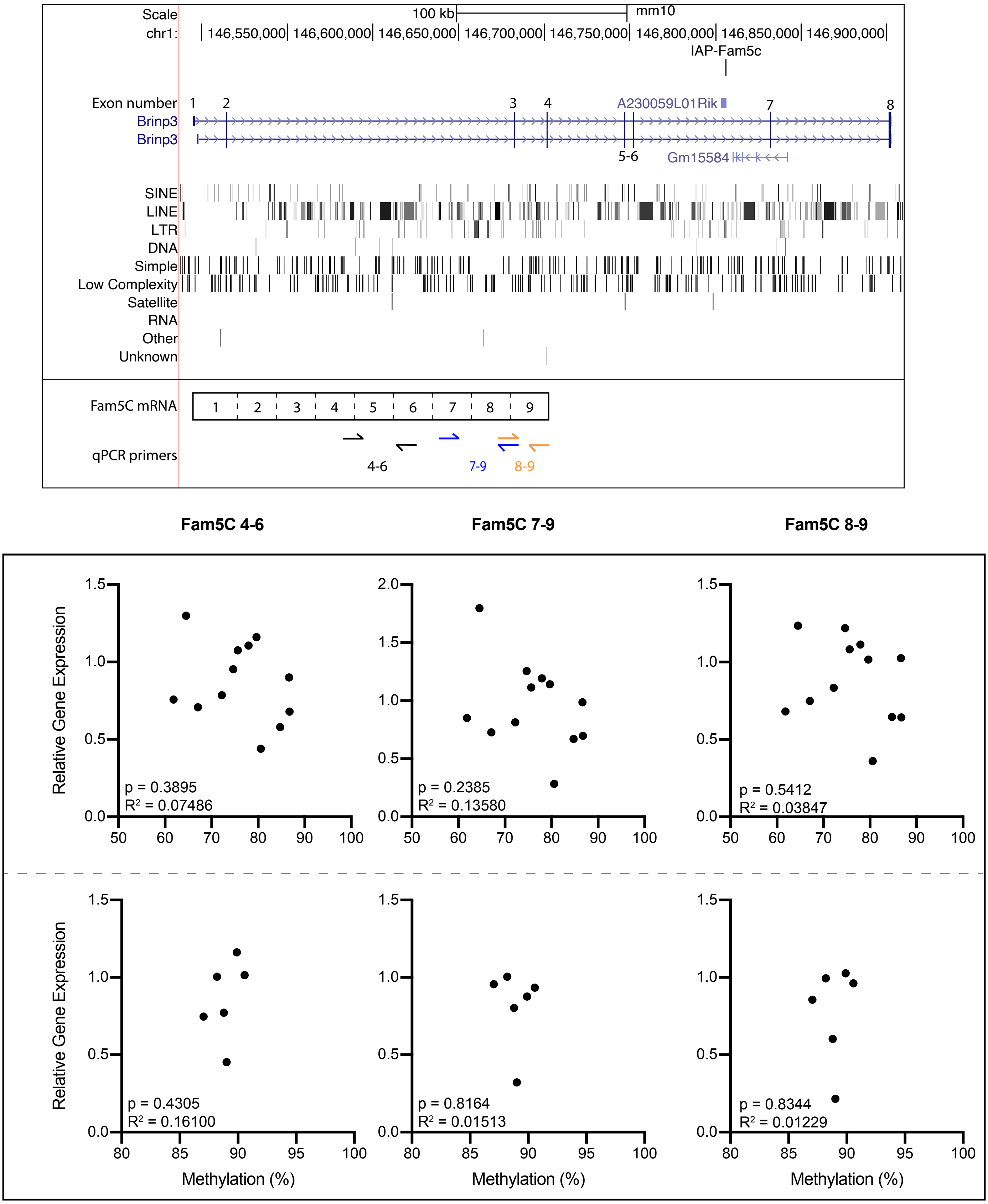

Scale
100 kb
mm10
chr1:
146,550,000
146,600,000
146,650,000
146,700,000
146,750,000
146,800,000
146,850,000
146,900,000
IAP-Fam5c
Exon number
Brinp3
Brinp3
A230059L01Rik
Gm15584
5-6
SINE
LINE
LTR
DNA
Simple
Low Complexity
Satellite
RNA
Other
Unknown
Fam5C mRNA
qPCR primers
4-6
7-9
8-9
Fam5C 4-6
Fam5C 7-9
Fam5C 8-9
BRAIN
TESTES
Relative Gene Expression
Methylation (%)
p = 0.3895
R² = 0.07486
p = 0.2385
R² = 0.13580
p = 0.5412
R² = 0.03847
p = 0.4305
R² = 0.16100
p = 0.8164
R² = 0.01513
p = 0.8344
R² = 0.01229

C

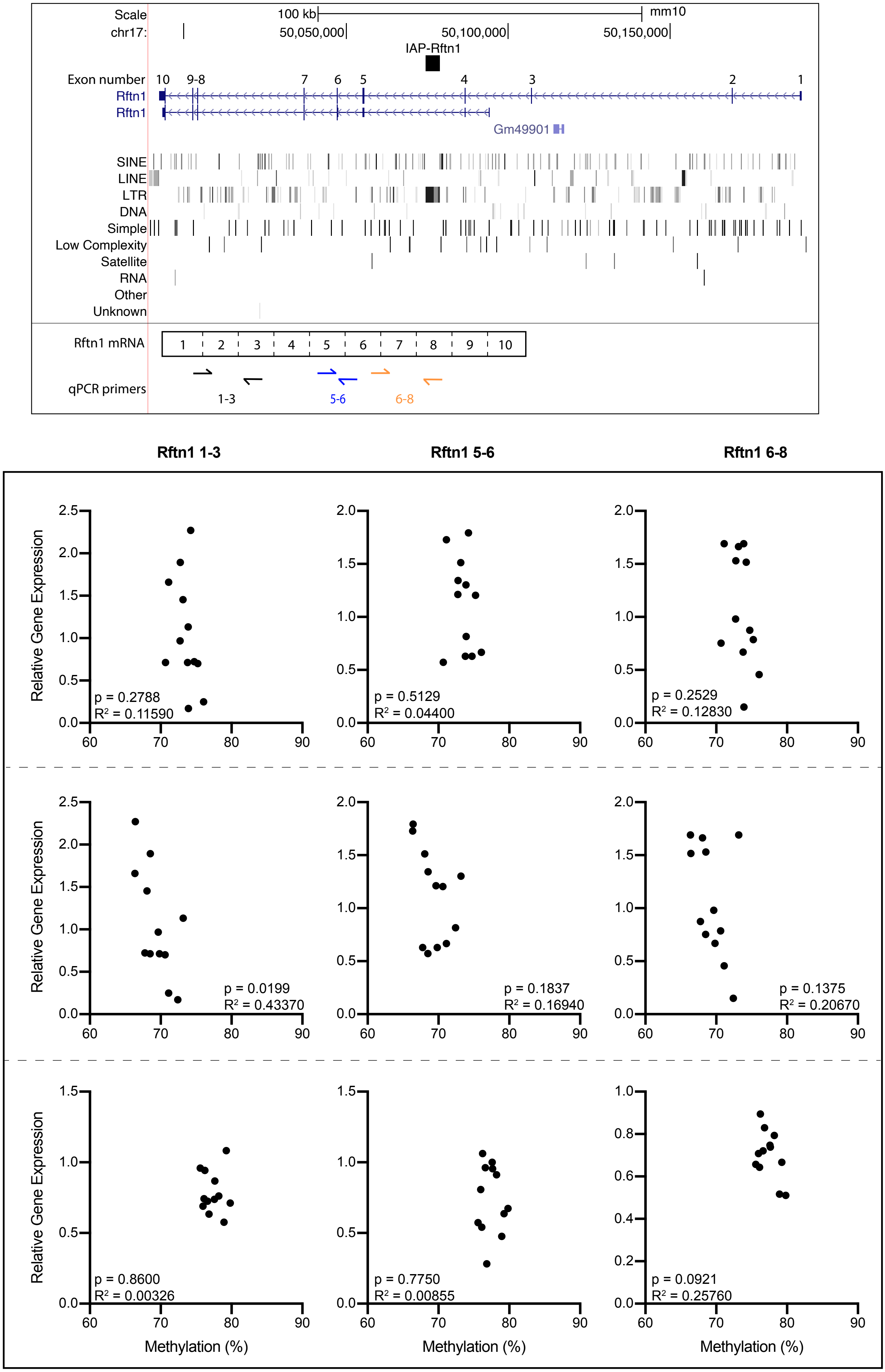
Scale
100 kb
mm10
chr17:
50,050,000
50,100,000
50,150,000
IAP-Rftn1
Exon number
10 9-8 7 6 5 4 3 2 1
Rftn1
Rftn1
Gm49901
SINE
LINE
LTR
DNA
Simple
Low Complexity
Satellite
RNA
Other
Unknown
Rftn1 mRNA
1 2 3 4 5 6 7 8 9 10
qPCR primers
1-3
5-6
6-8
Rftn1 1-3
Rftn1 5-6
Rftn1 6-8
BRAIN
B CELLS
T CELLS
Relative Gene Expression
Methylation (%)
p = 0.2788
R² = 0.11590
p = 0.5129
R² = 0.04400
p = 0.2529
R² = 0.12830
p = 0.0199
R² = 0.43370
p = 0.1837
R² = 0.16940
p = 0.1375
R² = 0.20670
p = 0.8600
R² = 0.00326
p = 0.7750
R² = 0.00855
p = 0.0921
R² = 0.25760

**D**

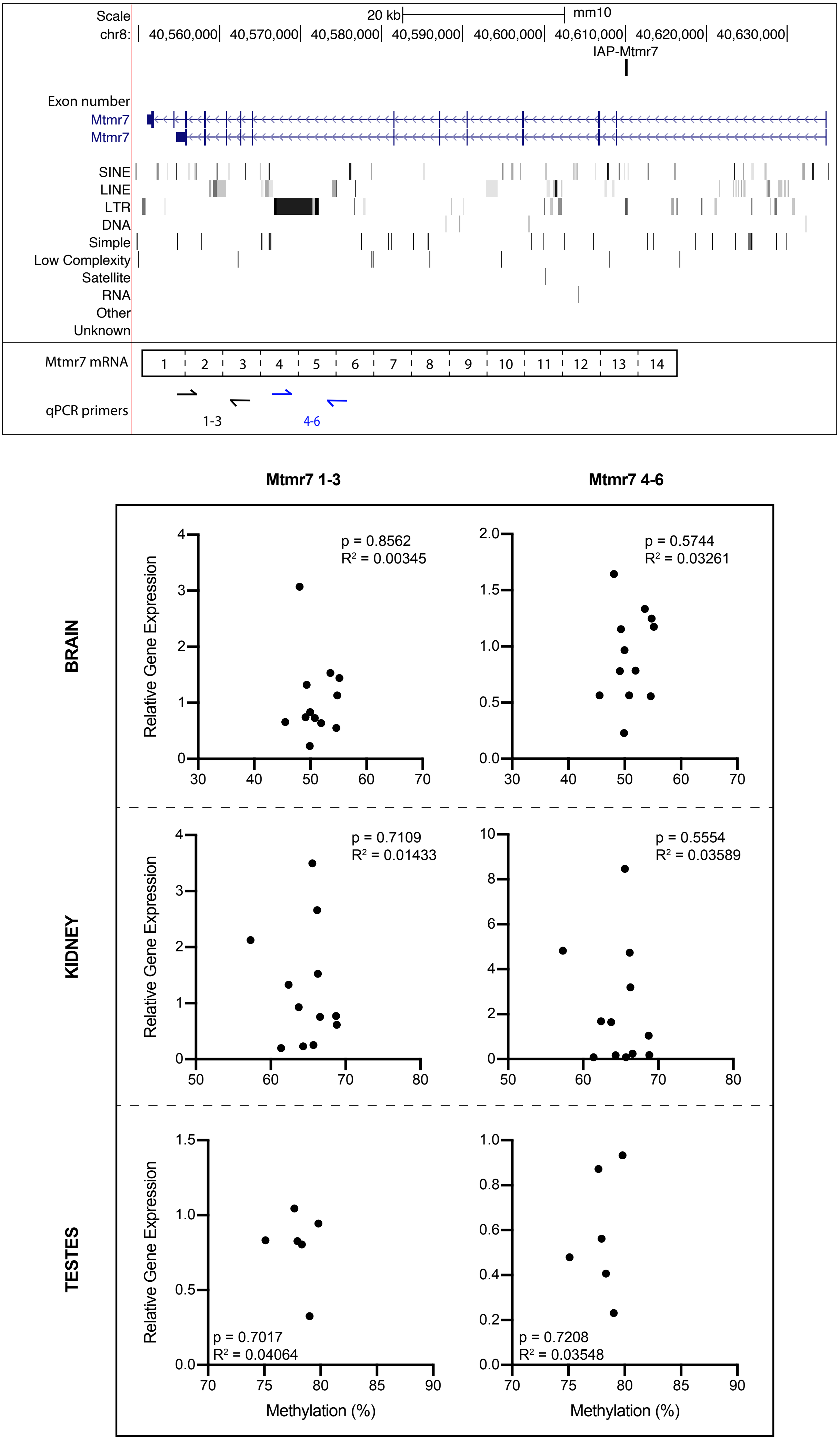

Scale
20 kb
mm10
chr8:
40,560,000
40,570,000
40,580,000
40,590,000
40,600,000
40,610,000
40,620,000
40,630,000
IAP-Mtmr7
Exon number
Mtmr7
Mtmr7
SINE
LINE
LTR
DNA
Simple
Low Complexity
Satellite
RNA
Other
Unknown
Mtmr7 mRNA
1 2 3 4 5 6 7 8 9 10 11 12 13 14
qPCR primers
1-3
4-6
Mtmr7 1-3
Mtmr7 4-6
BRAIN
KIDNEY
TESTES
Relative Gene Expression
p = 0.8562
R² = 0.00345
p = 0.5744
R² = 0.03261
p = 0.7109
R² = 0.01433
p = 0.5554
R² = 0.03589
p = 0.7017
R² = 0.04064
p = 0.7208
R² = 0.03548
Methylation (%)

**Supplemental Figure S2. Tissue-specific VM-IAPs rarely affect gene expression.**

Screenshots of the UCSC genome browser in each top panel showing the position of the tsVM-IAP within the gene selected for expression analysis, as well as a diagram of locations of the qPCR primers relative to the exons. The relationship between gene expression and methylation of the tsVM-IAP is shown in each bottom panel. tsVM-IAPs shown are (A) IAP-*Acss2*, (B) IAP-*Fam5C*, (C) IAP-*Rftn1*, and (D) IAP-*Mtmr7*. Expression was quantified by qPCR, with input cDNA normalised to two housekeeping genes, *Pgk1* and *Gapdh*; p-values are from two-tailed Pearson test. n = 6 for testes and 12 for all other tissues.

#### Supplemental Figure S3

**A**

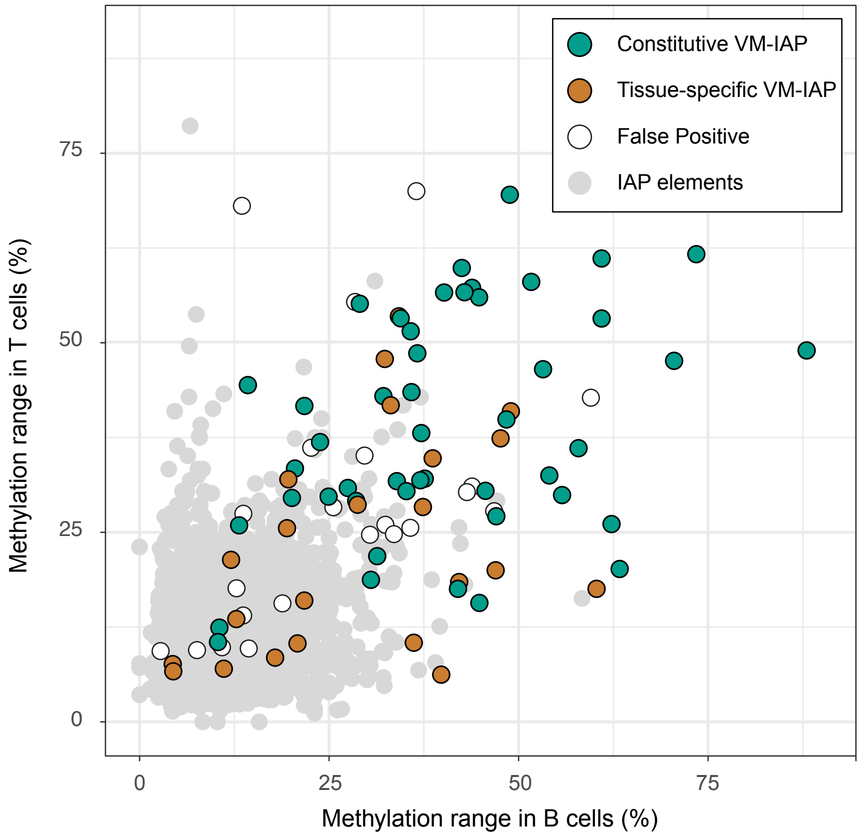


**B**

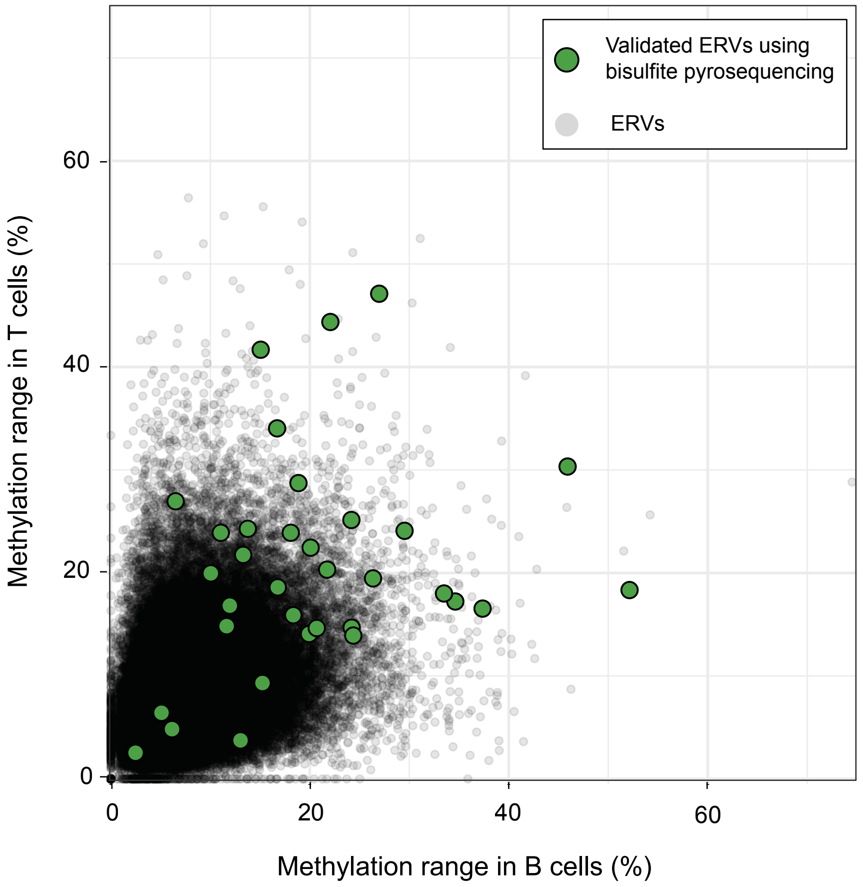

C

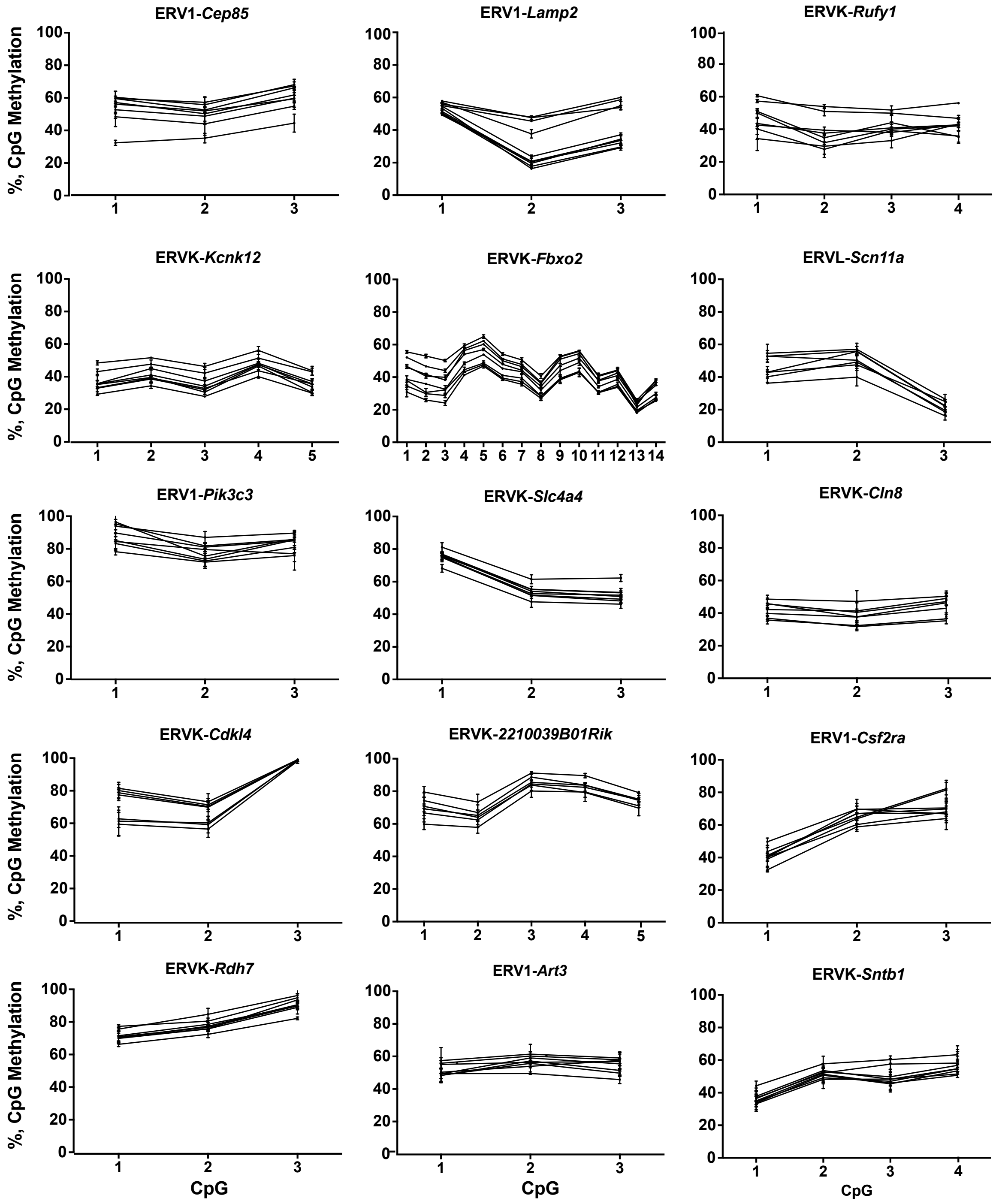
ERV1-Cep85
ERV1-Lamp2
ERVK-Rufy1
ERVK-Kcnk12
ERVK-Fbxo2
ERVL-Scn11a
ERV1-Pik3c3
ERVK-Slc4a4
ERVK-Cln8
ERVK-Cdkl4
ERVK-2210039B01Rik
ERV1-Csf2ra
ERVK-Rdh7
ERV1-Art3
ERVK-Sntb1
%, CpG Methylation
CpG

D

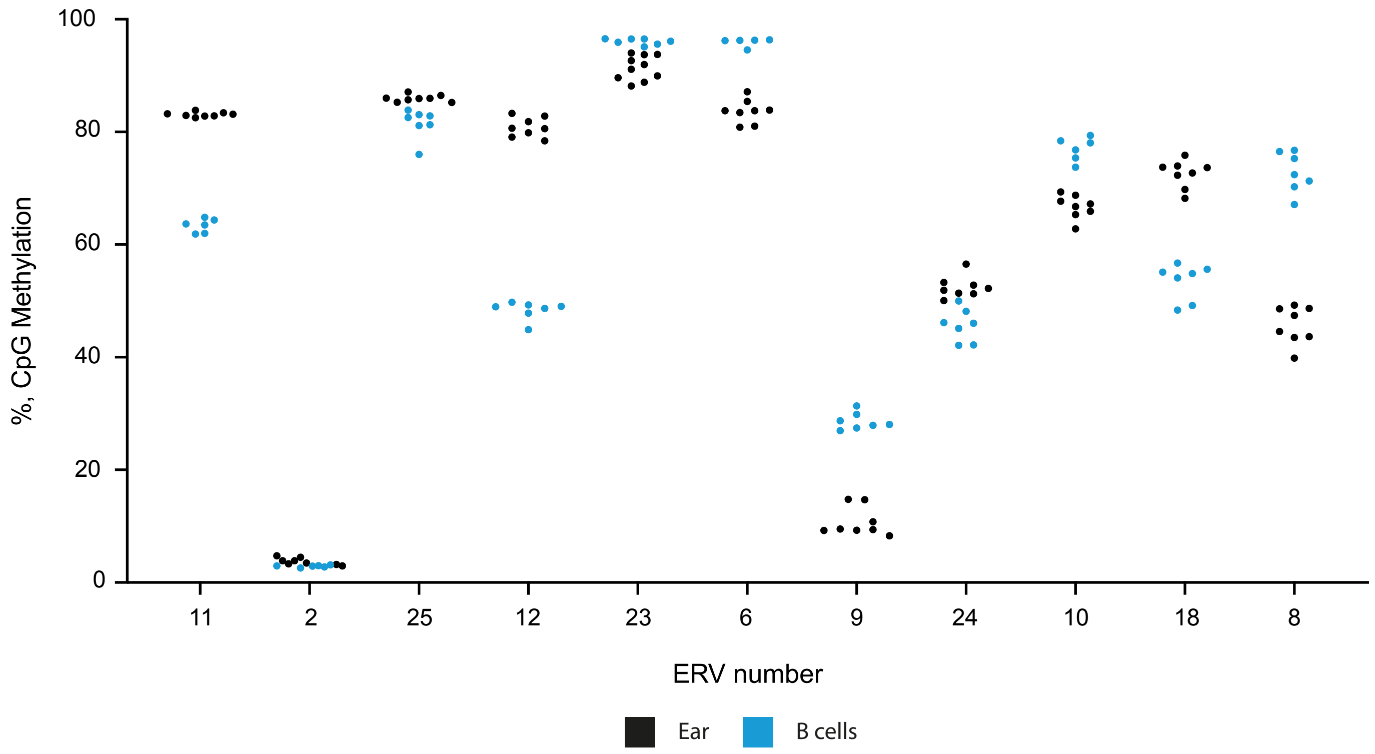

100
80
60
40
20
0
%, CpG Methylation
11
2
25
12
23
6
9
24
10
18
8
ERV number
Ear
B cells

E

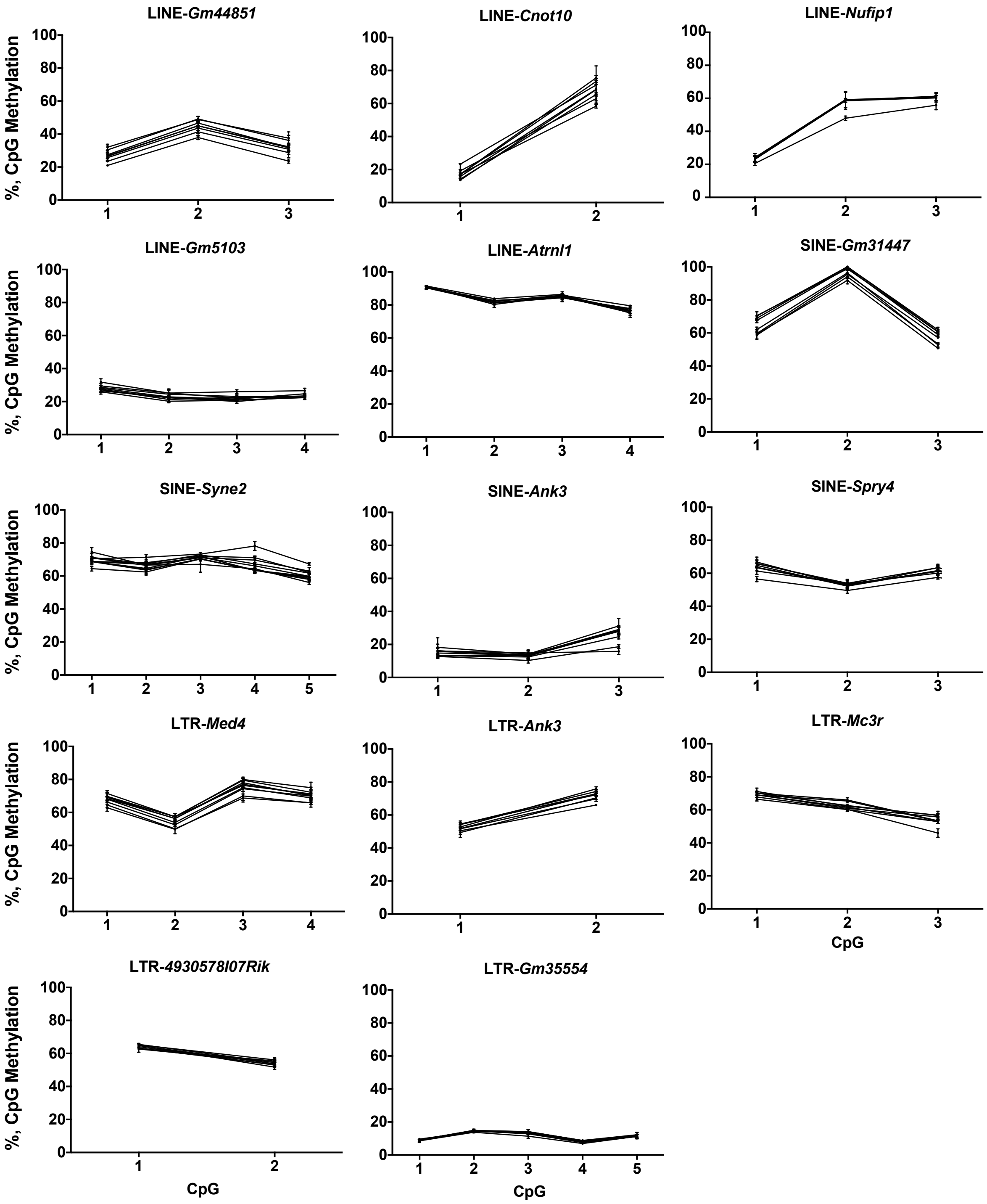
LINE-*Gm44851*
LINE-*Cnot10*
LINE-*Nufip1*
LINE-*Gm5103*
LINE-*Atrnl1*
SINE-*Gm31447*
SINE-*Syne2*
SINE-*Ank3*
SINE-*Spry4*
LTR-*Med4*
LTR-*Ank3*
LTR-*Mc3r*
LTR-*4930578I07Rik*
LTR-*Gm35554*
%, CpG Methylation
CpG

F

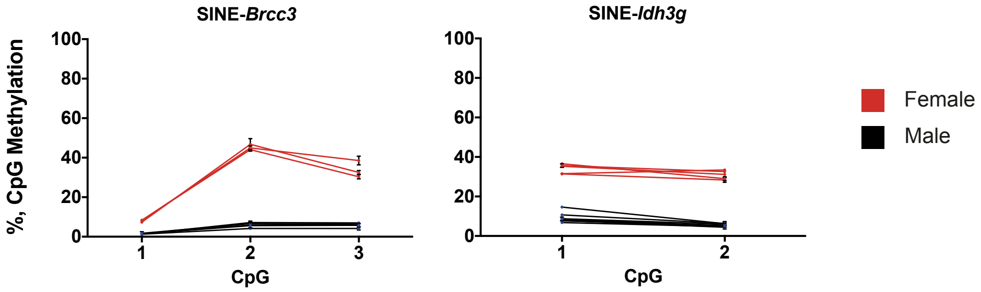
SINE-*Brcc3*
SINE-*Idh3g*
%, CpG Methylation
CpG
Female
Male

**Supplemental Figure S3. Results and validations of screens for variable methylation in retrotransposons.**

(A-B) Methylation ranges in B and T cell WGBS data for (A) IAP elements and (B) ERVs. Each point represents an individual element: cVM-IAPs, tsVM-IAPs and false positives are coloured in (A); validated ERVs (15 VM-ERVs and 18 false positives) are coloured in (B).

(C) Bisulfite pyrosequencing of candidate ERVs assayed in ear samples. Only those passing the ≥10% methylation range threshold are shown. Each line represents an individual mouse. Samples were run in triplicate; the error bar shows the standard deviation of technical replicates. CpGs are assayed from the distal end of the ERV.

(D) Methylation level of individual ERVs which did not cross the ≥10% methylation range threshold, assayed in ear and B cells. Each point represents an individual and is the average methylation level across the CpGs located at the distal end of the ERV. Ear and B cell samples are shown in black and blue, respectively. 'ERV numbers' are shown, as gene-based identifiers were only generated for VM-ERVs.

(E) Same as panel (C), showing pyrosequencing of LINE, SINE, and non-ERV LTR candidates. All tested elements are shown (except the two SINEs in *(*F*)*), including those that did not pass the ≥10% methylation range threshold.

(F) Two tested SINEs are bistable epialleles due to their location on the X chromosome. Female and male individuals are coloured in red and black, respectively.

#### Supplemental Figure S4

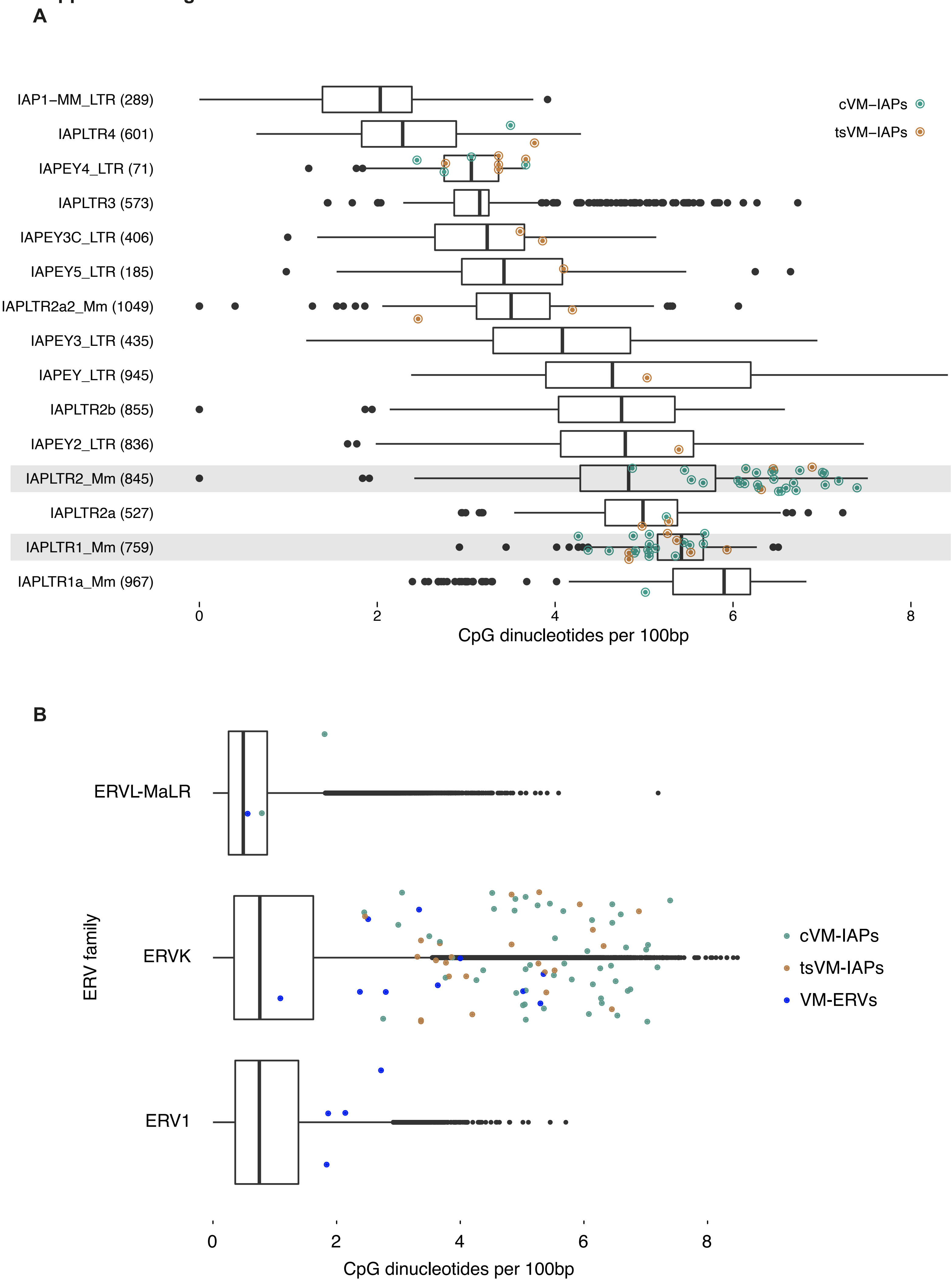

**Supplemental Figure S4. Increased CpG density in IAPLTR2_Mm cVM-IAPs.**

(A) The two IAP LTR types over-represented in cVM-IAPs, LTR1_Mm and LTR2_Mm, have higher average CpG density compared to all IAP LTRs (means: LTR1_Mm=5.496, LTR2_Mm=4.993, all IAP LTRs=4.345; p<2.2e-16 for both). LTR1_Mm cVM-IAPs have significantly lower CpG density compared to non-variable LTR1_Mm elements (difference of means=-0.348, Cohen's d=-0.891, p<0.003), while LTR2_Mm cVM-IAPs have significantly higher CpG density compared to non-variable LTR2_Mm elements (difference of means=1.44, Cohen's d=1.431, p<1.3e-12). The effect size measured by Cohen's d is much larger for LTR2_Mm elements than LTR1_Mm elements. tsVM-IAPs are not enriched for a particular LTR type and do not show any difference in CpG density compared to all IAP LTRs (p= 0.18). The number of elements analysed per IAP LTR type is shown in parentheses to the right of the IAP LTR type names. CpG density was calculated by normalising the number of CpGs to the LTR length in base pairs for each IAP LTR; p-values are from Welch's t test. Cohen's d is an unbiased measure of effect size. Analyses only included LTRs; for fully-structured IAPs, only the 5' LTR was used.

(B) VM-ERVs have higher average CpG density compared to all other ERVs (means: VM-ERVs=2.968, all ERVs=0.804; p=5.99e-5), but lower average CpG density compared to cVM-IAPs and tsVM-IAPs (means: cVM-IAPs=5.259, tsVM-IAPs=4.628). CpG density was calculated by normalising the number of CpGs to the LTR length in base pairs for each ERV LTR; p-values are from Welch's t test.

#### Supplemental Figure S5

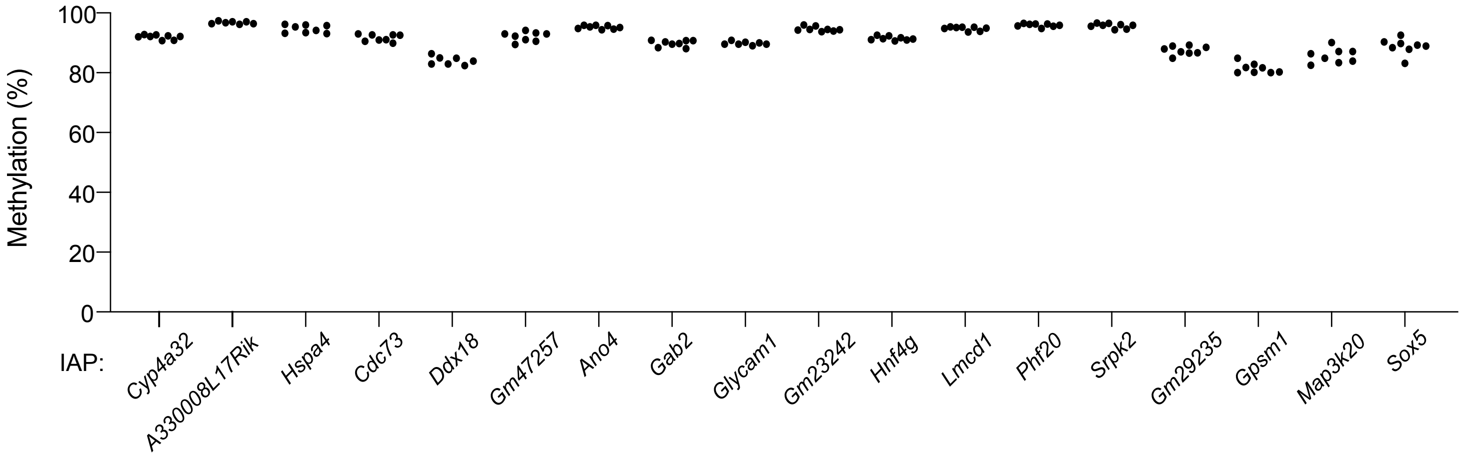

**Supplemental Figure S5. 18 IAP elements containing sequences over-represented in cVM-IAPs are hypermethylated.**

Each point represents an individual mouse and is the average methylation level across the CpGs at the distal end of the IAP LTR, n = 8.

**Supplemental Figure S6**

**A**

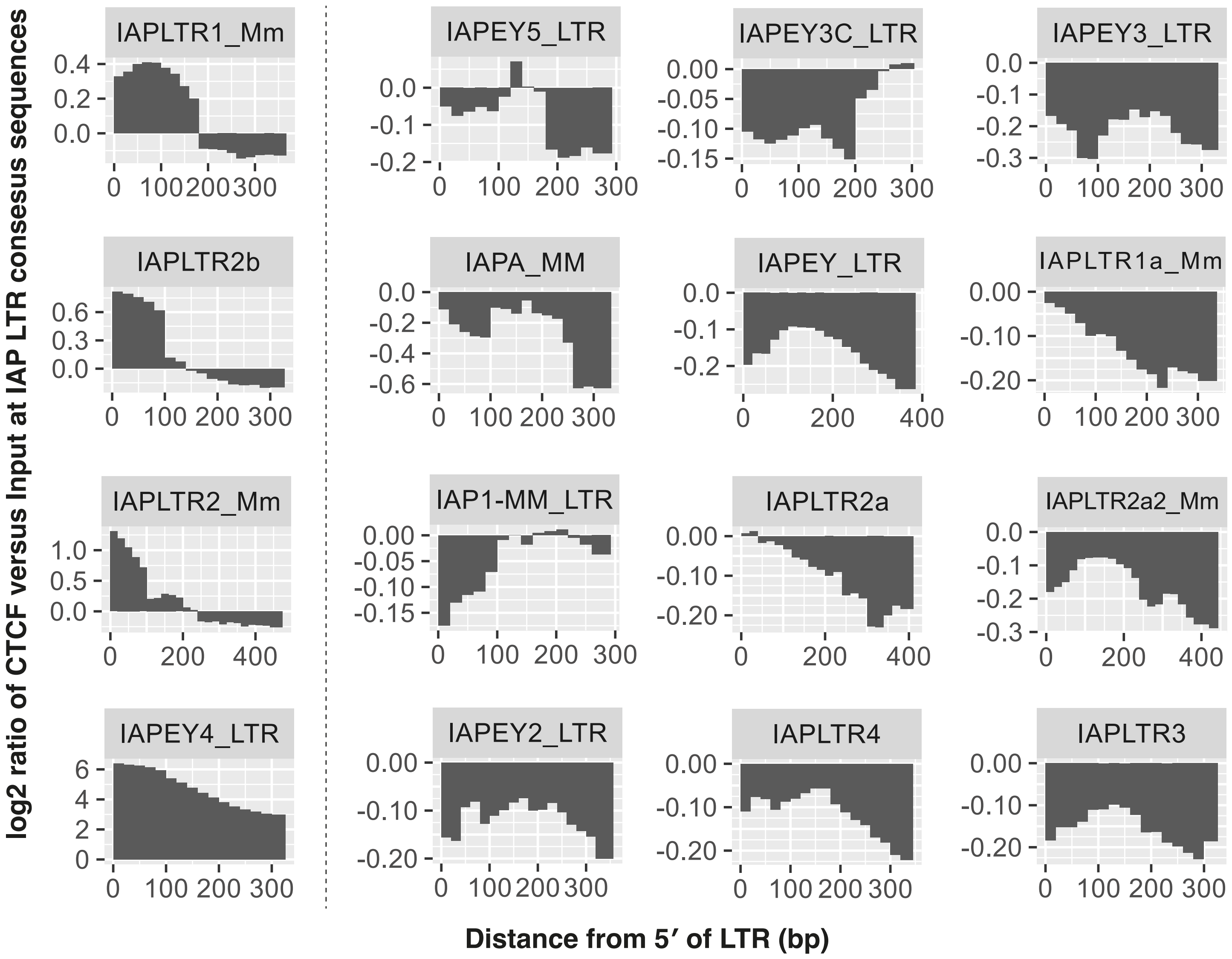


**B**

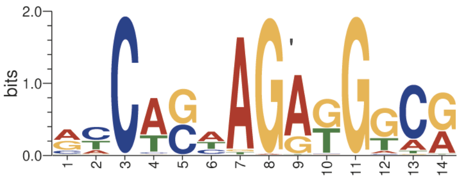


CTCF binding site in IAP LTRs

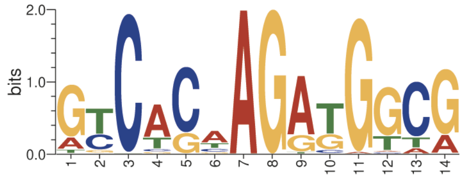


CTCF binding site in constitutive VM-IAP LTRs

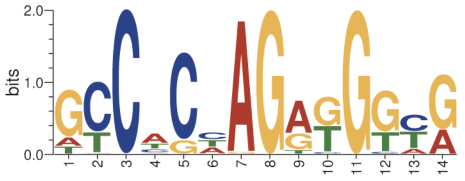


CTCF binding site in tissue-specific VM-IAP LTRs

**Supplemental Figure S6. CTCF preferentially binds at a subset of IAP LTR types.**

(A) CTCF ChIP-seq data mapped to consensus sequences of each type of IAP LTR. CTCF appears to bind at the LTR types left of the dashed line, including the two types for which there is enrichment in cVM-IAPs, and not at other types (right of the dashed line).

(B) Motifs generated from CTCF binding sites located within LTRs of all IAPs, cVM-IAPs, and tsVM-IAPs. CTCF binding sites at these regions were originally identified by mapping the motif generated in Figure 3A to the genome.

#### Supplemental Figure S7

**A**

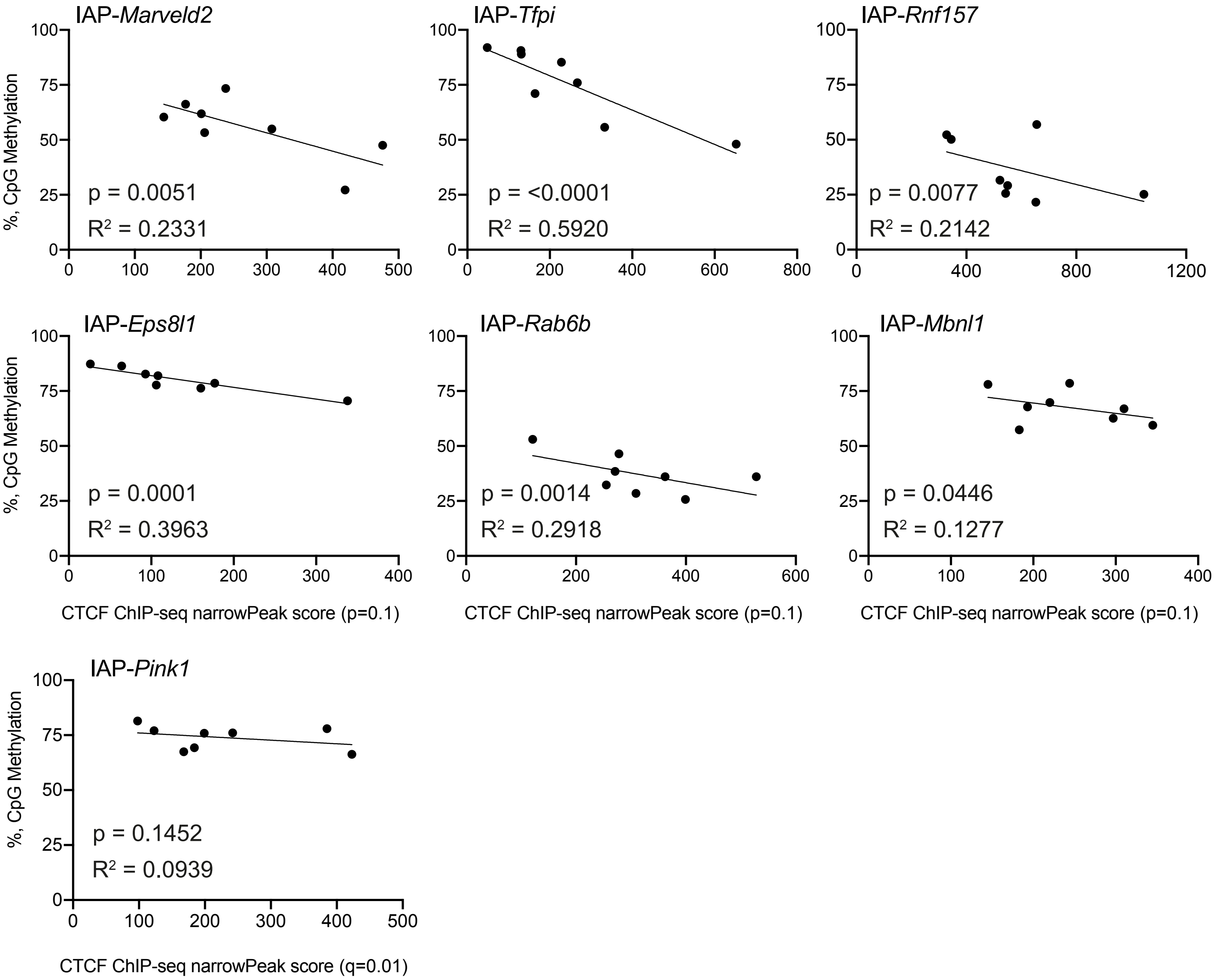

**B**

**IAP-*Tfpi***

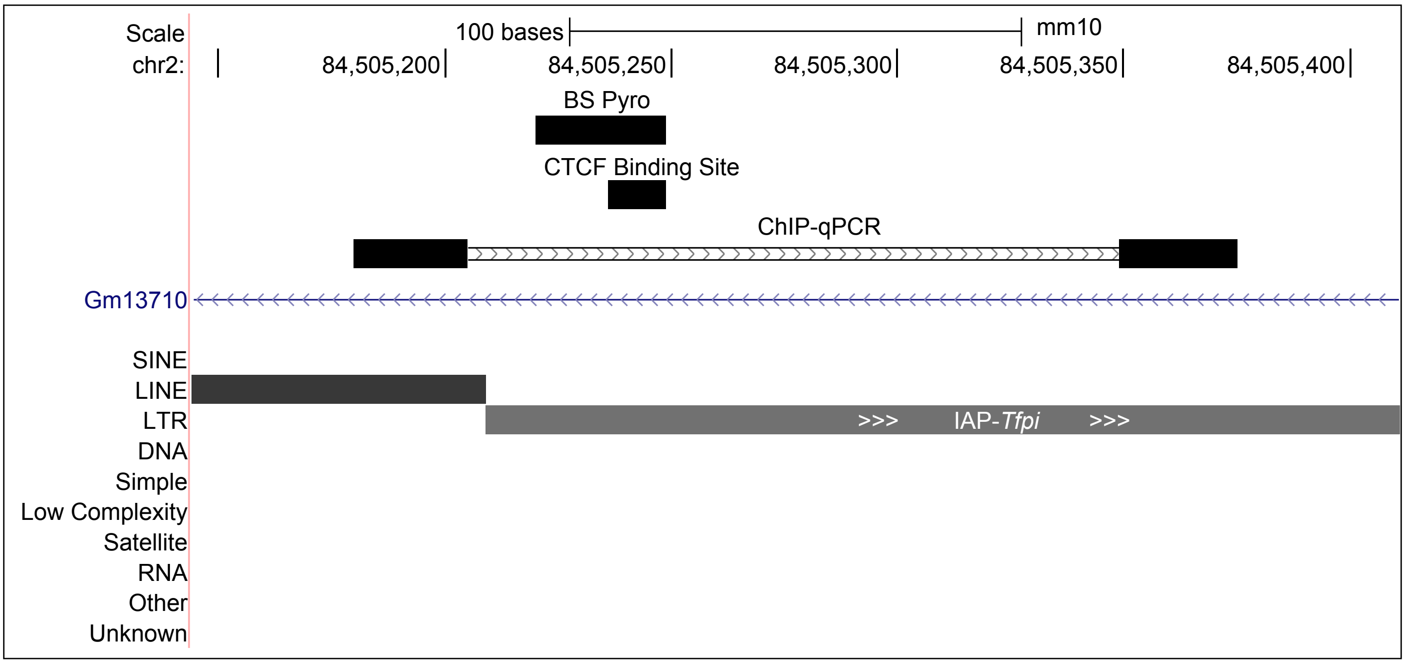


**IAP-*Marveld2***

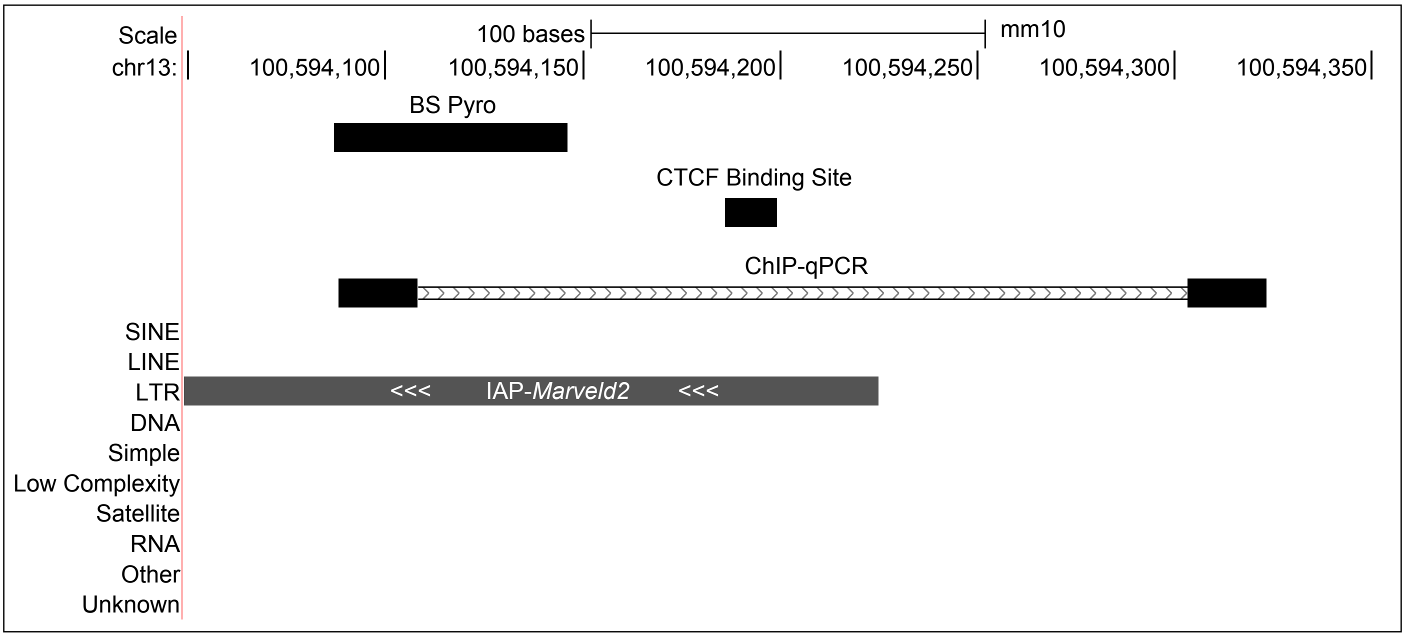

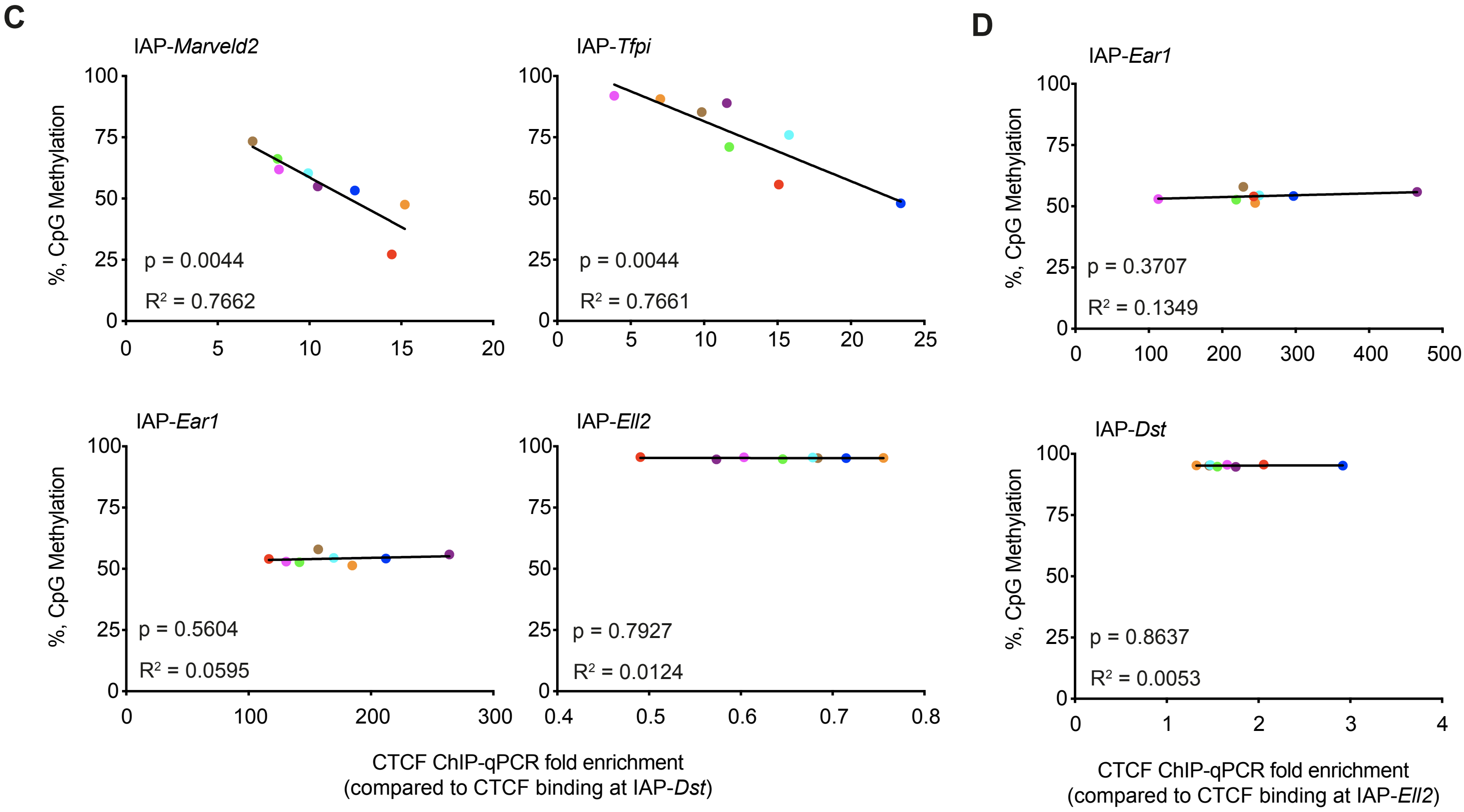
C
IAP-Marveld2
IAP-Tfpi
IAP-Ear1
IAP-Ell2
%, CpG Methylation
p = 0.0044
R² = 0.7662
p = 0.0044
R² = 0.7661
p = 0.5604
R² = 0.0595
p = 0.7927
R² = 0.0124
CTCF ChIP-qPCR fold enrichment
(compared to CTCF binding at IAP-Dst)
D
IAP-Ear1
IAP-Dst
p = 0.3707
R² = 0.1349
p = 0.8637
R² = 0.0053
CTCF ChIP-qPCR fold enrichment
(compared to CTCF binding at IAP-Ell2)

**Supplemental Figure S7. Methylation at many cVM-IAPs correlates inversely with CTCF binding.**

(A) There is an inverse relationship between CTCF ChIP-seq narrowPeak scores and DNA methylation (measured by bisulfite pyrosequencing) in eight liver samples at six out of seven assayed cVM-IAPs: IAP-*Marveld2,* IAP-*Tfpi*, IAP-*Rnf157*, IAP-*Eps8l1*, IAP-*Rab6b*, and IAP-*Mbnl1*; inverse correlation was not observed for IAP-*Pink1*. NarrowPeak scores were generating using a p-value of 0.1 to ensure that peaks were called in all individuals in regions with low CTCF occupancy. The correlations are consistent using a q-value of 0.01 (data only shown for IAP-*Pink1*), but at some loci, peaks were unable to be called for all individuals if using this stricter threshold. Each point represents the average methylation of the three to four distal CpGs in the LTR of an individual. P-values of the correlations are from two-tailed Pearson test.

(B) Screenshots of the UCSC genome browser showing the span of the CpGs assayed by bisulfite pyrosequencing (BS Pyro), the CTCF binding site, and the ChIP-qPCR assay (primers represented by boxes) relative to the 5′ edges of the cVM-IAPs, IAP-*Tfpi* and IAP-*Marveld2*.

(C-D) CTCF ChIP-qPCR fold enrichment versus DNA methylation at several IAPs in the same eight individual liver samples as in (A). Each coloured point represents the average methylation of the four distal CpGs in the LTR of an individual. The colours represent the same individuals across all plots (including those in Fig. 4) and show that the relative order of CTCF binding is not the same in each element. P-values are from two-tailed Pearson test. (C) CTCF ChIP-qPCR fold enrichment calculated relative to IAP-*Dst* at cVM-IAPs, IAP-*Marveld2* and IAP-*Tfpi*, and non-variably methylated IAPs, IAP-*Ear1* and IAP-*Ell2*. (D) There is no significant correlation between CTCF ChIP-qPCR and DNA methylation at the non-variably methylated IAPs, IAP-*Ear1* and IAP-*Dst.*

#### Supplemental Figure S8

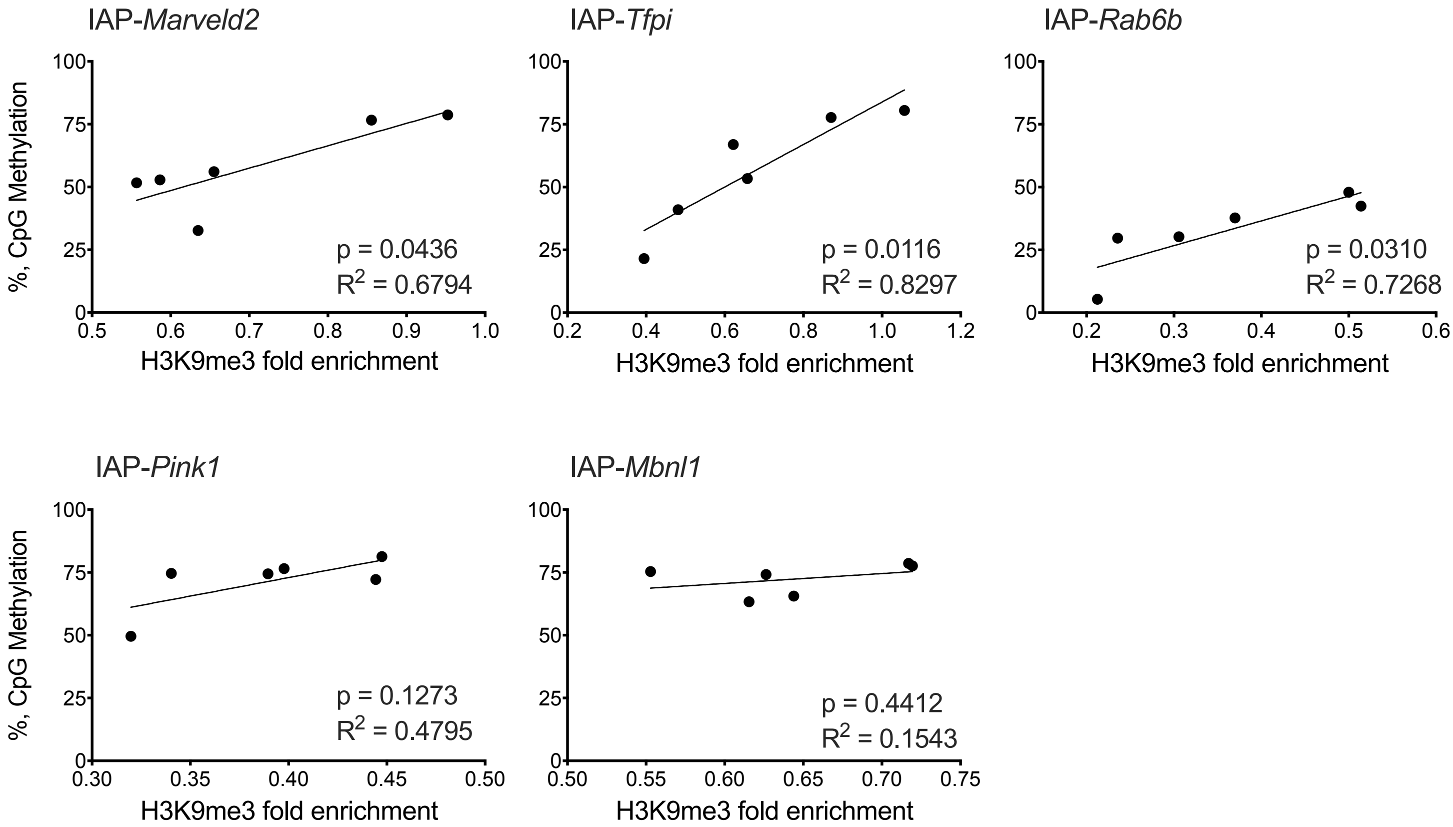

**Supplemental Figure S8. cVM-IAP methylation levels correlate with H3K9me3 enrichment.**

H3K9me3 ChIP-qPCR fold enrichment (relative to H3K9me3 enrichment at non-variable IAP-*Asxl3*) is correlated with DNA methylation at IAP-*Tfpi*, IAP-*Marveld2*, and IAP-*Rab6b* in adult liver. IAP-*Pink1* and IAP-*Mbnl1* do not exhibit significant correlations. Each point represents the average methylation level of the three or four distal CpGs in the 5′ LTR of an individual. P-values are from two-tailed Pearson test.

#### Supplemental Figure S9

**IAP-*Marveld2* (12.8% - 59.1% methylation range)**

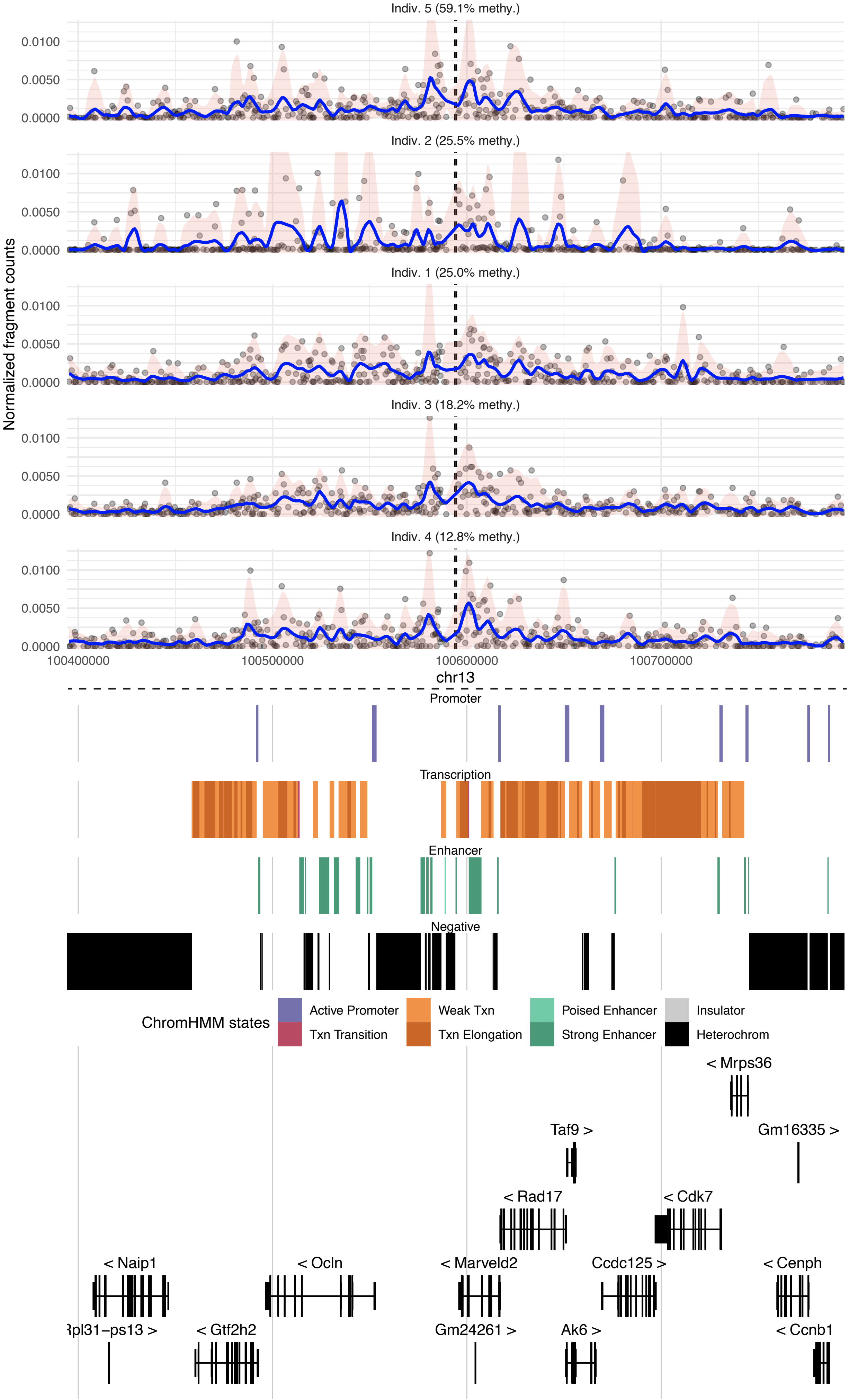

### IAP-*Tfec* (24.4% - 50.3% methylation range)

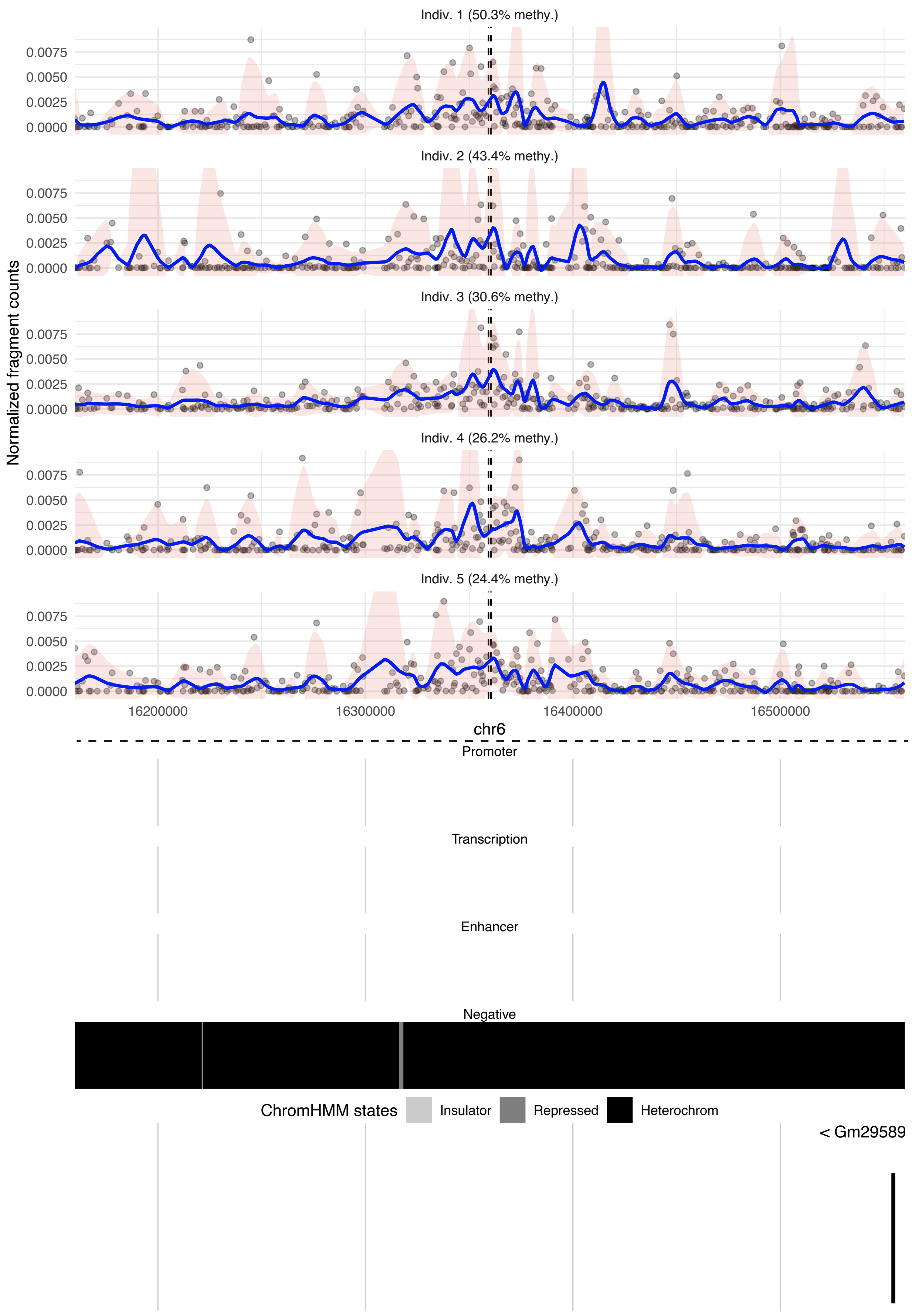

### IAP-*Pink1* (73.4% - 86.0% methylation range)

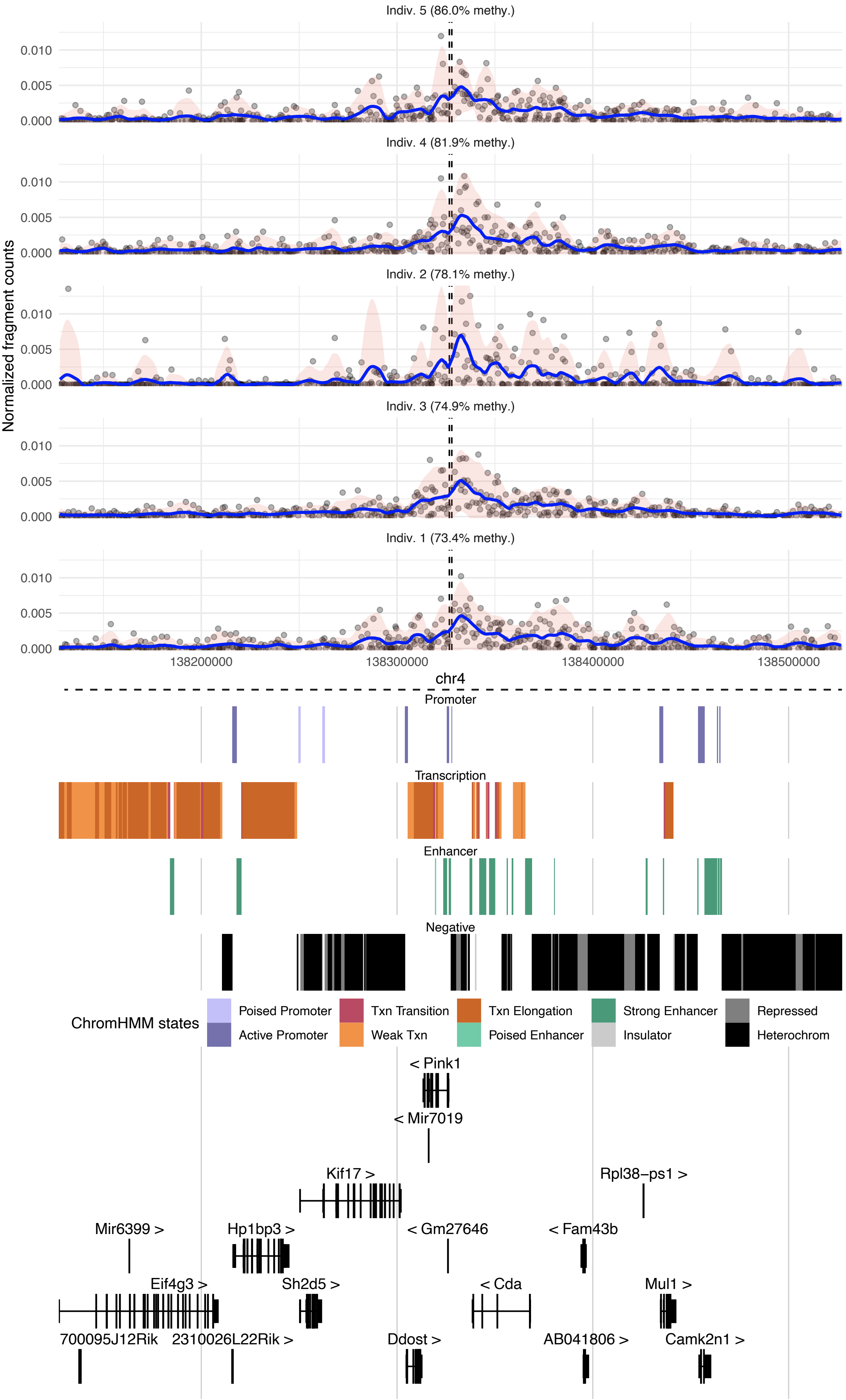

IAP-*Mbnl1* (74.6% - 93.8% methylation range)

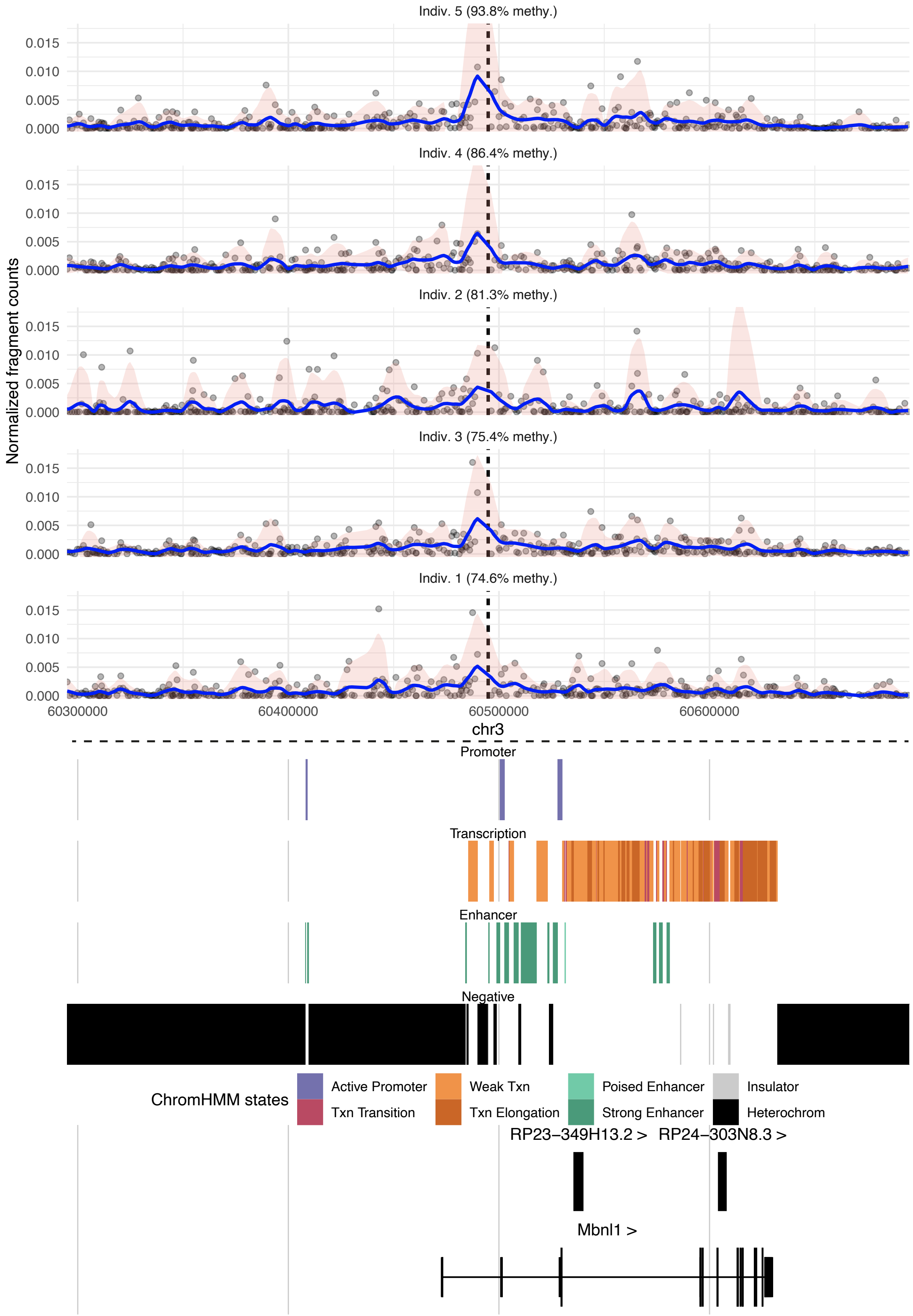

#### IAP-*Dst* (95.4% - 97.1% methylation range)

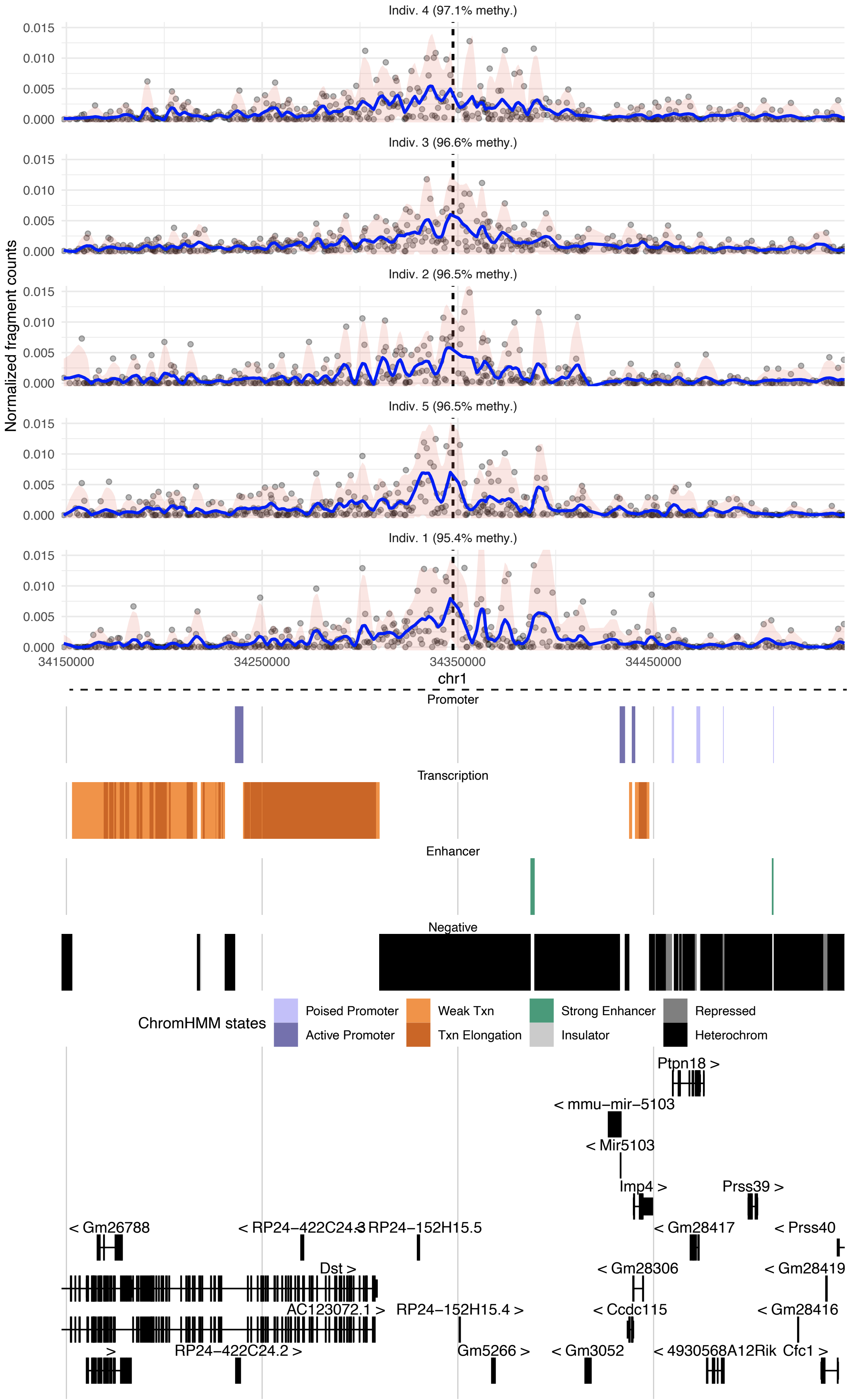

**Supplemental Figure S9. Genomic interactions of VM-IAPs.**

Above the dashed horizontal line are chromatin interaction plots generated from 4C-seq of the following five regions: two cVM-IAPs with low methylation values (IAP-*Marveld2* and IAP-*Tfec*), two cVM-IAPs with high methylation values (IAP-*Pink1* and IAP-*Mbnl1*), and one non-variable highly methylated IAP (IAP-*Dst*). The interaction plots for each region are shown for five individuals and are organised from highest to lowest methylation; each plot shows a 400kb region centred on the 4C viewpoint. The dashed vertical lines at the centre of the plots denote the viewpoint of the 4C-seq experiment and the grey dots represent fragments that interact with this viewpoint – the height of each dot reflects the intensity of the interaction. The trend line (blue) shows the general interaction profile of the region; the pink ribbon delimits the 95% confidence interval of the trend line. Immediately below the dashed horizontal line are the chromHMM chromatin states of the region (largely derived from histone tail modifications) followed by gene annotations.

#### Supplemental Figure S10

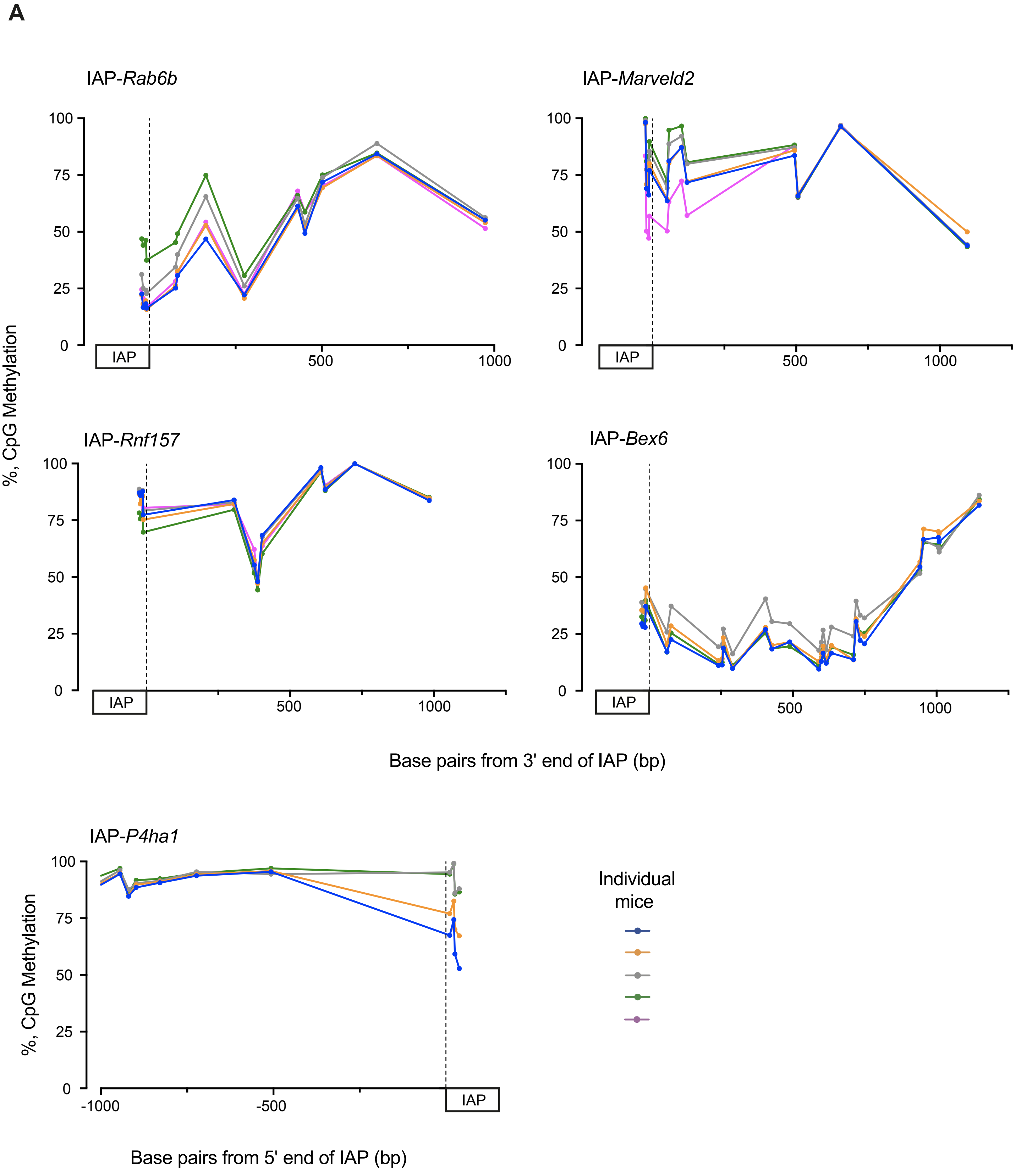

B

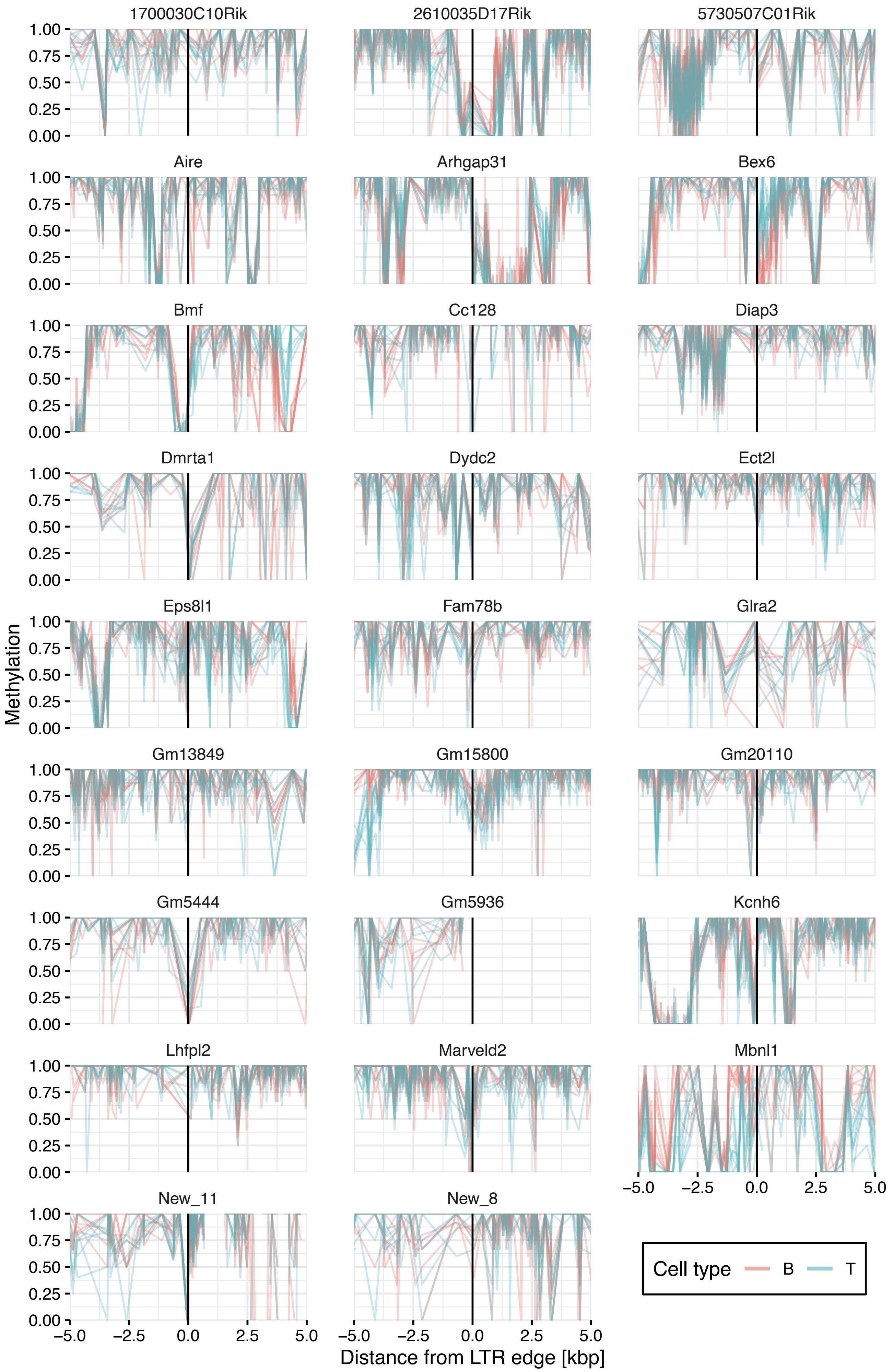

1700030C10Rik
2610035D17Rik
5730507C01Rik
Aire
Arhgap31
Bex6
Bmf
Cc128
Diap3
Dmrta1
Dydc2
Ect2l
Eps8l1
Fam78b
Glra2
Gm13849
Gm15800
Gm20110
Gm5444
Gm5936
Kcnh6
Lhfpl2
Marveld2
Mbnl1
New_11
New_8
Methylation
1.00
0.75
0.50
0.25
0.00
−5.0
−2.5
0.0
2.5
5.0
Distance from LTR edge [kbp]
Cell type
B
T

**B (continued)**

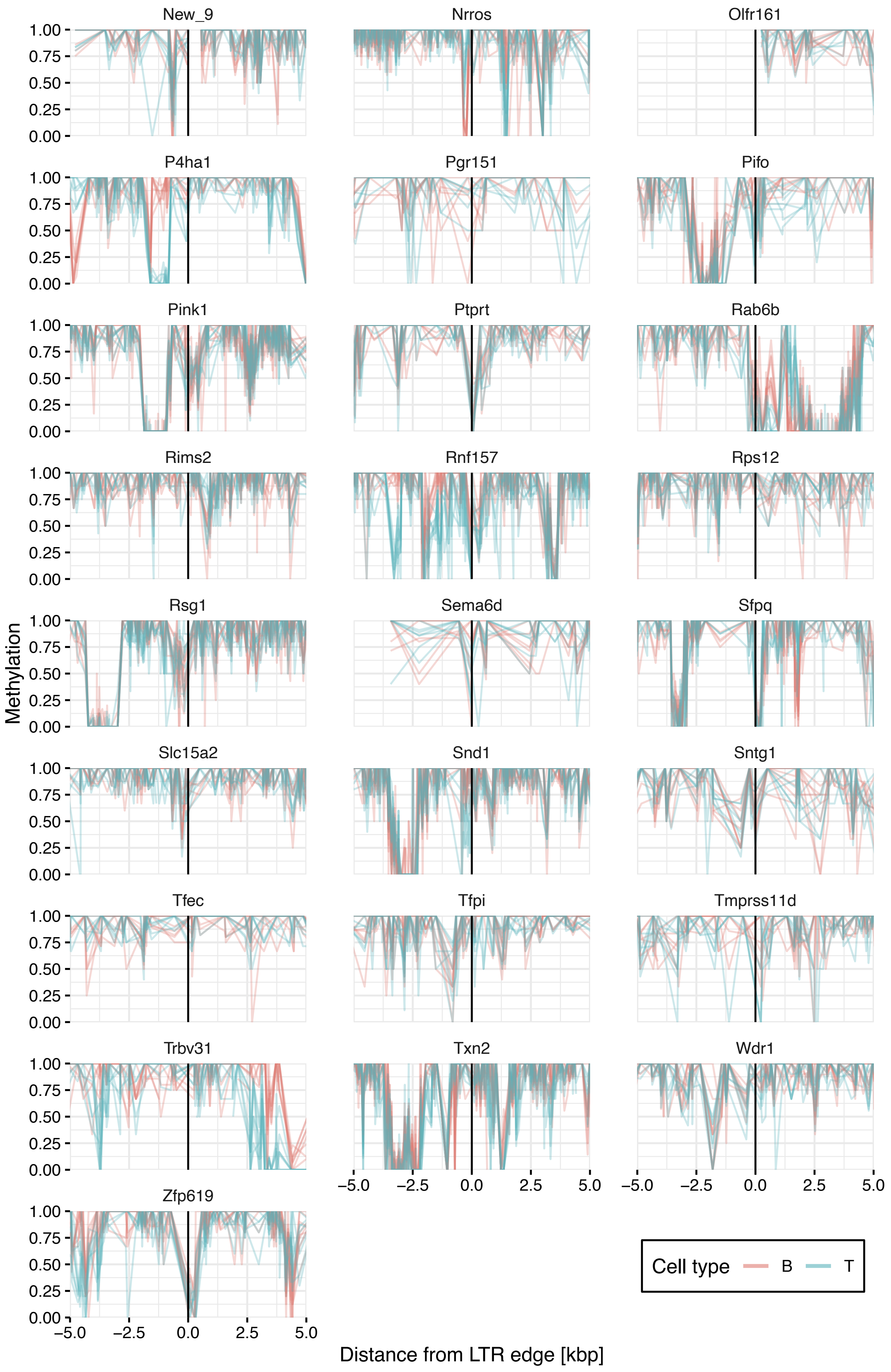

New_9
Nrros
Olfr161
P4ha1
Pgr151
Pifo
Pink1
Ptprt
Rab6b
Rims2
Rnf157
Rps12
Rsg1
Sema6d
Sfpq
Slc15a2
Snd1
Sntg1
Tfec
Tfpi
Tmprss11d
Trbv31
Txn2
Wdr1
Zfp619
Methylation
1.00
0.75
0.50
0.25
0.00
−5.0
−2.5
0.0
2.5
5.0
Distance from LTR edge [kbp]
Cell type
B
T

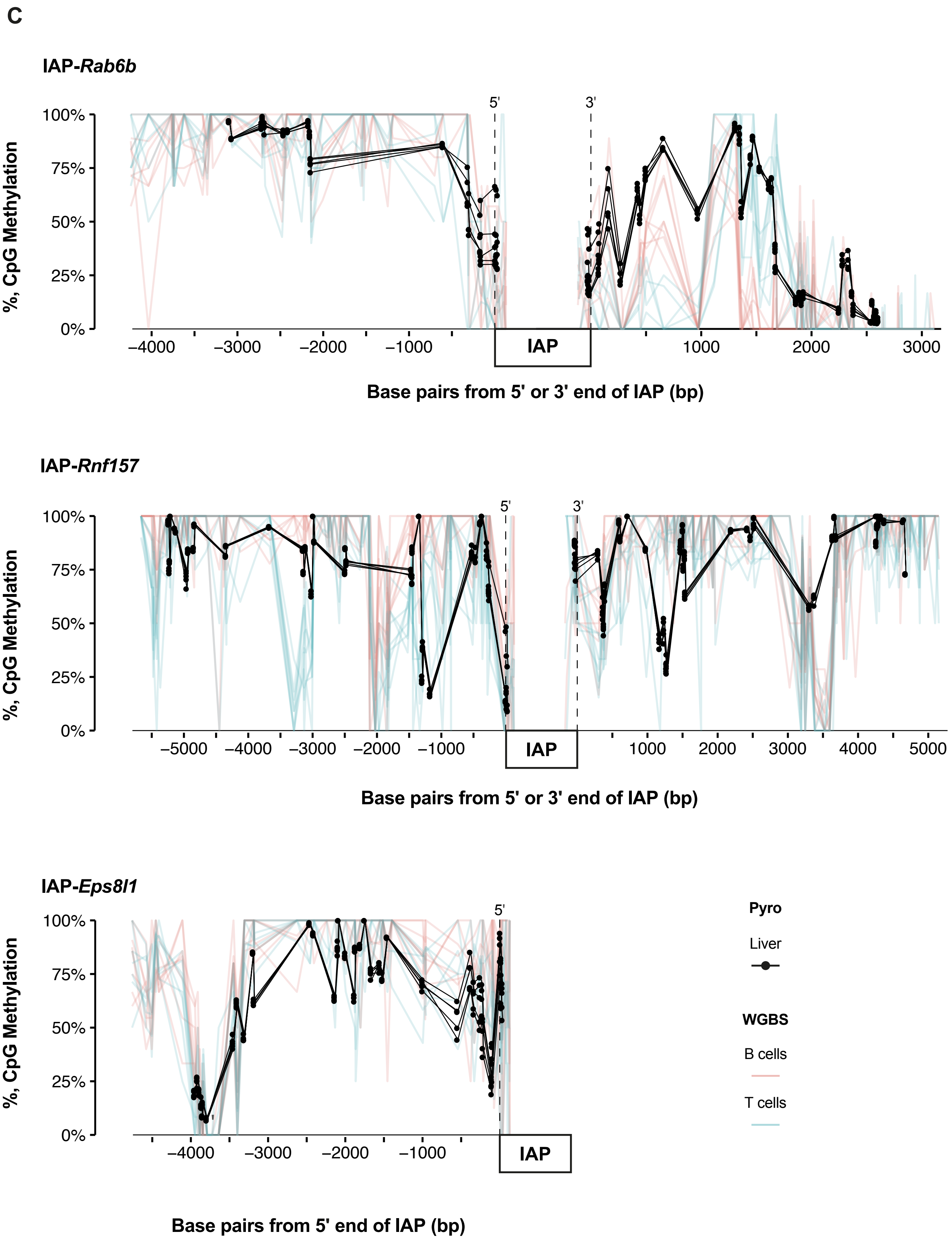
C
IAP-Rab6b
100%
75%
50%
25%
0%
%, CpG Methylation
5'
3'
IAP
−4000
−3000
−2000
−1000
1000
2000
3000
Base pairs from 5' or 3' end of IAP (bp)
IAP-Rnf157
5'
3'
IAP
−5000
−4000
−3000
−2000
−1000
1000
2000
3000
4000
5000
Base pairs from 5' or 3' end of IAP (bp)
IAP-Eps8l1
5'
IAP
−4000
−3000
−2000
−1000
Base pairs from 5' end of IAP (bp)
Pyro
Liver
WGBS
B cells
T cells

C (continued)

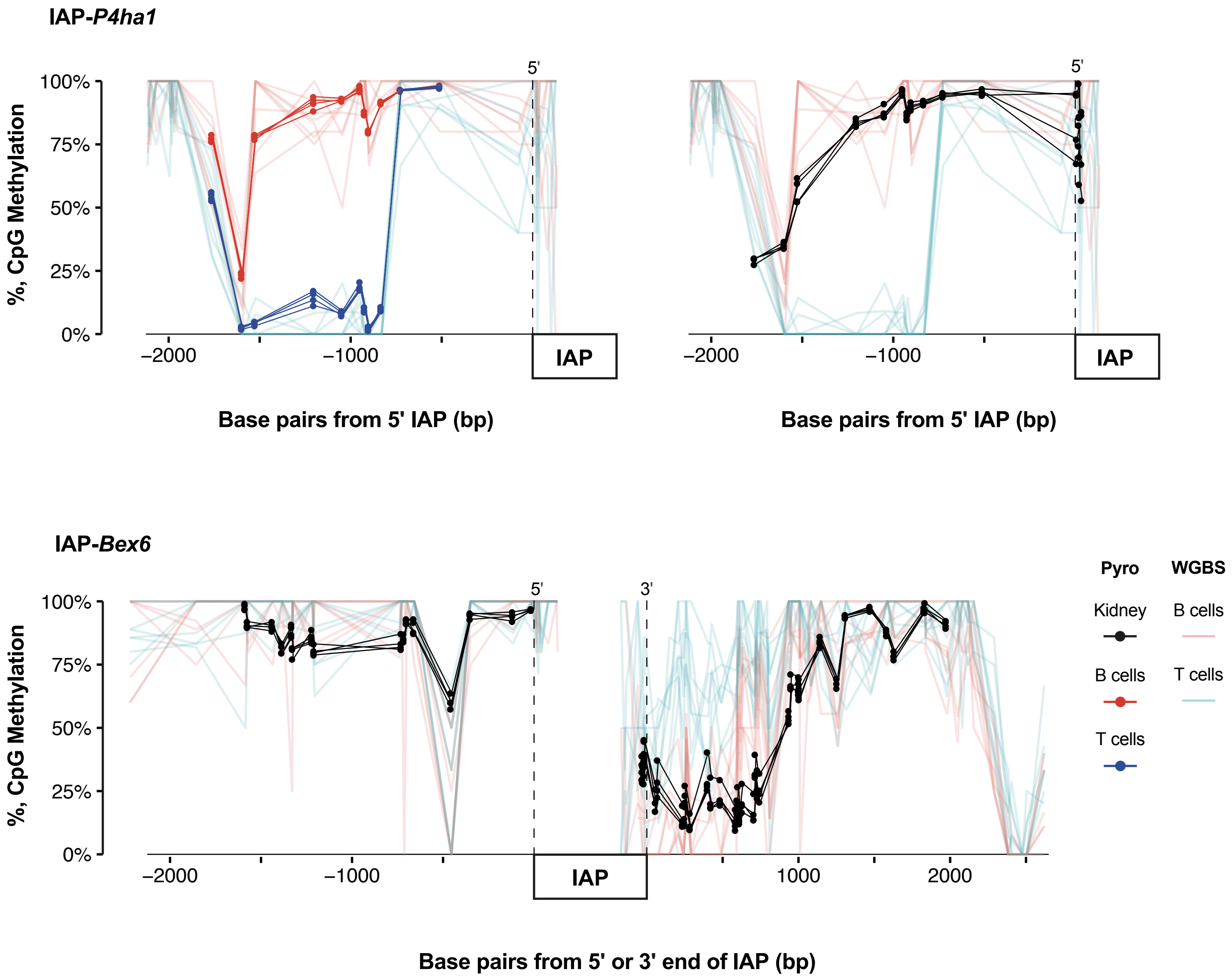

IAP-*P4ha1*
%, CpG Methylation
100%
75%
50%
25%
0%
5'
−2000
−1000
IAP
Base pairs from 5' IAP (bp)
IAP-*Bex6*
3'
1000
2000
Base pairs from 5' or 3' end of IAP (bp)
Pyro
WGBS
Kidney
B cells
T cells

**Supplemental Figure S10. Methylation variability exists beyond the edges of VM-IAPs, but the surrounding methylation landscape varies with each element.**

(A) Bisulfite pyrosequencing of liver (5 individuals; IAP-*Rab6b,* IAP-*Marveld2,* and IAP-*Rnf157*) and kidney (4 individuals; IAP-*Bex6* and IAP-*P4ha1*) shows inter-individual methylation variation within 500-1000bp beyond the edge of the cVM-IAP LTRs.

(B) WGBS data from B and T cells (light red and blue) in the 10kb window surrounding the 5′ and 3′ ends of all validated cVM-IAPs. Methylation within the IAP (black line) is not shown. The y-axis shows DNA methylation from 0 to 1.

(C) WGBS of B and T cells (light red and blue) validated by bisulfite pyrosequencing (Pyro) in liver (black, n=5 for all; IAP-*Rab6b*, IAP-*Rnf157*, IAP-*Eps8l1*), kidney (black, n=4 for both; IAP-*P4ha1*, IAP-*Bex6*), or B and T cells (red and blue, n=4; IAP-*P4ha1*) extracted from individual mice at CpGs several kilobases beyond the 5′ and 3′ edges of cVM-IAPs. IAP length is not to scale. For the bisulfite pyrosequencing data, each dot represents a CpG. The dashed vertical lines represent the 5′ or 3′ ends of the IAPs.
